## Supplemental Material 1. Annotations_duplications_resolved_by_metaannotator for "Preprints in motion: tracking changes between preprint posting and journal publication during a pandemic"

line_break_and_label	doi	line_break_version	n_versions	line_break_and_fiducial	published_pubmed_abstract	page_breaks1
 Preprint DOI:	10.1101/19005710
1

Existing methods to infer the relative roles of age groups in epidemic transmission can normally only accommodate a few age classes, and/or require data that are highly specific for the disease being studied. Here, symbolic transfer entropy (STE), a measure developed to identify asymmetric transfer of information between stochastic processes, is presented as a way to reveal asymmetric transmission patterns between age groups in an epidemic. STE provides a ranking of which age groups may dominate transmission, rather than a reconstruction of the explicit between-age-group transmission matrix. Using simulations, we establish that STE can identify which age groups dominate transmission even when there are differences in reporting rates between age groups and even if the data are noisy. Then, the pairwise STE is calculated between time series of influenza-like illness for 12 age groups in 884 US cities during the autumn of 2009. Elevated STE from 5 to 19 year-olds indicates that school-aged children were likely the most important transmitters of infection during the autumn wave of the 2009 pandemic in the USA. The results may be partially confounded by higher rates of physician-seeking behaviour in children compared to adults, but it is unlikely that differences in reporting rates can explain the observed differences in STE.
2
 Preprint DOI:	10.1101/19006031
2

"Background.: Cognitive impairment associated with lifetime major depressive disorder (MDD) is well-supported by meta-analytic studies, but population-based estimates remain scarce. .Previous UK Biobank studies have only shown limited evidence of cognitive differences related to probable MDD. Using updated cognitive and clinical assessments in UK Biobank, this study investigated population-level differences in cognitive functioning associated with lifetime MDD. Methods.: Associations between lifetime MDD and cognition (performance on six tasks and general cognitive functioning [g-factor]) were investigated in UK Biobank (N-range 7,457-14,836, age 45-81 years, 52% female), adjusting for demographics, education, and lifestyle. Lifetime MDD classifications were based on the Composite International Diagnostic Interview. Within the lifetime MDD group, we additionally investigated relationships between cognition and (a) recurrence, (b) current symptoms, (c) severity of psychosocial impairment (while symptomatic), and (d) concurrent psychotropic medication use. Results.: Lifetime MDD was robustly associated with a lower g-factor (ß = -0.10, PFDR = 4.7 × 10-5), with impairments in attention, processing speed, and executive functioning (ß = 0.06). Clinical characteristics revealed differential profiles of cognitive impairment among case individuals; those who reported severe psychosocial impairment and use of psychotropic medication performed worse on cognitive tests. Severe psychosocial impairment and reasoning showed the strongest association (ß = -0.18, PFDR = 7.5 × 10-5). Conclusions.: Findings describe small but robust associations between lifetime MDD and lower cognitive performance within a population-based sample. Overall effects were of modest effect size, suggesting limited clinical relevance. However, deficits within specific cognitive domains were more pronounced in relation to clinical characteristics, particularly severe psychosocial impairment."
3
 Preprint DOI:	10.1101/19006171
2

Both genetic and environmental factors influence the etiology of age-related macular degeneration (AMD), a leading cause of blindness. AMD severity is primarily measured by images of the fundus of the retina and recently developed machine learning methods can successfully predict AMD progression using image data. However, none of these methods have used both genetic and image data for predicting AMD progression. Here we used both genotypes and fundus images to predict whether an eye had progressed to late AMD with a modified deep convolutional neural network. In total, we used 31,262 fundus images and 52 AMD-associated genetic variants from 1,351 subjects from the Age-Related Eye Disease Study, which provided disease severity phenotypes and fundus images available at baseline and follow-up visits over a period of 12 years. Our results showed that fundus images coupled with genotypes could predict late AMD progression with an averaged area-under-the-curve value of 0.85 (95% confidence interval 0.83–0.86). The results using fundus images alone showed an averaged area under the receiver operating characteristic curve value of 0.81 (95% confidence interval 0.80–0.83). We implemented our model in a cloud-based application for individual risk assessment.
4
 Preprint DOI:	10.1101/19006478
2

Multifocal (MF)/multicentric (MC) ) breast cancer is generally considered to be where two or more breast tumours are present within the same breast, and is seen in ~10% of breast cancer cases. This study investigates the prevalence of multifocality/multicentricity in a cohort of BRCA1/2 mutation carriers with breast cancer from Northern Ireland via cross‐sectional analysis. Data from 211 women with BRCA1/2 mutations (BRCA1 ‐91, BRCA2 ‐120) and breast cancer were collected including age, tumour focality, size, type, grade and receptor profile. The prevalence of multifocality/multicentricity within this group was 25% but, within subgroups, prevalence amongst BRCA2 carriers was more than double that of BRCA1 carriers (p = 0.001). Women affected by MF/MC tumours had proportionately higher oestrogen receptor positivity (p = 0.001) and lower triple negativity (p = 0.004). These observations are likely to be driven by the higher BRCA2 mutation prevalence observed within this cohort. The odds of a BRCA2 carrier developing MF/MC cancer were almost four‐fold higher than a BRCA1 carrier (odds ratio: 3.71, CI: 1.77–7.78, p = 0.001). These findings were subsequently validated in a second, large independent cohort of patients with BRCA ‐associated breast cancers from a UK‐wide multicentre study. This confirmed a significantly higher prevalence of MF/MC tumours amongst BRCA2 mutation carriers compared with BRCA1 mutation carriers.. This has important implications for clinicians involved in the treatment of BRCA2 ‐associated breast cancer, both in the diagnostic process, in ensuring that tumour focality is adequately assessed to facilitate treatment decision‐making, and for breast surgeons, particularly if breast conserving surgery is being considered as a treatment option for these patients.
5
 Preprint DOI:	10.1101/19006502
1

Gambiense human African trypanosomiasis (gHAT) is one of several neglected tropical diseases that is targeted for elimination by the World Health Organization. Recent years have seen a substantial decline in the number of globally reported cases, largely driven by an intensive process of screening and treatment. However, this infection is highly focal, continuing to persist at low prevalence even in small populations. Regional elimination, and ultimately global eradication, rests on understanding the dynamics and persistence of this infection at the local population scale. Here we develop a stochastic model of gHAT dynamics, which is underpinned by screening and reporting data from one of the highest gHAT incidence regions, Kwilu Province, in the Democratic Republic of Congo. We use this model to explore the persistence of gHAT in villages of different population sizes and subject to different patterns of screening. Our models demonstrate that infection is expected to persist for long periods even in relatively small isolated populations. We further use the model to assess the risk of recrudescence following local elimination and consider how failing to detect cases during active screening events informs the probability of elimination. These quantitative results provide insights for public health policy in the region, particularly highlighting the difficulties in achieving and measuring the 2030 elimination goal.
6
 Preprint DOI:	10.1101/19006577
1

"Background There is a wealth of literature on the observed association between childhood trauma and psychotic illness. However, the relationship between childhood trauma and psychosis is complex and could be explained, in part, by gene–environment correlation. Methods The association between schizophrenia polygenic scores (PGS) and experiencing childhood trauma was investigated using data from the Avon Longitudinal Study of Parents and Children (ALSPAC) and the Norwegian Mother, Father and Child Cohort Study (MoBa). Schizophrenia PGS were derived in each cohort for children, mothers, and fathers where genetic data were available. Measures of trauma exposure were derived based on data collected throughout childhood and adolescence (0–17 years; ALSPAC) and at age 8 years (MoBa). Results Within ALSPAC, we found a positive association between schizophrenia PGS and exposure to trauma across childhood and adolescence; effect sizes were consistent for both child or maternal PGS. We found evidence of an association between the schizophrenia PGS and the majority of trauma subtypes investigated, with the exception of bullying. These results were comparable with those of MoBa. Within ALSPAC, genetic liability to a range of additional psychiatric traits was also associated with a greater trauma exposure. Conclusions Results from two international birth cohorts indicate that genetic liability for a range of psychiatric traits is associated with experiencing childhood trauma. Genome-wide association study of psychiatric phenotypes may also reflect risk factors for these phenotypes. Our findings also suggest that youth at higher genetic risk might require greater resources/support to ensure they grow-up in a healthy environment."
7
 Preprint DOI:	10.1101/19006635
2

"PurposeSpinal muscular atrophy (SMA), caused by loss of the SMN1 gene, is a leading cause of early childhood death. Due to the near identical sequences of SMN1 and SMN2, analysis of this region is challenging. Population-wide SMA screening to quantify the SMN1 copy number (CN) is recommended by the American College of Medical Genetics and Genomics.MethodsWe developed a method that accurately identifies the CN of SMN1 and SMN2 using genome sequencing (GS) data by analyzing read depth and eight informative reference genome differences between SMN1/2.ResultsWe characterized SMN1/2 in 12,747 genomes, identified 1568 samples with SMN1 gains or losses and 6615 samples with SMN2 gains or losses, and calculated a pan-ethnic carrier frequency of 2%, consistent with previous studies. Additionally, 99.8% of our SMN1 and 99.7% of SMN2 CN calls agreed with orthogonal methods, with a recall of 100% for SMA and 97.8% for carriers, and a precision of 100% for both SMA and carriers.carrierscarrierscarrierscarriers.ConclusionThis SMN copy-number caller can be used to identify both carrier and affected status of SMA, enabling SMA testing to be offered as a comprehensive test in neonatal care and an accurate carrier screening tool in GS sequencing projects."
8
 Preprint DOI:	10.1101/19006650
1

Patients with major depressive disorder (MDD) show heterogeneous treatment response and highly variable clinical trajectories: while some patients experience swift recovery, others show relapsing-remitting or chronic courses. Predicting individual clinical trajectories at an early stage is a key challenge for psychiatry and might facilitate individually tailored interventions. So far, however, reliable predictors at the single-patient level are absent. Here, we evaluated the utility of a machine learning strategy – generative embedding (GE) – which combines interpretable generative models with discriminative classifiers. Specifically, we used functional magnetic resonance imaging (fMRI) data of emotional face perception in 85 MDD patients from the NEtherlands Study of Depression and Anxiety (NESDA) who had been followed up over two years and classified into three subgroups with distinct clinical trajectories. Combining a generative model of effective (directed) connectivity with support vector machines (SVMs), we could predict whether a given patient would experience chronic depression vs. fast remission with a balanced accuracy of 79%. Gradual improvement vs. fast remission could still be predicted above-chance, but less convincingly, with a balanced accuracy of 61%. Generative embedding outperformed classification based on conventional (descriptive) features, such as functional connectivity or local activation estimates, which were obtained from the same data and did not allow for above-chance classification accuracy. Furthermore, predictive performance of GE could be assigned to a specific network property: the trial-by-trial modulation of connections by emotional content. Given the limited sample size of our study, the present results are preliminary but may serve as proof-of-concept, illustrating the potential of GE for obtaining clinical predictions that are interpretable in terms of network mechanisms. Our findings suggest that abnormal dynamic changes of connections involved in emotional face processing might be associated with higher risk of developing a less favorable clinical course.	.
9
 Preprint DOI:	10.1101/19006833

1
 Higher proportions of early-onset and estrogen receptor (ER) negative cancers are observed in women of African ancestry than in women of European ancestry. Differences in risk factor distributions and associations by age at diagnosis and ER status may explain this disparity. We analyzed data from 1,126 cases (aged 18–74<U+2009>years) with invasive breast cancer and 2,106 controls recruited from a population-based case–control study in Ghana. Odds ratios (OR) and 95% confidence intervals (CI) were estimated for menstrual and reproductive factors using polytomous logistic regression models adjusted for potential confounders. Among controls, medians for age at menarche, parity, age at first birth, and breastfeeding/pregnancy were 15<U+2009>years, 4 births, 20<U+2009>years and 18<U+2009>months, respectively. For women =50<U+2009>years, parity and extended breastfeeding were associated with decreased risks: >5 births vs . nulliparous, OR 0.40 (95% CI 0.20–0.83) and 0.71 (95% CI 0.51–0.98) for =19 vs . <13 breastfeeding months/pregnancy, which did not differ by ER. In contrast, for earlier onset cases (<50<U+2009>years) parity was associated with increased risk for ER-negative tumors (p -heterogeneity by ER = 0.02), which was offset by extended breastfeeding. Similar associations were observed by intrinsic-like subtypes. Less consistent relationships were observed with ages at menarche and first birth. Reproductive risk factor distributions are different from European populations but exhibited etiologic heterogeneity by age at diagnosis and ER status similar to other populations. Differences in reproductive patterns and subtype heterogeneity are consistent with racial disparities in subtype distributions....
10
 Preprint DOI:	10.1101/19006957
2

"Background The English Indices of Multiple Deprivation (IMD) is widely used as a measure of deprivation. However, similarly ranked areas can differ substantially in the underlying domains of deprivation. These domains contain a richer set of data that might be useful for classifying local authorities. Clustering methods offer a set of techniques to identify groups of areas with similar patterns of deprivation. Methods Hierarchical agglomerative (i.e. bottom-up) clustering methods were applied to domain scores for 152 upper tier local authorities. Advances in statistical testing allow clusters to be identified that are unlikely to have arisen from random partitioning of a homogeneous group. The resulting clusters are described in terms of their subdomain scores and basic geographic and demographic characteristics. Results Five statistically significant clusters of local authorities were identified. These clusters only partially reflect different levels of overall deprivation. . In particular, two clusters share similar overall IMD scores but have contrasting patterns of deprivation. . Conclusion Hierarchical clustering methods identify five distinct clusters that do not correspond closely to quintiles of deprivation. This approach may help to distinguish between places that face similar underlying challenges, and places that appear similar in terms of overall deprivation scores, but that face different challenges."
11
 Preprint DOI:	10.1101/19007013
2

"Background Previous literature has demonstrated a strong association between cigarette smoking, suicidal ideation and suicide attempts. This association has not previously been examined in a causal inference framework and could have important implications for suicide prevention strategies. Aims We aimed to examine the evidence for an association between smoking behaviours (initiation, smoking status, heaviness, lifetime smoking) and suicidal thoughts or attempts by triangulating across observational and Mendelian randomisation analyses. Method First, in the UK Biobank, we calculated observed associations between smoking behaviours and suicidal thoughts or attempts. Second, we used Mendelian randomisation to explore the relationship between smoking and suicide attempts and ideation, using genetic variants as instruments to reduce bias from residual confounding and reverse causation. Results Our observational analysis showed a relationship between smoking behaviour, suicidal ideation and attempts, particularly between smoking initiation and suicide attempts (odds ratio, 2.07; 95% CI 1.91–2.26; P < 0.001). The Mendelian randomisation analysis and single-nucleotide polymorphism analysis, however, did not support this (odds ratio for lifetime smoking on suicidal ideation, 0.050; 95% CI -0.027 to 0.127; odds ratio on suicide attempts, 0.053; 95% CI, -0.003 to 0.110). ). Despite past literature showing a positive dose-response relationship, our results showed no clear evidence for a causal effect of smoking on suicidal ideation or attempts. Conclusions This was the first Mendelian randomisation study to explore the effect of smoking on suicidal ideation and attempts. .Our results suggest that, despite observed associations, there is no clear evidence for a causal effect."."."."
12
 Preprint DOI:	10.1101/19007021
1

"BackgroundThe triage of patients in prehospital care is a difficult task, and improved risk assessment tools are needed both at the dispatch center and on the ambulance to differentiate between low- and high-risk patients. This study validates a machine learning-based approach to generating risk scores based on hospital outcomes using routinely collected prehospital data.MethodsDispatch, ambulance, and hospital data were collected in one Swedish region from 2016–2017. Dispatch center and ambulance records were used to develop gradient boosting models predicting hospital admission, critical care (defined as admission to an intensive care unit or in-hospital mortality), and two-day mortality. Composite risk scores were generated based on the models and compared to National Early Warning Scores (NEWS) and actual dispatched priorities in a prospectively gathered dataset from 2018.ResultsA total of 38203 patients were included from 2016–2018. Concordance indexes (or areas under the receiver operating characteristics curve) for dispatched priorities ranged from 0.51–0.66, while those for NEWS ranged from 0.66–0.85. Concordance ranged from 0.70–0.79 for risk scores based only on dispatch data, and 0.79–0.89 for risk scores including ambulance data. Dispatch data-based risk scores consistently outperformed dispatched priorities in predicting hospital outcomes, while models including ambulance data also consistently outperformed NEWS. Model performance in the prospective test dataset was similar to that found using cross-validation, and calibration was comparable to that of NEWS.ConclusionsMachineConclusionsMachineConclusionsMachineConclusionsMachineConclusionsMachineConclusionsMachineConclusionsMachineConclusionsMachine learning-based risk scores outperformed a widely-used rule-based triage algorithm and human prioritization decisions in predicting hospital outcomes. Performance was robust in a prospectively gathered dataset, and scores demonstrated adequate calibration. Future research should explore the robustness of these methods when applied to other settings, establish appropriate outcome measures for use in determining the need for prehospital care, and investigate the clinical impact of interventions based on these methods."
13
 Preprint DOI:	10.1101/19007161
1

Treatment-resistant depression (TRD) occurs in ~30% of patients with major depressive disorder (MDD) but the genetics of TRD was previously poorly investigated. Whole exome sequencing and genome-wide genotyping were available in 1209 MDD patients after quality control. Antidepressant response was compared to non-response to one treatment and non-response to two or more treatments (TRD). Differences in the risk of carrying damaging variants were tested. A score expressing the burden of variants in genes and pathways was calculated weighting each variant for its functional (Eigen) score and frequency. Gene-based and pathway-based scores were used to develop predictive models of TRD and non-response using gradient boosting in 70% of the sample (training) which were tested in the remaining 30% (testing), evaluating also the addition of clinical predictors. Independent replication was tested in STAR*D and GENDEP using exome array-based data. TRD and non-responders did not show higher risk to carry damaging variants compared to responders. Genes/pathways associated with TRD included those modulating cell survival and proliferation, neurodegeneration, and immune response. Genetic models showed significant prediction of TRD vs. response and they were improved by the addition of clinical predictors, but they were not significantly better than clinical predictors alone. Replication results were driven by clinical factors, except for a model developed in subjects treated with serotonergic antidepressants, which showed a clear improvement in prediction at the extremes of the genetic score distribution in STAR*D. These results suggested relevant biological mechanisms implicated in TRD and a new methodological approach to the prediction of TRD.
14
 Preprint DOI:	10.1101/19007401
1

"BackgroundDiabetes mellitus (DM) is a global health care problem that can impose a substantial economic burden. Diabetic peripheral neuropathy (DPN) is a common microvascular complication of DM that increases the potential for morbidity and disability due to ulceration and amputation. . Though there is a significant amount of variation in the primary studies on DM regarding the prevalence of DPN in Africa. Hence, this study was aimed to estimate the overall prevalence of DPN in DM patients in Africa.MethodsPubMed, Scopus, Google Scholar, African Journals OnLine, WHO African Library, and the Cochrane Review were systematically searched online to retrieve related articles. The Preferred Reporting Items for Systematic Review and Meta-Analysis (PRISMA) guidelines was followed. Heterogeneity across the included studies was evaluated by the inconsistency index (I2). Publication bias was examined by funnel plot and Egger’s regression test. The random-effect model was fitted to estimate the pooled prevalence of diabetic peripheral neuropathy among patients in Africa. The meta-analysis was performed using the STATA™ Version 14 software.ResultsTwenty-three studies which includes 269,691 participants were included in the meta-analysis. The overall pooled prevalence of diabetic peripheral neuropathy was 46% (95% CI:36.21–55.78%). Based on the subgroup analysis, the highest prevalence of diabetic peripheral neuropathy in DM patients was reported in West Africa at 49.4% (95% CI: 32.74, 66.06).ConclusionThis study revealed that the overall prevalence of diabetic peripheral neuropathy is relatively high in Africa. Hence, DPN needs situation-based interventions and preventive strategies, which are specific to the country. Further meta-analysis is needed to identify associated factors for the occurrence of diabetic peripheral neuropathy."
15
 Preprint DOI:	10.1101/19007450
1

"Background: Epstein–Barr virus (EBV) infection is thought to play a central role in the development of multiple sclerosis (MS). If causal, it represents a target for interventions to reduce MS risk. Objective: To examine the evidence for interaction between EBV and other risk factors, and explore mechanisms via which EBV infection may influence MS risk. Methods: Pubmed was searched using the terms ‘multiple sclerosis’ AND ‘Epstein Barr virus’, ‘multiple sclerosis’ AND EBV, ‘clinically isolated syndrome’ AND ‘Epstein Barr virus’ and ‘clinically isolated syndrome’ AND EBV. All abstracts were reviewed for possible inclusion. Results: A total of 262 full-text papers were reviewed. There was evidence of interaction on the additive scale between anti-EBV antibody titre and HLA genotype (attributable proportion due to interaction ((((AP)<U+2009>=<U+2009>0.48, p<U+2009><<U+2009>1<U+2009>×<U+2009>10-4)).)..).. Previous infectious mononucleosis (IM) was associated with increased odds ratio (OR) of MS in HLA-DRB1*1501 positive but not HLA-DRB1*1501 negative persons. Smoking was associated with a greater risk of MS in those with high anti-EBV antibodies (OR<U+2009>=<U+2009>2.76) but not low anti-EBV antibodies (OR<U+2009>=<U+2009>1.16). No interaction between EBV and risk factors was found on a multiplicative scale. Conclusion: EBV appears to interact with at least some established MS risk factors. The mechanism via which EBV influences MS risk remains unknown."
16
 Preprint DOI:	10.1101/19007542
2

"Background: Neuroimaging studies have just begun to explore the acute effects of psychedelics on large-scale brain networks' functional organization. Even less is known about the neural correlates of subacute effects taking place days after the psychedelic experience. This study explores the subacute changes of primary sensory brain networks and networks supporting higher-order affective and self-referential functions 24 hours after a single session with the psychedelic ayahuasca. Methods: We leveraged task-free functional magnetic resonance imaging data 1 day before and 1 day after a randomized placebo-controlled trial exploring the effects of ayahuasca in naïve healthy participants (21 placebo/22 ayahuasca). We derived intra- and inter-network functional connectivity of the salience, default mode, visual, and sensorimotor networks, and assessed post-session connectivity changes between the ayahuasca and placebo groups. Connectivity changes were associated with Hallucinogen Rating Scale scores assessed during the acute effects. Results: Our findings revealed increased anterior cingulate cortex connectivity within the salience network, decreased posterior cingulate cortex connectivity within the default mode network, and increased connectivity between the salience and default mode networks 1 day after the session in the ayahuasca group compared to placebo. Connectivity of primary sensory networks did not differ between groups. Salience network connectivity increases correlated with altered somesthesia scores, decreased default mode network connectivity correlated with altered volition scores, and increased salience default mode network connectivity correlated with altered affect scores. . Conclusion: These findings provide preliminary evidence for subacute functional changes induced by the psychedelic ayahuasca on higher-order cognitive brain networks that support interoceptive, affective, and self-referential functions."
17
 Preprint DOI:	10.1101/19007666
2

This large, retrospective case-control study of electronic health records from 56 million unique adult patients examined whether or not treatment with a Tumor Necrosis Factor (TNF) blocking agent is associated with lower risk for Alzheimer’s disease (AD) in patients with rheumatoid arthritis (RA), psoriasis, and other inflammatory diseases which are mediated in part by TNF and for which a TNF blocker is an approved treatment. The analysis compared the diagnosis of AD as an outcome measure in patients receiving at least one prescription for a TNF blocking agent (etanercept, adalimumab, and infliximab) or for methotrexate. Adjusted odds ratios (AORs) were estimated using the Cochran-Mantel-Haenszel (CMH) method and presented with 95% confidence intervals (CIs) and p-values. . RA was associated with a higher risk for AD (Adjusted Odds Ratio (AOR) = 2.06, 95% Confidence Interval: (2.02–2.10), P-value <0.0001) as did psoriasis (AOR = 1.37 (1.31–1.42), P <0.0001), ankylosing spondylitis (AOR = 1.57 (1.39–1.77), P <0.0001), inflammatory bowel disease (AOR = 2.46 (2.33–2.59), P < 0.0001), ulcerative colitis (AOR = 1.82 (1.74–1.91), P <0.0001), and Crohn’s disease (AOR = 2.33 (2.22–2.43), P <0.0001). The risk for AD in patients with RA was lower among patients treated with etanercept (AOR = 0.34 (0.25–0.47), P <0.0001), adalimumab (AOR = 0.28 (0.19–0.39), P < 0.0001), or infliximab (AOR = 0.52 (0.39–0.69), P <0.0001). Methotrexate was also associated with a lower risk for AD (AOR = 0.64 (0.61–0.68), P <0.0001), while lower risk was found in patients with a prescription history for both a TNF blocker and methotrexate. Etanercept and adalimumab also were associated with lower risk for AD in patients with psoriasis: AOR = 0.47 (0.30–0.73 and 0.41 (0.20–0.76), respectively. There was no effect of gender or race, while younger patients showed greater benefit from a TNF blocker than did older patients. This study identifies a subset of patients in whom systemic inflammation contributes to risk for AD through a pathological mechanism involving TNF and who therefore may benefit from treatment with a TNF blocking agent.
18
 Preprint DOI:	10.1101/19007773
1

"Objective To determine whether naturally occurring autoantibodies against the prion protein are present in individuals with genetic prion disease mutations and controls, and if so, whether they are protective against prion disease.Methods In this case–control study, we collected 124 blood samples from individuals with a variety of pathogenic PRNP mutations and 78 control individuals with a positive family history of genetic prion disease but lacking disease-associated PRNP mutations. Antibody reactivity was measured using an indirect ELISA for the detection of human immunoglobulin G1–4 antibodies against wild-type human prion protein. Multivariate linear regression models were constructed to analyze differences in autoantibody reactivity between (1) PRNP mutation carriers vs controls and (2) asymptomatic vs symptomatic PRNP mutation carriers. Robustness of results was examined in matched cohorts.Results We found that antibody reactivity was present in a subset of both PRNP mutation carriers and controls. Autoantibody levels were not influenced by PRNP mutation status or clinical manifestation of prion disease. Post hoc analyses showed anti-PrPC autoantibody titers to be independent of personal history of autoimmune disease and other immunologic disorders, as well as PRNP codon 129 polymorphism.Conclusions Pathogenic PRNP variants do not notably stimulate antibody-mediated anti-PrPC immunity. Anti-PrPC immunoglobulin G autoantibodies are not associated with the onset of prion disease. The presence of anti-PrPC autoantibodies in the general population without any disease-specific association suggests that relatively high titers of naturally occurring antibodies are well-tolerated."
19
 Preprint DOI:	10.1101/19007849
1

"Introduction Patients with type 2 diabetes mellitus (T2DM) are at risk for a variety of severe debilitating effects. One of the most serious complications experienced by patients with T2DM are skeletal diseases caused by changes in the bone microenvironment. As a result, patients with T2DM are at risk for higher prevalence of fragility fractures.... There are a variety of treatments available for counteracting this effect. Some antidiabetic medications, such as metformin, have been shown to have a positive effect on bone health without the addition of additional drugs into patients’ treatment plans. Chinese randomised controlled trial (RCT) studies have also proposed antiosteoporotic pharmacotherapies as a viable alternative treatment strategy. .Previous network meta-analyses (NMAs) and meta-analyses regarding this topic did not include all available RCT trials, or only performed pairwise comparisons. We present a protocol for a two-part NMA that incorporates all available RCT data to provide the most comprehensive ranking of antidiabetics (part I) and antiosteoporotic (part II) pharmacotherapies in terms of their ability to decrease fracture incidences, increase bone mineral density (BMD) and improve indications of bone turnover markers (BTMs) in adult patients with T2DM.Methods and analysis We will search Medical Literature Analysis and Retrieval System Online, Excerpta Medica Database, PubMed, Web of Science, Cumulative Index to Nursing and Allied Health Literature, Cochrane Central Register of Controlled Trials and Chinese literature sources (China National Knowledge Infrastructure, Chongqing VIP Information, Wanfang Data, Wanfang Med Online) for RCTs, which fit our criteria. We will include adult patients with T2DM who have taken antidiabetics (part I) or antiosteoporotic (part II) therapies with relevant outcome measures in our study. We will perform title/abstract and full-text screening as well as data extraction in duplicate. Risk of bias will be evaluated in duplicate for each study, and the quality of evidence will be examined using Confidence in Network Meta-Analysis in accordance to the Grading of Recommendations Assessment, Development and Evaluation framework. We will use R and gemtc to perform the NMA. We will report changes in BMD and BTMs in either weighted or standardised mean difference, and we will report fracture incidences as ORs. We will use the Surface Under the Cumulative Ranking Curve scores to provide numerical estimates of the rankings of interventions.Ethics and dissemination The study will not require ethics approval. The findings of the two-part NMA will be disseminated in peer-reviewed journals and presented at conferences. We aim to produce the most comprehensive quantitative analysis regarding the management of T2DM bone disease. Our analysis should be able to provide physicians and patients with up-to-date recommendations for antidiabetic medications and antiosteoporotic pharmacotherapies for maintaining bone health in patients with T2DM."
21
 Preprint DOI:	10.1101/19008102
1

"BackgroundDisability-Adjusted Life Years (DALYs) are an established method for quantifying population health needs and guiding prioritisation decisions. Global Burden of Disease (GBD) estimates aim to ensure comparability between countries and over time by using age-standardised rates (ASR) to account for differences in the age structure of different populations. Different standard populations are used for this purpose but it is not widely appreciated that the choice of standard may affect not only the resulting rates but also the rankings of causes of DALYs. We aimed to evaluate the impact of the choice of standard, using the example of Scotland.MethodsDALY estimates were derived from the 2016 Scottish Burden of Disease (SBoD) study for an abridged list of 68 causes of disease/injury, representing a three-year annual average across 2014–16. Crude DALY rates were calculated using Scottish national population estimates. DALY ASRs standardised using the GBD World Standard Population (GBD WSP) were compared to those using the 2013 European Standard Population (ESP2013). Differences in ASR and in rank order within the cause list were summarised for all-cause and for each individual cause.ResultsThe ranking of causes by DALYs were similar using crude rates or ASR (ESP2013). All-cause DALY rates using ASR (GBD WSP) were around 26% lower. Overall 58 out of 68 causes had a lower ASR using GBD WSP compared with ESP2013, with the largest falls occurring for leading causes of mortality observed in older ages. Gains in ASR were much smaller in absolute scale and largely affected causes that operated early in life. These differences were associated with a substantial change to the ranking of causes when GBD WSP was used compared with ESP2013.ConclusionDisease rankings based on DALY ASRs are strongly influenced by the choice of standard population. While GBD WSP offers international comparability, within-country analyses based on DALY ASRs should reflect local age structures. For European countries, including Scotland, ESP2013 may better guide local priority setting by avoiding large disparities occurring between crude and age-standardised results sets, which could potentially confuse non-technical audiences."."	."	."	."
22
 Preprint DOI:	10.1101/19008136
1

"
"Objectives: Deep brain stimulation (DBS) of the centromedian thalamic nucleus (CM) is an emerging treatment for multiple brain diseases, including the drug-resistant epilepsy Lennox-Gastaut syndrome (LGS). We aimed to improve neurosurgical targeting of the CM by: (1) developing a structural MRI approach for CM visualisation, (2) identifying the CM's neurophysiological characteristics using microelectrode recordings (MERs) and (3) mapping connectivity from CM-DBS sites using functional MRI (fMRI). Methods: 19 patients with LGS (mean age=28 years) underwent presurgical 3T MRI using magnetisation-prepared 2 rapid acquisition gradient-echoes (MP2RAGE) and fMRI sequences; 16 patients proceeded to bilateral CM-DBS implantation and intraoperative thalamic MERs. CM visualisation was achieved by highlighting intrathalamic borders on MP2RAGE using Sobel edge detection. Mixed-effects analysis compared two MER features (spike firing rate and background noise) between ventrolateral, CM and parafasicular nuclei. Resting-state fMRI connectivity was assessed using implanted CM-DBS electrode positions as regions of interest. Results: The CM appeared as a hyperintense region bordering the comparatively hypointense pulvinar, mediodorsal and parafasicular nuclei. At the group level, reduced spike firing and background noise distinguished CM from the ventrolateral nucleus; however, these trends were not found in 20%-25% of individual MER trajectories. Areas of fMRI connectivity included basal ganglia, brainstem, cerebellum, sensorimotor/premotor and limbic cortex. Conclusions: In the largest clinical trial of DBS undertaken in patients with LGS to date, we show that accurate targeting of the CM is achievable using 3T MP2RAGE MRI. Intraoperative MERs may provide additional localising features in some cases; however, their utility is limited by interpatient variability. Therapeutic effects of CM-DBS may be mediated via connectivity with brain networks that support diverse arousal, cognitive and sensorimotor processes."
23
 Preprint DOI:	10.1101/19008227
1

"Aims Point-of-care (POC) tests for influenza and respiratory syncytial virus (RSV) offer the potential to improve patient management and antimicrobial stewardship. Studies have focused on performance; however, no workflow assessments have been published comparing POC molecular tests. This study compared the Liat and ID Now systems workflow, to assist end-users in selecting an influenza and/or RSV POC test.Methods Staffing, walk-away and turnaround time (TAT) of the Liat and ID Now systems were determined using 40 nasopharyngeal samples, positive for influenza or RSV. The ID Now system requires separate tests for influenza and RSV, so parallel (two instruments) and sequential (one instrument) workflows were evaluated.Results The ID Now ranged 4.1–6.2 min for staffing, 1.9–10.9 min for walk-away and 6.4–15.8 min for TAT per result. The Liat ranged 1.1–1.8 min for staffing, 20.0–20.5 min for walk-away and 21.3–22.0 min for TAT. Mean walk-away time comprised 38.0% (influenza positive) and 68.1% (influenza negative) of TAT for ID Now and 93.7% (influenza/RSV) for Liat. The ID Now parallel workflow resulted in medians of 5.9 min for staffing, 9.7 min for walk-away and 15.6 min for TAT. Assuming prevalence of 20% influenza and 20% RSV, the ID Now sequential workflow resulted in medians of 9.4 min for staffing, 17.4 min for walk-away, and 27.1 min for TAT.Conclusions The ID Now and Liat systems offer different workflow characteristics. Key considerations for implementation include value of both influenza and RSV results, clinical setting, staffing capacity, and instrument(s) placement."
24
 Preprint DOI:	10.1101/19008276
1

Hyperosmolar solutions are widely used to treat raised intracranial pressure following severe traumatic brain injury. Although mannitol has historically been the most frequently administered, hypertonic saline solutions are increasingly being used. However, definitive evidence regarding their comparative effectiveness is lacking. The Sugar or Salt Trial is a UK randomised, allocation concealed open label multicentre pragmatic trial designed to determine the clinical and cost-effectiveness of hypertonic saline compared with mannitol in the management of patients with severe traumatic brain injury. Patients requiring intensive care unit admission and intracranial pressure monitoring post-traumatic brain injury will be allocated at random to receive equi-osmolar boluses of either mannitol or hypertonic saline following failure of routine first-line measures to control intracranial pressure. The primary outcome for the study will be the Extended Glasgow Outcome Scale assessed at six months after randomisation. Results will inform current clinical practice in the routine use of hyperosmolar therapy as well as assess the impact of potential side effects. Pre-planned longer term clinical and cost effectiveness analyses will further inform the use of these treatments.
25
 Preprint DOI:	10.1101/19008300
1

"Purpose: Neurodevelopmental disorders represent a frequent indication for clinical exome sequencing. Fifty percent of cases, however, remain undiagnosed even upon exome reanalysis. Here we show RNA sequencing (RNA-seq) on human B-lymphoblastoid cell lines (LCL) is highly suitable for neurodevelopmental Mendelian gene testing and demonstrate the utility of this approach in suspected cases of Cornelia de Lange syndrome (CdLS). Methods: Genotype-Tissue Expression project transcriptome data for LCL, blood, and brain were assessed for neurodevelopmental Mendelian gene expression. Detection of abnormal splicing and pathogenic variants in these genes was performed with a novel RNA-seq diagnostic pipeline and using a validation CdLS-LCL cohort (n = 10) and test cohort of patients who carry a clinical diagnosis of CdLS but negative genetic testing (n = 5). ). ). Results: LCLs share isoform diversity of brain tissue for a large subset of neurodevelopmental genes and express 1.8-fold more of these genes compared with blood (LCL, n = 1706; whole blood, n = 917). This enables testing of more than 1000 genetic syndromes. . . The RNA-seq pipeline had 90% sensitivity for detecting pathogenic events and revealed novel diagnoses such as abnormal splice products in NIPBL and pathogenic coding variants in BRD4 and ANKRD11. . Conclusion: The LCL transcriptome enables robust frontline and/or reflexive diagnostic testing for neurodevelopmental disorders."
26
 Preprint DOI:	10.1101/19008342
1

In a case–control study of patients with Clostridium difficile infection, we found no statistically significant association between the presence of trehalose utilization variants in infecting C. difficile strains and development of severe infection outcome. These results do not support trehalose utilization conferring enhanced virulence in the context of human C. difficile infections.
27
 Preprint DOI:	10.1101/19008367
1

"Background.: Current approaches for early identification of individuals at high risk for autism spectrum disorder (ASD) in the general population are limited, and most ASD patients are not identified until after the age of 4. This is despite substantial evidence suggesting that early diagnosis and intervention improves developmental course and outcome. The aim of the current study was to test the ability of machine learning (ML) models applied to electronic medical records (EMRs) to predict ASD early in life, in a general population sample. Methods.: We used EMR data from a single Israeli Health Maintenance Organization, including EMR information for parents of 1,397 ASD children (ICD-9/10) and 94,741 non-ASD children born between January 1st, 1997 and December 31st, 2008. Routinely available parental sociodemographic information, parental medical histories, and prescribed medications data were used to generate features to train various ML algorithms, including multivariate logistic regression, artificial neural networks, and random forest. Prediction performance was evaluated with 10-fold cross-validation by computing the area under the receiver operating characteristic curve (AUC; C-statistic), sensitivity, specificity, accuracy, false positive rate, and precision (positive predictive value [PPV]). Results.: All ML models tested had similar performance. The average performance across all models had C-statistic of 0.709, sensitivity of 29.93%, specificity of 98.18%, accuracy of 95.62%, false positive rate of 1.81%, and PPV of 43.35% for predicting ASD in this dataset. Conclusions.: We conclude that ML algorithms combined with EMR capture early life ASD risk as well as reveal previously unknown features to be associated with ASD-risk. .Such approaches may be able to enhance the ability for accurate and efficient early detection of ASD in large populations of children."
28
 Preprint DOI:	10.1101/19008615
1

Aetiological and clinical heterogeneity is increasingly recognized as a common characteristic of Alzheimer’s disease and related dementias. This heterogeneity complicates diagnosis, treatment, and the design and testing of new drugs. An important line of research is discovery of multimodal biomarkers that will facilitate the targeting of subpopulations with homogeneous pathophysiological signatures. High-throughput ‘omics’ are unbiased data-driven techniques that probe the complex aetiology of Alzheimer’s disease from multiple levels (e.g. network, cellular, and molecular) and thereby account for pathophysiological heterogeneity in clinical populations. This review focuses on data reduction analyses that identify complementary disease-relevant perturbations for three omics techniques: neuroimaging-based subtypes, metabolomics-derived metabolite panels, and genomics-related polygenic risk scores. Neuroimaging can track accrued neurodegeneration and other sources of network impairments, metabolomics provides a global small-molecule snapshot that is sensitive to ongoing pathological processes, and genomics characterizes relatively invariant genetic risk factors representing key pathways associated with Alzheimer’s disease. Following this focused review, we present a roadmap for assembling these multiomics measurements into a diagnostic tool highly predictive of individual clinical trajectories, to further the goal of personalized medicine in Alzheimer’s disease.
29
 Preprint DOI:	10.1101/19008953
1

"BackgroundExisting physical activity guidelines predominantly focus on healthy age-stratified target groups. The objective of this study was to develop evidence-based recommendations for physical activity (PA) and PA promotion for German adults (18–65 years) with noncommunicable diseases (NCDs).MethodsThe PA recommendations were developed based on existing PA recommendations. In phase 1, systematic literature searches were conducted for current PA recommendations for seven chronic conditions (osteoarthrosis of the hip and knee, chronic obstructive pulmonary disease, stable ischemic heart disease, stroke, clinical depression, and chronic non-specific back pain). In phase 2, the PA recommendations were evaluated on the basis of 28 quality criteria, and high-quality recommendations were analysed. In phase 3, PA recommendations for seven chronic conditions were deducted and then synthesised to generate generic German PA recommendations for adults with NCDs. In relation to the recommendations for PA promotion, a systematic literature review was conducted on papers that reviewed the efficacy/effectiveness of interventions for PA promotion in adults with NCDs.ResultsThe German recommendations for physical activity state that adults with NCDs should, over the course of a week, do at least 150 min of moderate-intensity aerobic PA, or 75 min of vigorous-intensity aerobic PA, or a combination of both. Furthermore, muscle-strengthening activities should be performed at least twice a week. The promotion of PA among adults with NCDs should be theory-based, specifically target PA behaviour, and be tailored to the respective target group. In this context, and as an intervention method, exercise referral schemes are one of the more promising methods of promoting PA in adults with NCDs.ConclusionThe development of evidence-based recommendations for PA and PA promotion is an important step in terms of the initiation and implementation of actions for PA-related health promotion in Germany. The German recommendations for PA and PA promotion inform adults affected by NCDs and health professionals on how much PA would be optimal for adults with NCDs. Additionally, the recommendations provide professionals entrusted in PA promotion the best strategies and interventions to raise low PA levels in adults with NCDs. The formulation of specific PA recommendations for adults with NCDs and their combination with recommendations on PA promotion is a unique characteristic of the German recommendations."
30
 Preprint DOI:	10.1101/19009472

1
 "Introduction:Constraint-induced movement therapy (CIMT) improves upper limb (UL) motor execution in unilateral cerebral palsy (uCP). As these children also show motor planning deficits, action-observation training (AOT) might be of additional value. Here, we investigated the combined effect of AOT to CIMT and identified factors influencing treatment response.Methods:A total of 44 children with uCP (mean 9 years 6 months, SD 1 year 10 months) participated in a 9-day camp wearing a splint for 6 h/day and were allocated to the CIMT + AOT (n = 22) and the CIMT + placebo group (n = 22). The CIMT + AOT group received 15 h of AOT (i.e. video-observation) and executed the observed tasks, whilst the CIMT + AOT group watched videos free of biological motion and executed the same tasks. The primary outcome measure was bimanual performance. Secondary outcomes included measures of body function and activity level assessed before (T1), after the intervention (T2), and at 6 months follow-up (T3). Influencing factors included behavioural and neurological characteristics.Results:Although no between-groups differences were found (p > 0.05; η2 = 0–16), ), the addition of AOT led to higher gains in children with initially poorer bimanual performance (p = 0.02; η2 = 0.14). ). Both groups improved in all outcome measures after the intervention and retained the gains at follow up (p < 0.01; η2 = 0.02–0.71)).).).. Poor sensory function resulted in larger improvements in the total group (p = 0.03; η2 = 0.25) and high amounts of mirror movements tended to result in a better response to the additional AOT training (p = 0.06; η2 = 0.18). ). Improvements were similar irrespective of the type of brain lesion or corticospinal tract wiring pattern.Conclusions:Adding AOT to CIMT, resulted in a better outcome for children with poor motor function and high amounts of mirror movements. CIMT with or without AOT seems to be more beneficial for children with poor sensory function."
31
 Preprint DOI:	10.1101/19009522
1

Anal squamous cell carcinoma is a rare tumor. Chemo-radiotherapy yields a 50% 3-year relapse-free survival rate in advanced anal cancer, so improved predictive markers and therapeutic options are needed. High-throughput proteomics and whole-exome sequencing were performed in 46 paraffin samples from anal squamous cell carcinoma patients. Hierarchical clustering was used to establish groups de novo. Then, probabilistic graphical models were used to study the differences between groups of patients at the biological process level. A molecular classification into two groups of patients was established, one group with increased expression of proteins related to adhesion, T lymphocytes and glycolysis; and the other group with increased expression of proteins related to translation and ribosomes. The functional analysis by the probabilistic graphical model showed that these two groups presented differences in metabolism, mitochondria, translation, splicing and adhesion processes. Additionally, these groups showed different frequencies of genetic variants in some genes, such as ATM, SLFN11, and DST. Finally, genetic and proteomic characteristics of these groups suggested the use of some possible targeted therapies, such as PARP inhibitors or immunotherapy
32
 Preprint DOI:	10.1101/19009589
1

"
"The spread of dengue through global human mobility is a major public health concern. A key challenge is understanding the transmission pathways and mediating factors that characterized the patterns of dengue importation into non-endemic areas. Utilizing a network connectivity-based approach, we analyze the importation patterns of dengue fever into European countries. Seven connectivity indices were developed to characterize the role of the air passenger traffic, seasonality, incidence rate, geographical proximity, epidemic vulnerability, and wealth of a source country, in facilitating the transport and importation of dengue fever. We used generalized linear mixed models (GLMMs) to examine the relationship between dengue importation and the connectivity indices while accounting for the air transport network structure. We also incorporated network autocorrelation within a GLMM framework to investigate the propensity of a European country to receive an imported case, by virtue of its position within the air transport network. The connectivity indices and dynamical processes of the air transport network were strong predictors of dengue importation in Europe. With more than 70% of the variation in dengue importation patterns explained. We found that transportation potential was higher for source countries with seasonal dengue activity, high passenger traffic, high incidence rates, high epidemic vulnerability, and in geographical proximity to a destination country in Europe. We also found that position of a European country within the air transport network was a strong predictor of the country’s propensity to receive an imported case. Our findings provide evidence that the importation patterns of dengue into Europe can be largely explained by appropriately characterizing the heterogeneities of the source, and topology of the air transport network. This contributes to the foundational framework for building integrated predictive models for bio-surveillance of dengue importation."
33
 Preprint DOI:	10.1101/19009647
1

"Background and ObjectiveUnderstanding pharmacokinetic disposition of cefepime, a β-lactam antibiotic, is crucial for developing regimens to achieve optimal exposure and improved clinical outcomes. This study sought to develop and evaluate a unified population pharmacokinetic model in both pediatric and adult patients receiving cefepime treatment.MethodsMultiple physiologically relevant models were fit to pediatric and adult subject data. To evaluate the final model performance, a withheld group of 12 pediatric patients and two separate adult populations were assessed.ResultsSeventy subjects with a total of 604 cefepime concentrations were included in this study. All adults (n = 34) on average weighed 82.7 kg and displayed a mean creatinine clearance of 106.7 mL/min. All pediatric subjects (n = 36) had mean weight and creatinine clearance of 16.0 kg and 195.6 mL/min, respectively. A covariate-adjusted two-compartment model described the observed concentrations well (population model R2, 87.0%; Bayesian model R2, 96.5%). In the evaluation subsets, the model performed similarly well (population R2, 84.0%; Bayesian R2, 90.2%).ConclusionThe identified model serves well for population dosing and as a Bayesian prior for precision dosing."
34
 Preprint DOI:	10.1101/19010322
1

"ObjectivesWe examine long-term retention of adults, adolescents and children on antiretroviral therapy under different HIV treatment guidelines in Malawi.DesignProspective cohort study.Setting and participantsAdults and children starting ART between 2005 and 2015 in 21 health facilities in southern Malawi.MethodsWe used survival analysis to assess retention at clinic level, Cox regression to examine risk factors for loss to follow up, and competing risk analysis to assess long-term outcomes of people on antiretroviral therapy (ART).ResultsWe included 132,274 individuals in our analysis, totalling 270,256 person years of follow up (PYFU; median per patient 1.3, interquartile range (IQR) 0.26–3.1), 62% were female and the median age was 32 years. Retention on ART was lower in the first year on ART compared to subsequent years for all guideline periods and age groups. Infants (0–3 years), adolescents and young adults (15–24 years) were at highest risk of LTFU. Comparing the different calendar periods of ART initiation we found that retention improved initially, but remained stable thereafter.ConclusionEven though the number of patients and the burden on health care system increased substantially during the study period of rapid ART expansion, retention on ART improved in the early years of ART provision, but gains in retention were not maintained over 5 years on ART. Reducing high attrition in the first year of ART should remain a priority for ART programs, and so should addressing poor retention among adolescents, young adults and men."
35
 Preprint DOI:	10.1101/19010462
3

"Background: Acute and chronic low back pain (LBP) are different conditions with different treatments. However, they are coded in electronic health records with the same International Classification of Diseases, 10th revision (ICD-10) code (M54.5) and can be differentiated only by retrospective chart reviews. This prevents an efficient definition of data-driven guidelines for billing and therapy recommendations, such as return-to-work options.Objective: The objective of this study was to evaluate the feasibility of automatically distinguishing acute LBP episodes by analyzing free-text clinical notes.Methods: We used a dataset of 17,409 clinical notes from different primary care practices; of these, 891 documents were manually annotated as acute LBP and 2973 were generally associated with LBP via the recorded ICD-10 code. We compared different supervised and unsupervised strategies for automated identification: keyword search, topic modeling, logistic regression with bag of n-grams and manual features, and deep learning (a convolutional neural network-based architecture [ConvNet]). We trained the supervised models using either manual annotations or ICD-10 codes as positive labels.Results: ConvNet trained using manual annotations obtained the best results with an area under the receiver operating characteristic curve of 0.98 and an F score of 0.70. . ConvNet’s results were also robust to reduction of the number of manually annotated documents. In the absence of manual annotations, topic models performed better than methods trained using ICD-10 codes, which were unsatisfactory for identifying LBP acuity.Conclusions: This study uses clinical notes to delineate a potential path toward systematic learning of therapeutic strategies, billing guidelines, and management options for acute LBP at the point of care."
36
 Preprint DOI:	10.1101/19010579
1

"In this paper, we present an overview and descriptive results from one of the first egocentric network studies of men who have sex with men (MSM) from across the United States: the ARTnet study. ARTnet was designed to support prevention research for human immunodeficiency virus (HIV) and other sexually transmitted infections (STIs) that are transmitted across partnership networks. ARTnet implemented a population-based egocentric network study design that sampled egos from the target population and asked them to report on the number, attributes, and timing of their sexual partnerships. Such data provide the foundation needed for parameterizing stochastic network models that are used for disease projection and intervention planning. ARTnet collected data online from 2017 to 2019, with a final sample of 4904 participants who reported on 16198 sexual partnerships. The aims of this paper were to characterize the joint distribution of three network parameters needed for modeling: degree distributions, assortative mixing, and partnership age, with heterogeneity by partnership type (main, casual and one-time), demography, and geography. Participants had an average of 1.19 currently active partnerships (“mean degree”), which was higher for casual partnerships (0.74) than main partnerships (0.45). The mean rate of one-time partnership acquisition was 0.16 per week (8.5 partners per year). Main partnerships lasted 272.5 weeks on average, while casual partnerships lasted 133.0 weeks. There was strong but heterogenous assortative mixing by race/ethnicity for all groups. The mean absolute age difference for all partnership types was 9.5 years, with main partners differing by 6.3 years compared to 10.8 years for casual partners. Our analysis suggests that MSM may be at sustained risk for HIV/STI acquisition and transmission through high network degree of sexual partnerships. The ARTnet network study provides a robust and reproducible foundation for understanding the dynamics of HIV/STI epidemiology among U.S. MSM and supporting the implementation science that seeks to address persistent challenges in HIV/STI prevention."
37
 Preprint DOI:	10.1101/19010728

1

"BackgroundDNA methylation outlier burden has been suggested as a potential marker of biological age. An outlier is typically defined as DNA methylation levels at any one CpG site that are three times beyond the inter-quartile range from the 25th or 75th percentiles compared to the rest of the population. DNA methylation outlier burden (the number of such outlier sites per individual) increases exponentially with age. However, these findings have been observed in small samples.ResultsHere, we showed an association between age and log10-transformed DNA methylation outlier burden in a large cross-sectional cohort, the Generation Scotland Family Health Study (N = 7010, β = 0.0091, p < 2 × 10−16), and in two longitudinal cohort studies, the Lothian Birth Cohorts of 1921 (N = 430, β = 0.033, p = 7.9 × 10−4) and 1936 (N = 898, β = 0.0079, p = 0.074). Significant confounders of both cross-sectional and longitudinal associations between outlier burden and age included white blood cell proportions, body mass index (BMI), smoking, and batch effects. In Generation Scotland, the increase in epigenetic outlier burden with age was not purely an artefact of an increase in DNA methylation level variability with age (epigenetic drift). Log10-transformed DNA methylation outlier burden in Generation Scotland was not related to self-reported, or family history of, age-related diseases, and it was not heritable (SNP-based heritability of 4.4%, p = 0.18). Finally, DNA methylation outlier burden was not significantly related to survival in either of the Lothian Birth Cohorts individually or in the meta-analysis after correction for multiple testing (HRmeta = 1.12; 95% CImeta = [1.02; 1.21]; pmeta = 0.021).ConclusionsThese findings suggest that, while it does not associate with ageing-related health outcomes, DNA methylation outlier burden does track chronological ageing and may also relate to survival. DNA methylation outlier burden may thus be useful as a marker of biological ageing."
38
 Preprint DOI:	10.1101/19010975
1

State-of-the-art machine learning (ML) artificial intelligence methods are increasingly leveraged in clinical predictive modeling to provide clinical decision support systems to physicians. Modern ML approaches such as artificial neural networks (ANNs) and tree boosting often perform better than more traditional methods like logistic regression. On the other hand, these modern methods yield a limited understanding of the resulting predictions. However, in the medical domain, understanding of applied models is essential, in particular, when informing clinical decision support. Thus, in recent years, interpretability methods for modern ML methods have emerged to potentially allow explainable predictions paired with high performance. To our knowledge, we present in this work the first explainability comparison of two modern ML methods, tree boosting and multilayer perceptrons (MLPs), to traditional logistic regression methods using a stroke outcome prediction paradigm. Here, we used clinical features to predict a dichotomized 90 days post-stroke modified Rankin Scale (mRS) score. For interpretability, we evaluated clinical features’ importance with regard to predictions using deep Taylor decomposition for MLP, Shapley values for tree boosting and model coefficients for logistic regression. With regard to performance as measured by Area under the Curve (AUC) values on the test dataset, all models performed comparably: Logistic regression AUCs were 0.83, 0.83, 0.81 for three different regularization schemes; tree boosting AUC was 0.81; MLP AUC was 0.83. Importantly, the interpretability analysis demonstrated consistent results across models by rating age and stroke severity consecutively amongst the most important predictive features. For less important features, some differences were observed between the methods. Our analysis suggests that modern machine learning methods can provide explainability which is compatible with domain knowledge interpretation and traditional method rankings. Future work should focus on replication of these findings in other datasets and further testing of different explainability methods.
39
 Preprint DOI:	10.1101/19011049

1

""Objective To compare the reproducibility of 11 antibody assays for immunoglobulin (Ig) G and IgM myelin oligodendrocyte glycoprotein antibodies (MOG-IgG and MOG-IgM) from 5 international centers.Methods The following samples were analyzed: MOG-IgG clearly positive sera (n = 39), MOG-IgG low positive sera (n = 39), borderline negative sera (n = 13), clearly negative sera (n = 40), and healthy blood donors (n = 30). As technical controls, 18 replicates (9 MOG-IgG positive and 9 negative) were included. All samples and controls were recoded, aliquoted, and distributed to the 5 testing centers, which performed the following antibody assays: 5 live and 1 fixed immunofluorescence cell-based assays (CBA-IF, 5 MOG-IgG, and 1 MOG-IgM), 3 live flow cytometry cell-based assays (CBA-FACS, all MOG-IgG), and 2 ELISAs (both MOG-IgG).Results We found excellent agreement (96%) between the live CBAs for MOG-IgG for samples previously identified as clearly positive or negative from 4 different national testing centers. The agreement was lower with fixed CBA-IF (90%), and the ELISA showed no concordance with CBAs for detection of human MOG-IgG. All CBAs showed excellent interassay reproducibility. The agreement of MOG-IgG CBAs for borderline negative (77%) and particularly low positive (33%) samples was less good. Finally, most samples from healthy blood donors (97%) were negative for MOG-IgG in all CBAs.Conclusions Live MOG-IgG CBAs showed excellent agreement for high positive and negative samples at 3 international testing centers. Low positive samples were more frequently discordant than in a similar comparison of aquaporin-4 antibody assays. Further research is needed to improve international standardization for clinical care."
40
 Preprint DOI:	10.1101/19011064
1

"Background50% of Malagasy children have moderate to severe stunting. In 2016, a new 10 year National Nutrition Action Plan (PNAN III) was initiated to help address stunting and developmental delay. We report factors associated with risk of developmental delay in 3 and 4 year olds in the rural district of Ifanadiana in southeastern Madagascar in 2016.MethodsThe data are from a cross-sectional analysis of the 2016 wave of IHOPE panel data (a population-representative cohort study begun in 2014). We interviewed women ages 15–49 using the MICS Early Child Development Indicator (ECDI) module, which includes questions for physical, socio-emotional, learning and literacy/numeracy domains. We analyzed ECDI data using standardized z scores for relative relationships for 2 outcomes: at-risk-for-for-delay vs. an international standard, and lower-development-than-peers if ECDI z scores were > 1 standard deviation below study mean. Covariates included demographics, adult involvement, household environment, and selected, and selected child health factors. Variables significant at alpha of 0.1 were included a multivariable model; final models used backward stepwise regression, clustered at the sampling level.ResultsOf 432 children ages 3 and 4 years, 173 (40%) were at risk for delay compared to international norms and 68 children (16.0%) had lower-development than peers. This was driven mostly by the literacy/numeracy domain, with only 7% of children considered developmentally on track in that domain. 50.5% of children had moderate to severe stunting. 76 (17.6%) had > = 4 stimulation activities in past 3 days.Greater paternal engagement (OR 1.5 (1.09, 2.07)))) was associated with increased delay vs. international norms. Adolescent motherhood (OR. 4.09 (1.40, 11.87)) decreased children’s development vs. peers. Engagement from a non-parental adult reduced odds of delay for both outcomes (OR (95%CI = 0.76 (0.63, 0.91) & 0.27 ((((0.15, 0 48) respectively). Stunting was not associated with delay risk (1.36 (0.85, 2.15) or low development (0.92 (0.48, 1.78)) when controlling for other factors.ConclusionsIn this setting of high child malnutrition, stunting is not independently associated with developmental risk. A low proportion of children receive developmentally supportive stimulation from adults, but non-parent adults provide more stimulation in general than either mother or father. Stimulation from non-parent adults is associated with lower odds of delay."
41
 Preprint DOI:	10.1101/19011130

1
 "BackgroundMalnutrition especially undernutrition is the main problem that is seen over people living with HIV/AIDS and can occur at any age. Multiple factors contributed to undernutrition of HIV/AIDS patients and it need immediate identification and prompt action. The objective of this study was to assess the nutritional status of patients and identify factors associated with undernutrition among HIV/AIDS patients on follow-up care in Jimma medical center, Southwest Ethiopia.MethodsA cross-sectional study design was conducted from March-April 2016. Data were collected retrospectively from clinical records of HIV/AIDS patients enrolled for follow up care in ART clinic from June 2010 to January 2016. Bivariate and multivariate logistic regression analysis were performed to identify independent predictor of undernutrition.ResultsData of 1062 patients were included in the study. The prevalence of undernutrition (BMI<18.5 kg/m2) and overweight or obesity were 34% and 9%, respectively. Out of undernourished patients, severely malnourished patients (BMI<16 kg/m2) accounted of 9%. Undernutrition was more likely among widowed patients (AOR = 1.7, 95% CI, 1.03–2.79),),)),,),),)),, patients with no access to water supply (AOR = 1.69, 95% CI, 1.16–2.47) ) and patients in the WHO clinical stage of three (AOR = 2.0, 95% CI, 1.33–2.97) and four (AOR = 3.0, 95% CI, 1.74–5.07).).).). Moreover, the odds of undernutrition was more likely among patients with CD4 cell count of <200 cells/mm3 (AOR = 2.0, 95% CI, 1.38–2.47) and patients with a functional status of bedridden (AOR = 3.6, 95% CI, 1.55–8.35) and ambulatory (AOR = 2.4, 95% CI, 1.66–3.51), respectively.ConclusionBoth undernutrition and overweight or obesity were prevalent among HIV/AIDS patients in Jimma Medical Center, Ethiopia. Undernutrition was significantly associated with clinical outcome of patients. Hence, nutritional assessment, care and support should be strengthened. Critical identification of malnourished patients and prompt interventions should be undertaken."
42
 Preprint DOI:	10.1101/19011213
1

"BackgroundConcern"BackgroundConcern has been raised about consequences of including patients with left ventricular assist device (LVAD) or heart transplantation in readmission and mortality measures.MethodsWe calculated unadjusted and hospital-specific 30-day risk-standardized mortality (RSMR) and readmission (RSRR) rates for all Medicare fee-for-service beneficiaries with a primary diagnosis of AMI or HF discharged between July 2010 and June 2013. Hospitals were compared before and after excluding LVAD and heart transplantation patients. LVAD indication was measured.ResultsIn the AMI mortality (n = 506,543) ) and readmission (n = 526,309) ) cohorts, 1,166 and 1,016 patients received an LVAD while 3 and 2 had a heart transplantation, respectively. In the HF mortality (n = 1,015,335) and readmission (n = 1,254,124) cohorts, 789 and 931 received an LVAD, while 212 and 202 received a heart transplantation, respectively. Less than 2% of hospitals had either ≥6 patients who received an LVAD or, independently, had ≥1 heart transplantation. The AMI mortality and readmission cohorts used 1.8% and 2.8% of LVADs for semi-permanent/permanent indications, versus 73.8% and 78.0% for HF patients, respectively. The rest were for temporary/external indications. In the AMI cohort, RSMR for hospitals without LVAD patients versus hospitals with ≥6 LVADs was 14.8% and 14.3%, and RSRR was 17.8% and 18.3%, respectively; the HF cohort RSMR was 11.9% and 9.7% and RSRR was 22.6% and 23.4%, respectively. In the AMI cohort, RSMR for hospitals without versus with heart transplantation patients was 14.7% and 13.9% and RSRR was 17.8% and 17.7%, respectively; in the HF cohort, RSMR was 11.9% and 11.0%, and RSRR was 22.6% and 22.6%, respectively. Estimations changed ≤0.1% after excluding LVAD or heart transplantation patients.ConclusionHospitals caring for ≥6 patients with LVAD or ≥1 heart transplantation typically had a trend toward lower RSMRs but higher RSRRs. Rates were insignificantly changed when these patients were excluded. LVADs were primarily for acute-care in the AMI cohort and chronic support in the HF cohort. LVAD and heart transplantation patients are a distinct group with differential care requirements and outcomes, thus should be considered separately from the rest of the HF cohort."
43
 Preprint DOI:	10.1101/19011452

1
 Many California nail salon workers are low-income Vietnamese women of reproductive age who use nail products daily that contain androgen-disrupting phthalates, which may increase risk of male reproductive tract abnormalities during pregnancy. Yet, few studies have characterized phthalate exposures among this workforce. To characterize individual metabolites and cumulative phthalates exposure among a potentially vulnerable occupational group of nail salon workers, we collected 17 post-shift urine samples from Vietnamese workers at six San Francisco Bay Area nail salons in 2011, which were analyzed for four primary phthalate metabolites: mono-n-butyl-, mono-isobutyl-, mono(2-Ethylhexyl)-, and monoethyl phthalates (MnBP, MiBP, MEHP, and MEP, respectively; μg/L). Phthalate metabolite concentrations and a potency-weighted sum of parent compound daily intake (Σandrogen-disruptor, μg/kg/day) were compared to 203 Asian Americans from the 2011–2012 National Health and Nutritional Examination Survey (NHANES) using Student’s t-test and Wilcoxin signed rank test. Creatinine-corrected MnBP, MiBP, MEHP (μg/g), and cumulative phthalates exposure (Σandrogen-disruptor, μg/kg/day) levels were 2.9 (p < 0.0001), 1.6 (p = 0.015), 2.6 (p < 0.0001), and 2.0 (p < 0.0001) times higher, respectively, in our nail salon worker population compared to NHANES Asian Americans. Levels exceeded the NHANES 95th or 75th percentiles among some workers. This pilot study suggests that nail salon workers are disproportionately exposed to multiple phthalates, a finding that warrants further investigation to assess their potential health significance.
44
 Preprint DOI:	10.1101/19011577
1

Leishmaniasis is a vector borne zoonosis which is classified as a neglected tropical disease. Among the three most common forms of the disease, Visceral Leishmaniasis (VL) is the most threatening to human health, causing 20,000 to 30,000 deaths worldwide each year. Areas where VL is mostly endemic have unprotected dogs in community and houses. The "presence of dogs usually increases VL risk for humans since dogs are the principal reservoir host for the parasite of the disease. Based on this fact, most earlier studies consider culling dogs as a control measure for the spread of VL. A more recent control measure has been the use of deltamethrin-impregnated dog collars (s) to protect both humans and dogs by putting s on dogs neck. The presence of dogs helps to grow the sandfly population faster by offering a more suitable blood-meal source. On the other hand, the presence of s on dogs helps to reduce sandfly population by the lethality of deltamethrin insecticide. This study brings an ecological perspective to this public health concern, aiming to understand the impact of an additional host (here, protected dogs) on disease risk to a primary host (here, humans). To answer this question, we compare two different settings: a community without dogs, and a community with dogs protected with . Our analysis shows the presence of protected dogs can reduce VL infection risk in humans. However, this disease risk reduction depends on dogs’ tolerance for sandfly bites.
45
 Preprint DOI:	10.1101/19011775
1

To determine the feasibility of complex home-based phenotyping, 1,876 research participants from the customer base of 23andMe completed an online version of a Pain Sensitivity Questionnaire (PSQ) as well as a cold pressor test (CPT) which is used in clinical assessments of pain. Overall our online version of the PSQ performed similarly to the original pen-and-paper version. Construct validity of the PSQ total was demonstrated by internal consistency and consistent discrimination between more and less painful items. Criterion validity was demonstrated by correlation with pain sensitivity as measured by the CPT. Within the same cohort we performed a cold pressor test using a layperson description and household equipment. Comparison with published reports from controlled studies revealed similar distributions of cold pain tolerance times (i.e., time elapsed before removing the hand from the water). Of those who elected to participate in the CPT, a large majority of participants did not report issues with the test procedure or noncompliance with the instructions (97%). We confirmed a large sex difference in CPT thresholds in line with published data, such that women removed their hands from the water at a median of 54.2 seconds, with men lasting for a median time of 82.7 seconds (Kruskal-Wallis statistic, p < 0.0001), but other factors like age or current pain treatment were at most weakly associated, and inconsistently between men and women. We introduce a new paradigm for performing pain testing, called testing@home, that, in the case of cold nociception, showed comparable results to studies conducted under controlled conditions and supervision of a health care professional.
46
 Preprint DOI:	10.1101/19011957
 1

"Background: In England, national safety guidance recommends that ciclosporin, tacrolimus and diltiazem are prescribed by brand name due to their narrow therapeutic windows and, in the case of tacrolimus, to reduce the chance of organ transplantation rejection. Various small studies have shown that changes to electronic health records (EHR) interface can affect prescribing choices. . Objective: Our objectives were to assess variation by EHR system in breaches of safety guidance around prescribing of ciclosporin, tacrolimus and diltiazem; and to conduct user-interface research into the causes of such breaches. Methods: We carried out a retrospective cohort study using prescribing data in English primary care. Participants were English general practices and their respective electronic health records. The main outcome measures were (1) variation in ratio of breaching / adherent prescribing all practices (2) description of observations of EHR usage. Results: A total of 2,575,411 prescriptions were issued in 2018 for ciclosporin, tacrolimus and diltiazem (over 60mg); of these, 316,119 prescriptions breached NHS guidance (12.3%). Breaches were most common amongst users of the EMIS EHR (in 23.2% of ciclosporin & tacrolimus prescriptions, and 22.7% of diltiazem prescriptions); but breaches were observed in all EHRs. Conclusions: Design choices in EHR strongly influence safe prescribing of ciclosporin, tacrolimus and diltiazem; and breaches are prevalent in general practices in England. We recommend that all EHR vendors review their systems to increase safe prescribing of these medicines in line with national guidance. Almost all clinical practice is now mediated through an EHR system: further quantitative research into the effect of EHR design on clinical practice is long overdue."
47
 Preprint DOI:	10.1101/19012112

1
 "BackgroundIntranasal administration of the “prosocial” neuropeptide oxytocin is increasingly explored as a potential treatment for targeting the core characteristics of autism spectrum disorder (ASD). However, long-term follow-up studies, evaluating the possibility of long-lasting retention effects, are currently lacking.MethodsUsing a double-blind, randomized, placebo-controlled, parallel design, this pilot clinical trial explored the possibility of long-lasting behavioral effects of 4 weeks of intranasal oxytocin treatment (24 International Units once daily in the morning) in 40 adult men with ASD. To do so, self-report and informant-based questionnaires assessing core autism symptoms and characterizations of attachment were administered at baseline, immediately after 4 weeks of treatment (approximately 24 h after the last nasal spray administration), and at two follow-up sessions, 4 weeks and 1 year post-treatment.ResultsNo treatment-specific effects were identified in the primary outcome assessing social symptoms (Social Responsiveness Scale, self- and informant-rated). In particular, with respect to self-reported social responsiveness, improvements were evident both in the oxytocin and in the placebo group, yielding no significant between-group difference (p = .37). Also informant-rated improvements in social responsiveness were not significantly larger in the oxytocin, compared to the placebo group (between-group difference: p = .19). Among the secondary outcome measures, treatment-specific improvements were identified in the Repetitive Behavior Scale and State Adult Attachment Measure, indicating reductions in self-reported repetitive behaviors (p = .04) and reduced feelings of avoidance toward others (p = .03) in the oxytocin group compared to the placebo group, up to 1 month and even 1 year post-treatment. Treatment-specific effects were also revealed in screenings of mood states (Profile of Mood States), indicating higher reports of “vigor” (feeling energetic, active, lively) in the oxytocin, compared to the placebo group (p = .03).ConclusionsWhile no treatment-specific improvements were evident in terms of core social symptoms, the current observations of long-term beneficial effects on repetitive behaviors and feelings of avoidance are promising and suggestive of a therapeutic potential of oxytocin treatment for ASD. However, given the exploratory nature of this pilot study, future studies are warranted to evaluate the long-term effects of OT administration further.Trial registrationThe trial was registered with the European Clinical Trial Registry (Eudract 2014-000586-45) on January 22, 2014 (https://www.clinicaltrialsregister.eu/ctr-search/trial/2014-000586-45/BE)."
48
 Preprint DOI:	10.1101/19012435

1
 "Background: Asthma diagnosis in the community is often made without objective testing. Objective: The aim of this study was to evaluate the cost-effectiveness of implementing a stepwise objective diagnostic verification algorithm among patients with community-diagnosed asthma in the United States. Methods: We developed a probabilistic time-in-state cohort model that compared a stepwise asthma verification algorithm on the basis of spirometry testing and a methacholine challenge test against the current standard of care over 20 years. Model input parameters were informed from the literature and with original data analyses when required. The target population was US adults (=15 years old) with physician-diagnosed asthma. The final outcomes were costs (in 2018 dollars) and quality-adjusted life years (QALYs), discounted at 3% annually. Deterministic and probabilistic analyses were undertaken to examine the effect of alternative assumptions and uncertainty in model parameters on the results. Results: In a simulated cohort of 10,000 adults with diagnosed asthma, the stepwise algorithm resulted in removal of the diagnosis of 3,366. This was projected to be associated with savings of $36.26 million in direct costs and a gain of 4,049.28 QALYs over 20 years. Extrapolating these results to the US population indicated an undiscounted potential savings of $56.48 billion over 20 years. The results were robust against alternative assumptions and plausible changes in values of input parameters. Conclusion: Implementation of a simple diagnostic testing algorithm to verify asthma diagnosis might result in substantial savings and improvement in patients' quality of life."
49
 Preprint DOI:	10.1101/19012823

1
 "Purpose: The use of mobile health (mHealth) technologies to augment patient care enables providers to communicate remotely with patients enhancing the quality of care and patient engagement. Few studies evaluated predictive factors of its acceptance and subsequent implementation, especially in medically underserved populations. Methods: A cross-sectional study of 151 cancer patients was conducted at an academic medical center in the USA. A trained interviewer performed structured interviews regarding the barriers and facilitators of patients' current and desired use of mHealth technology for healthcare services. Results: Of the 151 participants, 35.8% were male and ages ranged from 21 to 104 years. 73.5% of participants currently have daily access to internet, and 68.2% currently own a smartphone capable of displaying mobile applications. Among all participants, acceptability of a daily mHealth application was significantly higher in patients with a college-level degree (OR 2.78, CI95% 1.25-5.88) and lower in patients > 80 years of age (OR 0.05, CI95% 0.01-0.23). Differences in acceptability when adjusted for current smartphone use and daily access to internet were nonsignificant. Among smartphone users, the desire to increase cancer knowledge was associated with a higher likelihood of utilizing a mHealth application (OR 261.53, CI95% 10.13-6748.71). ). Conclusion: The study suggests that factors such as age, educational achievement, , and access to internet are significant predictors of acceptability of a mHealth application among cancer patients. Healthcare organizations should consider these factors when launching patient engagement platforms."
50
 Preprint DOI:	10.1101/19012880
2

"Background: Recently, stereotactic radiosurgery has been applied to arrhythmias (stereotactic arrhythmia radioablation [STAR]), with promising results reported in patients with refractory scar-related ventricular tachycardia (VT), a cohort with known high morbidity and mortality. Objective: Herein, we describe our experience with STAR, detailing its early and mid- to long-term results. Methods: This is a pilot prospective study of patients undergoing STAR for refractory scar-related VT. The anatomical target for radioablation was defined on the basis of the clinical VT morphology, electroanatomic mapping, and study-specific preprocedural imaging with cardiac computed tomography. The target volume was treated with a prescription radiation dose of 25 Gy delivered in a single fraction by CyberKnife in an outpatient setting. Ventricular arrhythmias and radiation-related adverse events were monitored at follow-up to determine STAR efficacy and safety. Results: Five patients (100% men; mean age 63 ± 12 years; 80% with ischemic cardiomyopathy; left ventricular ejection fraction 34% ± 15%) underwent STAR. . Radioablation was delivered in 82 ± 11 minutes without acute complications. During a mean follow-up of 12 ± 2 months, all patients experienced clinically significant mid- to late-term ventricular arrhythmia recurrence; 2 patients died of complications associated with their advanced heart failure. There were no clinical or imaging evidence of radiation----induced complications in the organs at risk surrounding the scar targeted by radioablation. Conclusion: Despite good initial results, STAR did not result in effective arrhythmia control in the long term in a selected high-risk population of patients with scar-related VT. The safety profile was confirmed to be favorable, with no radiation-related complications observed during follow-up. Further studies are needed to explain these disappointing results."
51
 Preprint DOI:	10.1101/19013318
 1

"Background: Optimal ablation technique, including catheter-tissue contact during atrial fibrillation (AF) radiofrequency (RF) ablation, is associated with improved procedural outcomes. We used a custom developed software to analyze high-frequency catheter position data to study the interaction between catheter excursion during lesion placement, lesion-set sequentiality, and arrhythmia recurrence. Methods: A total of 100 consecutive patients undergoing first-time RF ablation for paroxysmal AF were analyzed. Spatial positioning of the ablation catheter sampled at 60 Hz during RF application was extracted from the CARTO3 system (Biosense Webster Inc, USA) and analyzed using custom-developed MATLAB software to determine precise catheter spatial 3D excursion during RF ablation. The primary end point was freedom from atrial arrhythmia lasting longer than 30 seconds after a single ablation procedure. Results: At 1 year, 86% of patients were free from recurrent arrhythmia. There was no significant difference in clinical, echocardiographic, or ablation characteristics between patients with and without recurrent arrhythmia. Analyzing 15,356,998 position data points revealed that lesion-set sequentiality and mean lesion catheter excursion were predictors of arrhythmia recurrence. Analyzing arrhythmia recurrence by mean single-lesion catheter excursion (excursion >2.81 mm) and by sequentiality (using 46% of lesions with interlesion distance >6 mm as cutoff) revealed significantly increased arrhythmia recurrence in the higher excursion group (23% vs 6%, P = .03) and in the less sequential group (24% vs 4%, P = .02). Conclusions: Ablation lesion sequentiality measured by catheter interlesion distance and catheter stability measured by catheter excursion during lesion placement are potentially modifiable factors affecting arrhythmia recurrence after RF ablation for AF."
52
 Preprint DOI:	10.1101/19013656
1

"ObjectiveTo investigate the relation between deep brain stimulation (DBS) of the posterior-subthalamic-area (PSA) and the ventral-intermediate-nucleus (VIM) and the distance to the dentatorubrothalamic tract (DRTT) in essential tremor (ET).MethodsTremor rating scale (TRS) hemi-scores were analyzed in 13 ET patients, stimulated in both the VIM and the PSA in a randomized, crossover trial. Distances of PSA and VIM contacts to population-based DRTTs were calculated. The relationships between distance to DRTT and stimulation amplitude, as well as DBS efficiency (TRS improvement per amplitude) were investigated.ResultsPSA contacts were closer to the DRTT (p = 0.019) and led to a greater improvement in TRS hemi-scores (p = 0.005) than VIM contacts. Proximity to the DRTT was related to lower amplitudes (p < 0.001) and higher DBS efficiency (p = 0.017).ConclusionsDifferences in tremor outcome and stimulation parameters between contacts in the PSA and the VIM can be explained by their different distance to the DRTT."
53
 Preprint DOI:	10.1101/2019.12.12.19014696
1

Renal cell carcinoma comprises a variety of entities, the most common being the clear-cell, papillary and chromophobe subtypes. These subtypes are related to different clinical evolution; however, most therapies have been developed for clear-cell carcinoma and there is not a specific treatment based on different subtypes. In this study, one hundred and sixty-four paraffin samples from primary nephrectomies for localized tumors were analyzed. MiRNAs were isolated and measured by microRNA arrays. Significance Analysis of Microarrays and Consensus Cluster algorithm were used to characterize different renal subtypes. The analyses showed that chromophobe renal tumors are a homogeneous group characterized by an overexpression of miR 1229, miR 10a, miR 182, miR 1208, miR 222, miR 221, miR 891b, miR 629-5p and miR 221-5p. On the other hand, clear cell renal carcinomas presented two different groups inside this histological subtype, with differences in miRNAs that regulate focal adhesion, transcription, apoptosis and angiogenesis processes. Specifically, one of the defined groups had an overexpression of proangiogenic microRNAs miR185, miR126 and miR130a. In conclusion, differences in miRNA expression profiles between histological renal subtypes were established. In addition, clear cell renal carcinomas had different expression of proangiogenic miRNAs. With the emergence of antiangiogenic drugs, these differences could be used as therapeutic targets in the future or as a selection method for tailoring personalized treatments.
54
 Preprint DOI:	10.1101/2019.12.13.876110
1

Timely completion of DNA replication is central to ac- curate cell division and to the maintenance of genomic stability. However, certain DNA-protein in- teractions can physically impede DNA replication fork progression. Cells remove or bypass these physical impediments by different mechanisms to preserve DNA macromolecule integrity and genome stability. In Saccharomyces cerevisiae, Wss1, the DNA-protein crosslink repair protease, allows cells to tolerate hydroxyurea-induced replication stress, but the underlying mechanism by which Wss1 pro- motes this function has remained unknown. Here, we report that Wss1 provides cells tolerance to repli- cation stress by directly degrading core histone subunits that non-specifically and non-covalently bind to single-stranded DNA. Unlike Wss1-depen- dent proteolysis of covalent DNA-protein crosslinks, proteolysis of histones does not require Cdc48 nor SUMO-binding activities. Wss1 thus acts as a multi- functional protease capable of targeting a broad range of covalent and non-covalent DNA-binding proteins to preserve genome stability during adverse conditions.
55
 Preprint DOI:	10.1101/2019.12.15.877035
1

Genome replication perturbs the DNA regulatory environment by displacing DNA-bound proteins, replacing nucleosomes, and introducing dosage imbalance between regions replicating at different S-phase stages. Recently, we showed that these effects are integrated to maintain transcription homeostasis: replicated genes increase in dosage, but their expression remains stable due to replication-dependent epigenetic changes that suppress transcription. Here, we examine whether reduced transcription from replicated DNA results from limited accessibility to regulatory factors by measuring the time-resolved binding of RNA polymerase II (Pol II) and specific transcription factors (TFs) to DNA during S phase in budding yeast. We show that the Pol II binding pattern is largely insensitive to DNA dosage, indicating limited binding to replicated DNA. In contrast, binding of three TFs (Reb1, Abf1, and Rap1) to DNA increases with the increasing DNA dosage. We conclude that the replication-specific chromatin environment remains accessible to regulatory factors but suppresses RNA polymerase recruitment.
56
 Preprint DOI:	10.1101/2019.12.17.19014415
1

"Background: Persistent formal thought disorder (FTD) is a core feature of schizophrenia. Recent cognitive and neuroimaging studies indicate a distinct mechanistic pathway underlying the persistent positive FTD (pFTD or disorganized thinking), though its structural determinants are still elusive. Using network-based cortical thickness estimates from ultra-high field 7-Tesla Magnetic Resonance Imaging (7T MRI), we investigated the structural correlates of pFTD. Methods: We obtained speech samples and 7T MRI anatomical scans from medicated clinically stable patients with schizophrenia (n = 19) and healthy controls (n = 20). Network-based morphometry was used to estimate the mean cortical thickness of 17 functional networks covering the entire cortical surface from each subject. We also quantified the vertexwise variability of thickness within each network to quantify the spatial coherence of the 17 networks, estimated patients vs. controls differences, and related the thickness of the affected networks to the severity of pFTD. Results: Patients had reduced thickness of the frontoparietal and default mode networks, and reduced spatial coherence affecting the salience and the frontoparietal control network. A higher burden of positive FTD related to reduced frontoparietal thickness and reduced spatial coherence of the salience network. The presence of positive FTD, but not its severity, related to the reduced thickness of the language network comprising of the superior temporal cortex. Conclusions: These results suggest that cortical thickness of both cognitive control and language networks underlie the positive FTD in schizophrenia. The structural integrity of cognitive control networks is a critical determinant of the expressed severity of persistent FTD in schizophrenia."
57
 Preprint DOI:	10.1101/2019.12.17.19015198
1

"Background: Hypertension is a common vascular disease and the main risk factor for cardiovascular diseases. Since the incidence of hypertension is rising in Ethiopia, one may expect that the household’s cost of healthcare services related to the disease will increase in the near future. Yet the cost associated with the disease is not known. We aimed to estimate the total cost of hypertension illness and identify associated factors among patients attending hospitals in Southwest Shewa zone, Oromia regional state, Ethiopia.Patients and Methods: An institution-based cross-sectional study design was employed to conduct the study from 13 August to 2 September 2018. All hypertensive patients aged 18 years and older who were on follow-up were eligible for this study. The total cost of hypertension illness was estimated by summing the direct and indirect costs. Bivariate and multivariate linear regression analyses were performed to identify factors associated with hypertension costs of illnesses.Results: A total of 349 patients participated in the study. T. The mean monthly total cost of hypertension illness was US$ 22.3 (95% CI, 21.3– 23.3). Direct and indirect costs constitute 51% and 49% of the total cost, respectively. The mean direct cost of hypertension illness per patient per month was US US$ 11.39 (95% CI, 10.6– 12.1). Out of these, drugs comprised higher cost (31%), followed by food (25%). The mean indirect cost per patient per month was US$ 10.89 (95% CI,10.4– 11.4). In this study, the primary educational status, family size (4– 6 and > 6), distance from hospital (≥ 10 km), the presence of a companion and stage of hypertension (stage two) of patients were identified as the predictors of the cost of hypertension illnesses.Conclusion: The cost of hypertension illness was very high when compared to the monthly income of households, exposing patients to catastrophic costs. Hence, the government should give due attention to protect patients from catastrophic health expenditures."
58
 Preprint DOI:	10.1101/2019.12.17.880302
1

Self-assembled photonic crystals have proven to be a fascinating class of photonic materials for nonabsorbing structural colorizations over large areas and in diverse relevant applications, including tools for on-chip spectrometers and biosensors, platforms for reflective displays, and templates for energy devices. The most prevalent building blocks for the self-assembly of photonic crystals are spherical colloids and block copolymers (BCPs) because of the generic appeal of these materials, which can be crafted into large-area 3D lattices. However, because of the intrinsic limitations of these structures, these two building blocks are difficult to assemble into a direct rod-connected diamond lattice, which is considered to be a champion photonic crystal. Here, we present a DNA origami-route for a direct rod-connected diamond photonic crystal exhibiting a complete photonic bandgap (PBG) in the visible regime. Using a combination of electromagnetic, phononic, and mechanical numerical analyses, we identify (i) the structural constraints of the 50 megadalton-scale giant DNA origami building blocks that could self-assemble into a direct rod-connected diamond lattice with high accuracy, and (ii) the elastic moduli that are essentials for maintaining lattice integrity in a buffer solution. A solution molding process could enable the transformation of the as-assembled DNA origami lattice into a porous silicon- or germanium-coated composite crystal with enhanced refractive index contrast, in that a champion relative bandwidth for the photonic bandgap (i.e., 0.29) could become possible even for a relatively low volume fraction (i.e., 16 vol %).
59
 Preprint DOI:	10.1101/2019.12.18.19014548
 1

"Clinical research in neurodevelopmental disorders remains reliant upon clinician and caregiver measures. Limitations of these approaches indicate a need for objective, quantitative, and reliable biomarkers to advance clinical research. Extant research suggests the potential utility of multiple candidate biomarkers; however, effective application of these markers in trials requires additional understanding of replicability, individual differences, and intra-individual stability over time. The Autism Biomarkers Consortium for Clinical Trials (ABC-CT) is a multi-site study designed to investigate a battery of electrophysiological (EEG) and eye-tracking (ET) indices as candidate biomarkers for autism spectrum disorder (ASD). The study complements published biomarker research through: inclusion of large, deeply phenotyped cohorts of children with ASD and typical development; a longitudinal design; a focus on well-evidenced candidate biomarkers harmonized with an independent sample; high levels of clinical, regulatory, technical, and statistical rigor; adoption of a governance structure incorporating diverse expertise in the ASD biomarker discovery and qualification process; prioritization of open science, including creation of a repository containing biomarker, clinical, and genetic data; and use of economical and scalable technologies that are applicable in developmental populations and those with special needs. The ABC-CT approach has yielded encouraging results, with one measure accepted into the FDA’s Biomarker Qualification Program to date. Through these advances, the ABC-CT and other biomarker studies in progress hold promise to deliver novel tools to improve clinical trials research in ASD."
60
 Preprint DOI:	10.1101/2019.12.18.881391
 1

Fusion with, and subsequent entry into, the host cell is one of the critical steps in the life cycle of enveloped viruses. For Middle East respiratory syndrome coronavirus (MERS-CoV), the spike (S) protein is the main determinant of viral entry. Proteolytic cleavage of the S protein exposes its fusion peptide (FP), which initiates the process of membrane fusion. Previous studies on the related severe acute respiratory syndrome coronavirus (SARS-CoV) FP have shown that calcium ions (Ca2+) play an important role in fusogenic activity via a Ca2+ binding pocket with conserved glutamic acid (E) and aspartic acid (D) residues. SARS-CoV and MERS-CoV FPs share a high sequence homology, and here, we investigated whether Ca2+ is required for MERS-CoV fusion by screening a mutant array in which E and D residues in the MERS-CoV FP were substituted with neutrally charged alanines (A). Upon verifying mutant cell surface expression and proteolytic cleavage, we tested their ability to mediate pseudoparticle (PP) infection of host cells in modulating Ca2+ environments. Our results demonstrate that intracellular Ca2+ enhances MERS-CoV wild-type (WT) PP infection by approximately 2-fold and that E891 is a crucial residue for Ca2+ interaction. Subsequent electron spin resonance (ESR) experiments revealed that this enhancement could be attributed to Ca2+ increasing MERS-CoV FP fusion-relevant membrane ordering. Intriguingly, isothermal calorimetry showed an approximate 1:1 MERS-CoV FP to Ca2+ ratio, as opposed to an 1:2 SARS-CoV FP to Ca2+ ratio, suggesting significant differences in FP Ca2+ interactions of MERS-CoV and SARS-CoV FP despite their high sequence similarity.
61
 Preprint DOI:	10.1101/2019.12.19.882274
1

In barley (Hordeum vulgare L.), Agrobacterium-mediated transformation efficiency is highly dependent on genotype with very few cultivars being amenable to transformation. Golden Promise is the cultivar most widely used for barley transformation and developing embryos are the most common donor tissue. We tested whether barley mutants with abnormally large embryos were more or less amenable to transformation and discovered that mutant M1460 had a transformation efficiency similar to that of Golden Promise. The large-embryo phenotype of M1460 is due to mutation at the LYS3 locus. There are three other barley lines with independent mutations at the same LYS3 locus, and one of these, Risø1508 has an identical missense mutation to that in M1460. However, none of the lys3 mutants except M1460 were transformable showing that the locus responsible for transformation efficiency, TRA1, was not LYS3 but another locus unique to M1460. To identify TRA1, we generated a segregating population by crossing M1460 to the cultivar Optic, which is recalcitrant to transformation. After four rounds of backcrossing to Optic, plants were genotyped and their progeny were tested for transformability. Some of the progeny lines were transformable at high efficiencies similar to those seen for the parent M1460 and some were not transformable, like Optic. A region on chromosome 2H inherited from M1460 is present in transformable lines only. We propose that one of the 225 genes in this region is TRA1.
62
 Preprint DOI:	10.1101/2019.12.30.19016154
1

Background: Whether the bone mineral density (BMD) T-score performs differently in osteoporosis classification in women of different genetic profiling and race background remains unclear. Methods: The genomic data in the Women’s Health Initiative study was analyzed (n = 2417). The polygenic score (PGS) was calculated from 63 BMD-associated single nucleotide polymorphisms (SNPs) for each participant. The World Health Organization′s (WHO) definition of osteoporosis (BMD T-score ≤ −2.5) was used to estimate the cumulative incidence of fracture. Results: T-score classification significantly underestimated the risk of major osteoporotic fracture (MOF) in the WHI study. An enormous underestimation was observed in African American women (POR: 0.52, 95% CI: 0.30–0.83) and in women with low PGS (predicted/observed ratio [POR]: 0.43, 95% CI: 0.28–0.64). Compared to Caucasian women, African American, African Indian, and Hispanic women respectively had a 59%, 41%, and 55% lower hazard of MOF after the T-score was adjusted for. The results were similar when used for any fractures. Conclusions: Our study suggested the BMD T-score performance varies significantly by race in postmenopausal women.
63
 Preprint DOI:	10.1101/2020.01.03.20016436

1

"Background: Ethiopia is a priority country of Gavi, the Vaccine Alliance to improve vaccination coverage and equitable uptake. The Ethiopian National Expanded Programme on Immunisation (EPI) and the Global Vaccine Action Plan set coverage goals of 90% at national level and 80% at district level by 2020. This study analyses full vaccination coverage among children in Ethiopia and estimates the equity impact by socioeconomic, geographic, maternal and child characteristics based on the 2016 Ethiopia Demographic and Health Survey dataset.Methods: Full vaccination coverage (1-dose BCG, 3-dose DTP3-HepB-Hib, 3-dose polio, 1-dose measles (MCV1), 3-dose pneumococcal (PCV3), and 2-dose rotavirus vaccines) of 2,004 children aged 12-23 months was analysed. Mean coverage was disaggregated by socioeconomic (household wealth, religion, ethnicity), geographic (area of residence, region), maternal (maternal age at birth, maternal education, maternal marital status, sex of household head), and child (sex of child, birth order) characteristics. Concentration indices estimated wealth and education-related inequities, and multiple logistic regression assessed associations between full vaccination coverage and socioeconomic, geographic, maternal, and child characteristics.Results: Full vaccination coverage was 33.3% [29.4-37.2] in 2016. Single vaccination coverage ranged from 49.1% [45.1-53.1] for PCV3 to 69.2% [65.5-72.8] for BCG. Wealth and maternal education related inequities were pronounced with concentration indices of 0.30 and 0.23 respectively. .Children in Addis Ababa and Dire Dawa were seven times more likely to have full vaccination compared to children living in the Afar region.... Children in female-headed households were 49% less likely to have full vaccination.Conclusion: Vaccination coverage in Ethiopia has a pro-advantaged regressive distribution with respect to both household wealth and maternal education. Children from poorer households, rural regions of Afar and Somali, no maternal education, and female-headed households had lower full vaccination coverage. Targeted programmes to reach under-immunised children in these subpopulations will improve vaccination coverage and equity outcomes in Ethiopia."
64
 Preprint DOI:	10.1101/2020.01.05.20016568
2

"PurposeTo provide a quantitative clinical-regulatory insight into the status of FDA orphan drug designations for compounds intended to treat lysosomal storage disorders (LSDs).MethodsAssessment of the drug pipeline through analysis of the FDA database for orphan drug designations with descriptive and comparative statistics.ResultsBetween 1983 and 2019, 124 orphan drug designations were granted by the FDA for compounds intended to treat 28 lysosomal storage diseases. Orphan drug designations focused on Gaucher disease (N = 16), Pompe disease (N = 16), Fabry disease (N = 10), MPS II (N = 10), MPS I (N = 9), and MPS IIIA (N = 9), and included enzyme replacement therapies, gene therapies, and small molecules, and others. Twenty-three orphan drugs were approved for the treatment of 11 LSDs. Gaucher disease (N = 6), cystinosis (N = 5), Pompe disease (N = 3), and Fabry disease (N = 2) had multiple approvals, CLN2, LAL-D, MPS I, II, IVA, VI, and VII one approval each. This is an increase of nine more approved drugs and four more treatable LSDs (CLN2, MPS VII, LAL-D, and MPS IVA) since 2013. Mean time between orphan drug designation and FDA approval was 89.7 SD 55.00 (range 8–203, N = 23) months.ConclusionsThe drug development pipeline for LSDs is growing and evolving, with increased focus on diverse small-molecule targets and gene therapy. CLN2 was the first and only LSD with an approved therapy directly targeted to the brain. Newly approved products included “me-too”–enzymes and innovative compounds such as the first pharmacological chaperone for the treatment of Fabry disease."
65
 Preprint DOI:	10.1101/2020.01.07.20016816
1

Frailty indices (FIs) based on continuous valued health data, such as obtained from blood and urine tests, have been shown to be predictive of adverse health outcomes. However, creating FIs from such biomarker data requires a binarization treatment that is difficult to standardize across studies. In this work, we explore a “quantile” methodology for the generic treatment of biomarker data that allows us to construct an FI without preexisting medical knowledge (i.e. risk thresholds) of the included biomarkers. We show that our quantile approach performs as well as, or even slightly better than, established methods for the National Health and Nutrition Examination Survey and the Canadian Study of Health and Aging data sets. Furthermore, we show that our approach is robust to cohort effects within studies as compared to other data-based methods. The success of our binarization approaches provides insight into the robustness of the FI as a health measure, and the upper limits of the FI observed in various data sets, and also highlights general difficulties in obtaining absolute scales for comparing FIs between studies.
66
 Preprint DOI:	10.1101/2020.01.07.897496
1

An understanding of host–parasite interactions represents an important, but often overlooked, axis for predicting how polar marine biodiversity may be impacted by continued environmental change over the next century. Here, we survey three species of crocodile icefish (Notothenioidei: Channichthyidae) collected from two island archipelagos in the southern Scotia Arc region for evidence of leech infestations. Specifically, we report on infestation prevalence, intensity, spatial patterns of relative abundances, size distribution of parasitized fish, and patterns of host and attachment site specificity. Our results reveal high levels of attachment area fidelity for each leech species. These results suggest skin thickness and density of the vascular network constrain leech attachment sites and further suggest trophic (i.e., post-cyclic) transmission to be an important axis of parasitization. We also demonstrate that, while leech species appear to be clustered spatially, this clustering does not appear to be correlated with fish biomass. This study illuminates the complex interactions among fish hosts and leech parasites in the Southern Ocean and lays the groundwork for future studies of Antarctic marine leech ecology that can aid in forecasting how host–parasite interactions may shift in the face of ongoing climate change.
67
 Preprint DOI:	10.1101/2020.01.13.20016360
1

"The locus coeruleus is the major source of noradrenaline to the brain and contributes to a wide range of physiological and cognitive functions including arousal, attention, autonomic control, and adaptive behaviour. Neurodegeneration and pathological aggregation of tau protein in the locus coeruleus are early features of progressive supranuclear palsy (PSP). This pathology is proposed to contribute to the clinical expression of disease, including the PSP Richardson’s syndrome. We test the hypothesis that tau pathology and neuronal loss are associated with clinical heterogeneity and severity in PSP.We used immunohistochemistry in post mortem tissues from 31 patients with a clinical diagnosis of PSP (22 with Richardson’s syndrome) and 6 control cases. We quantified the presence of hyperphosphorylated tau, the number of pigmented cells indicative of noradrenergic neurons, and the percentage of pigmented neurons with tau-positive inclusions. Ante mortem assessment of clinical severity using the PSP rating scale was available within 1.8 (±0.9) years for 23 patients.We found an average 49% reduction of pigmented neurons in PSP patients relative to controls. The loss of pigmented neurons correlated with disease severity, even after adjusting for disease duration and the interval between clinical assessment and death. The degree of neuronal loss was negatively associated with tau-positive inclusions, with an average of 44% of pigmented neurons displaying tau-inclusions.Degeneration and tau pathology in the locus coeruleus are related to clinical heterogeneity of PSP. The noradrenergic deficit in the locus coeruleus is a candidate target for pharmacological treatment. Recent developments in ultra-high field magnetic resonance imaging to quantify in vivo structural integrity of the locus coeruleus may provide biomarkers for noradrenergic experimental medicines studies in PSP."
68
 Preprint DOI:	10.1101/2020.01.13.905190
1

Hemophilia A, an X-linked bleeding disorder caused by deficiency of factor VIII (FVIII), is treated by protein replacement. Unfortunately, this regimen is costly due to the expense of producing recombinant FVIII as a consequence of its low-level secretion from mammalian host cells. FVIII expression activates the endoplasmic reticulum (ER) stress response, causes oxidative stress, and induces apoptosis. Importantly, little is known about the factors that cause protein misfolding and aggregation in metazoans. Here, we identified intrinsic and extrinsic factors that cause FVIII to form aggregates. We show that FVIII forms amyloid-like fibrils within the ER lumen upon increased FVIII synthesis or inhibition of glucose metabolism. Significantly, FVIII amyloids can be dissolved upon restoration of glucose metabolism to produce functional secreted FVIII. Two ER chaperone families and their cochaperones, immunoglobulin binding protein (BiP) and calnexin/calreticulin, promote FVIII solubility in the ER, where the former is also required for disaggregation. A short aggregation motif in the FVIII A1 domain (termed Aggron) is necessary and sufficient to seed β-sheet polymerization, and BiP binding to this Aggron prevents amyloidogenesis. Our findings provide novel insight into mechanisms that limit FVIII secretion and ER protein aggregation in general and have implication for ongoing hemophilia A gene-therapy clinical trials.
69
 Preprint DOI:	10.1101/2020.01.21.20018101
1

Multiple genes have been associated with monogenic Parkinson's disease and Parkinsonism syndromes. Mutations in PINK1 (PARK6) have been shown to result in autosomal recessive early-onset Parkinson's disease. In the past decade, several studies have suggested that carrying a single heterozygous PINK1 mutation is associated with increased risk for Parkinson's disease. Here, we comprehensively assess the role of PINK1 variants in Parkinson's disease susceptibility using several large data sets totalling 376,558 individuals including 13,708 cases with Parkinson's disease and 362,850 control subjects. After combining these data, we did not find evidence to support a role for heterozygous PINK1 mutations as a robust risk factor for Parkinson's disease.
70
 Preprint DOI:	10.1101/2020.01.21.914929
1

Bunyaviruses are significant human pathogens, causing diseases ranging from hemorrhagic fevers to encephalitis. Among these viruses, La Crosse virus (LACV), a member of the California serogroup, circulates in the eastern and midwestern United States. While LACV infection is often asymptomatic, dozens of cases of encephalitis are reported yearly. Unfortunately, no antivirals have been approved to treat LACV infection. Here, we developed a method to rapidly test potential antivirals against LACV infection. From this screen, we identified several potential antiviral molecules, including known antivirals. Additionally, we identified many novel antivirals that exhibited antiviral activity without affecting cellular viability. Valinomycin, a potassium ionophore, was among our top targets. We found that valinomycin exhibited potent anti-LACV activity in multiple cell types in a dose-dependent manner. Valinomycin did not affect particle stability or infectivity, suggesting that it may preclude virus replication by altering cellular potassium ions, a known determinant of LACV entry. We extended these results to other ionophores and found that the antiviral activity of valinomycin extended to other viral families, including bunyaviruses (Rift Valley fever virus, Keystone virus), enteroviruses (coxsackievirus, rhinovirus), flavirivuses (Zika virus), and coronaviruses (human coronavirus 229E [HCoV-229E] and Middle East respiratory syndrome CoV [MERS-CoV]). In all viral infections, we observed significant reductions in virus titer in valinomycin-treated cells. In sum, we demonstrate the importance of potassium ions to virus infection, suggesting a potential therapeutic target to disrupt virus replication.
71
 Preprint DOI:	10.1101/2020.01.22.914952
2

Since the outbreak of severe acute respiratory syndrome (SARS) 18 years ago, a large number of SARS-related coronaviruses (SARSr-CoVs) have been discovered in their natural reservoir host, bats1,2,3,4. Previous studies have shown that some bat SARSr-CoVs have the potential to infect humans5,6,7. Here we report the identification and characterization of a new coronavirus (2019-nCoV), which caused an epidemic of acute respiratory syndrome in humans in Wuhan, China. The epidemic, which started on 12 December 2019, had caused 2,794 laboratory-confirmed infections including 80 deaths by 26 January 2020. Full-length genome sequences were obtained from five patients at an early stage of the outbreak. The sequences are almost identical and share 79.6% % sequence identity to SARS-CoV. Furthermore, we show that 2019-nCoV is 96% identical at the whole-genome level to a bat coronavirus. Pairwise protein sequence analysis of seven conserved non-structural proteins domains show that this virus belongs to the species of SARSr-CoV. In addition, 2019-nCoV virus isolated from the bronchoalveolar lavage fluid of a critically ill patient could be neutralized by sera from several patients. Notably, we confirmed that 2019-nCoV uses the same cell entry receptor—angiotensin converting enzyme II (ACE2)—as SARS-CoV.
72
 Preprint DOI:	10.1101/2020.01.22.915660
1

Over the past 20 years, several coronaviruses have crossed the species barrier into humans, causing outbreaks of severe, and often fatal, respiratory illness. Since SARS-CoV was first identified in animal markets, global viromics projects have discovered thousands of coronavirus sequences in diverse animals and geographic regions. Unfortunately, there are few tools available to functionally test these viruses for their ability to infect humans, which has severely hampered efforts to predict the next zoonotic viral outbreak. Here, we developed an approach to rapidly screen lineage B betacoronaviruses, such as SARS-CoV and the recent SARS-CoV-2, for receptor usage and their ability to infect cell types from different species. We show that host protease processing during viral entry is a significant barrier for several lineage B viruses and that bypassing this barrier allows several lineage B viruses to enter human cells through an unknown receptor. We also demonstrate how different lineage B viruses can recombine to gain entry into human cells, and confirm that human ACE2 is the receptor for the recently emerging SARS-CoV-2.
73
 Preprint DOI:	10.1101/2020.01.23.20017566
1

"Introduction Somatic mutations in STK11 and KEAP1, frequently comutated in non-squamous non-small cell lung cancer (NSQ NSCLC), have been associated with poor response to immune checkpoint blockade (ICB). However, previous reports lack non-ICB controls needed to properly ascertain the predictive nature of those biomarkers. The objective of this study was to evaluate the predictive versus prognostic effect of STK11 or KEAP1 mutations in NSQ NSCLC.Methods Patients diagnosed with stage IIIB, IIIC, IVA or IVB NSQ NSCLC from a real-world data cohort from the Flatiron Health Network linked with genetic testing from Foundation Medicine were retrospectively assessed. Real-world, progression-free survival (rwPFS) and overall survival (OS) were calculated from time of initiation of first-line treatment.Results We analysed clinical and mutational data for 2276 patients including patients treated with anti-programmed death-1 (PD-1)/anti-programmed death ligand 1 (PD-L1) inhibitors at first line (n=574). Mutations in STK11 or KEAP1 were associated with poor outcomes across multiple therapeutic classes and were not specifically associated with poor outcomes in ICB cohorts. There was no observable interaction between STK11 mutations and anti-PD-1/anti-PD-L1 treatment on rwPFSrwPFSrwPFSrwPFSrwPFSrwPFSrwPFSrwPFS (HR, 1.05; 95% CI 0.76 to 1.44; p=0.785) or OS (HR, 1.13; 95% CI 0.76 to 1.67; p=0.540).)). .)..). Similarly, there was no observable interaction between KEAP1 mutations and treatment on rwPFS (HR, 0.93; 95% CI 0.67 to 1.28; p=0.653) or OS (HR, 0.98; 95% CI 0.66 to 1.45; p=0.913))..).))..).Conclusion Our results show that STK11-KEAP1 mutations are prognostic, not predictive, biomarkers for anti-PD-1/anti-PD-L1 therapy."
74
 Preprint DOI:	10.1101/2020.01.23.916395
2

"BackgroundsAn ongoing outbreak of a novel coronavirus (2019-nCoV) pneumonia hit a major city in China, Wuhan, December 2019 and subsequently reached other provinces/regions of China and other countries. We present estimates of the basic reproduction number, R0, of 2019-nCoV in the early phase of the outbreak.MethodsAccounting for the impact of the variations in disease reporting rate, we modelled the epidemic curve of 2019-nCoV cases time series, in mainland China from January 10 to January 24, 2020, through the exponential growth. With the estimated intrinsic growth rate (γ), we estimated R0 by using the serial intervals (SI) of two other well-known coronavirus diseases, MERS and SARS, as approximations for the true unknown SI.FindingsThe early outbreak data largely follows the exponential growth. We estimated that the mean R0 ranges from 2.24 (95%CI: 1.96–2.55) to 3.58 (95%CI: 2.89–4.39) ) ) ) ) ) ) associated with 8-fold to 2-fold increase in the reporting rate. We demonstrated that changes in reporting rate substantially affect estimates of R0....ConclusionThe mean estimate of R0 for the 2019-nCoV ranges from 2.24 to 3.58, , and is significantly larger than 1. Our findings indicate the potential of 2019-nCoV to cause outbreaks."
75
 Preprint DOI:	10.1101/2020.01.24.915157
1

There is a worldwide concern about the new coronavirus 2019‐nCoV as a global public health threat. In this article, we provide a preliminary evolutionary and molecular epidemiological analysis of this new virus. A phylogenetic tree has been built using the 15 available whole genome sequences of 2019‐nCoV, 12 whole genome sequences of 2019‐nCoV, and 12 highly similar whole genome sequences available in gene bank (five from the severe acute respiratory syndrome, two from Middle East respiratory syndrome, and five from bat SARS‐like coronavirus). Fast unconstrained Bayesian approximation analysis shows that the nucleocapsid and the spike glycoprotein have some sites under positive pressure, whereas homology modeling revealed some molecular and structural differences between the viruses. The phylogenetic tree showed that 2019‐nCoV significantly clustered with bat SARS‐like coronavirus sequence isolated in 2015, whereas structural analysis revealed mutation in Spike Glycoprotein and nucleocapsid protein. From these results, the new 2019‐nCoV is distinct from SARS virus, probably trasmitted from bats after mutation conferring ability to infect humans.
76
 Preprint DOI:	10.1101/2020.01.24.919159
1

The outbreak of pneumonia originating in Wuhan, China, has generated 24,500 confirmed cases, including 492 deaths, as of 5 February 2020. The virus (2019-nCoV) has spread elsewhere in China and to 24 countries, including South Korea, Thailand, Japan and USA. Fortunately, there has only been limited human-to-human transmission outside of China. Here, we assess the risk of sustained transmission whenever the coronavirus arrives in other countries. Data describing the times from symptom onset to hospitalisation for 47 patients infected early in the current outbreak are used to generate an estimate for the probability that an imported case is followed by sustained human-to-human transmission. Under the assumptions that the imported case is representative of the patients in China, and that the 2019-nCoV is similarly transmissible to the SARS coronavirus, the probability that an imported case is followed by sustained human-to-human transmission is 0.41 (credible interval [0.27, 0.55]).]).]).])]). . However, if the mean time from symptom onset to hospitalisation can be halved by intense surveillance, then the probability that an imported case leads to sustained transmission is only 0.012 (credible interval [0, 0.099]). ]). ]). This emphasises the importance of current surveillance efforts in countries around the world, to ensure that the ongoing outbreak will not become a global pandemic.
77
 Preprint DOI:	10.1101/2020.01.26.20018754
2

The geographic spread of 2019 novel coronavirus (COVID-19) infections from the epicenter of Wuhan, China, , has provided an opportunity to study the natural history of the recently emerged virus. Using publicly available event-date data from the ongoing epidemic, the present study investigated the incubation period and other time intervals that govern the epidemiological dynamics of COVID-19 infections. Our results show that the incubation period falls within the range of 2–14 days with 95% confidence and has a mean of around 5 days when approximated using the best-fit lognormal distribution. The mean time from illness onset to hospital admission (for treatment and/or isolation) was estimated at 3–4 days without truncation and at 5–9 days when right truncated. Based on the 95th percentile estimate of the incubation period, we recommend that the length of quarantine should be at least 14 days.. The median time delay of 13 days from illness onset to death (17 days with right truncation) ) should be considered when estimating the COVID-19 case fatality risk.
78
 Preprint DOI:	10.1101/2020.01.26.920249
1

"Background: A novel coronavirus (2019-nCoV) associated with human to human transmission and severe human infection has been recently reported from the city of Wuhan in China. Our objectives were to characterize the genetic relationships of the 2019-nCoV and to search for putative recombination within the subgenus of sarbecovirus.Methods: Putative recombination was investigated by RDP4 and Simplot v3.5.1 and discordant phylogenetic clustering in individual genomic fragments was confirmed by phylogenetic analysis using maximum likelihood and Bayesian methods.Results: Our analysis suggests that the 2019-nCoV although closely related to BatCoV RaTG13 sequence throughout the genome (sequence similarity 96.3%), shows discordant clustering with the Bat_SARS-like coronavirus sequences. Specifically, in the 5'-part spanning the first 11,498 nucleotides and the last 3'-part spanning 24,341-30,696 positions, 2019-nCoV and RaTG13 formed a single cluster with Bat_SARS-like coronavirus sequences, whereas in the middle region spanning the 3'-end of ORF1a, the ORF1b and almost half of the spike regions, 2019-nCoV and RaTG13 grouped in a separate distant lineage within the sarbecovirus branch.Conclusions: The levels of genetic similarity between the 2019-nCoV and RaTG13 suggest that the latter does not provide the exact variant that caused the outbreak in humans, but the hypothesis that 2019-nCoV has originated from bats is very likely. We show evidence that the novel coronavirus (2019-nCov) is not-mosaic consisting in almost half of its genome of a distinct lineage within the betacoronavirus. These genomic features and their potential association with virus characteristics and virulence in humans need further attention."
79
 Preprint DOI:	10.1101/2020.01.27.20018689
3

Transcutaneous cervical vagal nerve stimulation (tcVNS) devices are attractive alternatives to surgical implants, and can be applied for a number of conditions in ambulatory settings, including stress-related neuropsychiatric disorders. Transferring tcVNS technologies to at-home settings brings challenges associated with the assessment of therapy response. The ability to accurately detect whether tcVNS has been effectively delivered in a remote setting such as the home has never been investigated. We designed and conducted a study in which 12 human subjects received active tcVNS and 14 received sham stimulation in tandem with traumatic stress, and measured continuous cardiopulmonary signals including the electrocardiogram (ECG), photoplethysmogram (PPG), seismocardiogram (SCG), and respiratory effort (RSP). We extracted physiological parameters related to autonomic nervous system activity, and created a feature set from these parameters to: 1) detect active (vs. sham) tcVNS stimulation presence with machine learning methods, and 2) determine which sensing modalities and features provide the most salient markers of tcVNS-based changes in physiological signals. Heart rate (ECG), vasomotor activity (PPG), and pulse arrival time (ECG+PPG) provided sufficient information to determine target engagement (compared to sham) in addition to other combinations of sensors. resulting in 96% accuracy, precision, and recall with a receiver operator characteristics area of 0.96. Two commonly utilized sensing modalities (ECG and PPG) that are suitable for home use can provide useful information on therapy response for tcVNS. The methods presented herein could be deployed in wearable devices to quantify adherence for at-home use of tcVNS technologies.
80
 Preprint DOI:	10.1101/2020.01.27.20018986
2

A novel coronavirus (2019-nCoV) is causing an outbreak of viral pneumonia that started in Wuhan, China. Using the travel history and symptom onset of 88 confirmed cases that were detected outside Wuhan in the early outbreak phase, we estimate the mean incubation period to be 6.4 days (95% credible interval: 5.6–7.7),)), , ranging from 2.1 to 11.1 days (2.5th to 97.5th percentile). These values should help inform 2019-nCoV case definitions and appropriate quarantine durations.
81
 Preprint DOI:	10.1101/2020.01.28.923011
1

The newly identified 2019 novel coronavirus (2019-nCoV) has caused more than 11,900 laboratory-confirmed human infections, including 259 deaths, posing a serious threat to human health. Currently, however, there is no specific antiviral treatment or vaccine. Considering the relatively high identity of receptor-binding domain (RBD) in 2019-nCoV and SARS-CoV, it is urgent to assess the cross-reactivity of anti-SARS CoV antibodies with 2019-nCoV spike protein, which could have important implications for rapid development of vaccines and therapeutic antibodies against 2019-nCoV. Here, we report for the first time that a SARS-CoV-specific human monoclonal antibody, CR3022, could bind potently with 2019-nCoV RBD (KD of 6.3 nM). The epitope of CR3022 does not overlap with the ACE2 binding site within 2019-nCoV RBD. These results suggest that CR3022 may havethe potential to be developed as candidate therapeutics, alone or in combination with other neutralizing antibodies, for the prevention and treatment of 2019-nCoV infections. Interestingly, some of the most potent SARS-CoV-specific neutralizing antibodies (e.g. m396, CR3014) that target the ACE2 binding site of SARS-CoV failed to bind 2019-nCoV spike protein, implying that the difference in the RBD of SARS-CoV and 2019-nCoV has a critical impact for the cross-reactivity of neutralizing antibodies, and that it is still necessary to develop novel monoclonal antibodies that could bind specifically to 2019-nCoV RBD.
82
 Preprint DOI:	10.1101/2020.01.31.20019901
2

"BackgroundAn outbreak of severe acute respiratory syndrome coronavirus 2 (SARS-CoV-2) has led to 95 333 confirmed cases as of March 5, 2020. Understanding the early transmission dynamics of the infection and evaluating the effectiveness of control measures is crucial for assessing the potential for sustained transmission to occur in new areas. Combining a mathematical model of severe SARS-CoV-2 transmission with four datasets from within and outside Wuhan, we estimated how transmission in Wuhan varied between December, 2019, and February, 2020. . We used these estimates to assess the potential for sustained human-to-human transmission to occur in locations outside Wuhan if cases were introducedintroducedintroduced.MethodsWe combined a stochastic transmission model with data on cases of coronavirus disease 2019 (COVID-19) in Wuhan and international cases that originated in Wuhan to estimate how transmission had varied over time during January, 2020, and February, 2020. Based on these estimates, we then calculated the probability that newly introduced cases might generate outbreaks in other areas. To estimate the early dynamics of transmission in Wuhan, we fitted a stochastic transmission dynamic model to multiple publicly available datasets on cases in Wuhan and internationally exported cases from Wuhan. The four datasets we fitted to were: daily number of new internationally exported cases (or lack thereof), by date of onset, as of Jan 26, 2020; daily number of new cases in Wuhan with no market exposure, by date of onset, between Dec 1, 2019, and Jan 1, 2020; daily number of new cases in China, by date of onset, between Dec 29, 2019, and Jan 23, 2020; and proportion of infected passengers on evacuation flights between Jan 29, 2020, and Feb 4, 2020. . We used an additional two datasets for comparison with model outputs: daily number of new exported cases from Wuhan (or lack thereof) in countries with high connectivity to Wuhan (ie, top 20 most at-risk countries), by date of confirmation, as of Feb 10, 2020; and data on new confirmed cases reported in Wuhan between Jan 16, 2020, and Feb 11, 2020.FinFindingsWe estimated that the median daily reproduction number (Rt) in Wuhan declined from 2·35 (95% CI 1·15–4·77) 1 week before travel restrictions were introduced on Jan 23, 2020, to 1·05 (0·41–2·39) 1 week after.Based on our estimates of Rt, assuming SARS-like variation, , we calculated that in locations with similar transmission potential to Wuhan in early January, once there are at least four independently introduced cases, there is a more than 50% chance the infection will establish within that population.InterpretationOur results show that COVID-19 transmission probably declined in Wuhan during late January, 2020, coinciding with the introduction of travel control measures. . As more cases arrive in international locations with similar transmission potential to Wuhan before these control measures, it is likely many chains of transmission will fail to establish initially, but might lead to new outbreaks eventually."
83
 Preprint DOI:	10.1101/2020.01.31.928796
1

Genome detective is a web-based, user-friendly software application to quickly and accurately assemble all known virus genomes from next-generation sequencing datasets. This application allows the identification of phylogenetic clusters and genotypes from assembled genomes in FASTA format. Since its release in 2019, we have produced a number of typing tools for emergent viruses that have caused large outbreaks, such as Zika and Yellow Fever Virus in Brazil. Here, we present the Genome Detective Coronavirus Typing Tool that can accurately identify the novel severe acute respiratory syndrome (SARS)-related coronavirus (SARS-CoV-2) sequences isolated in China and around the world. The tool can accept up to 2000 sequences per submission and the analysis of a new whole-genome sequence will take approximately 1 min. The tool has been tested and validated with hundreds of whole genomes from 10 coronavirus species, and correctly classified all of the SARS-related coronavirus (SARSr-CoV) and all of the available public data for SARS-CoV-2. The tool also allows tracking of new viral mutations as the outbreak expands globally, which may help to accelerate the development of novel diagnostics, drugs and vaccines to stop the COVID-19 disease.
84
 Preprint DOI:	10.1101/2020.02.02.931162
2

There is a rising global concern for the recently emerged novel coronavirus (2019‐nCoV). Full genomic sequences have been released by the worldwide scientific community in the last few weeks to understand the evolutionary origin and molecular characteristics of this virus. Taking advantage of all the genomic information currently available, we constructed a phylogenetic tree including also representatives of other coronaviridae, such as Bat coronavirus (BCoV) and severe acute respiratory syndrome. We confirm high sequence similarity (>99%) between all sequenced 2019‐nCoVs genomes available, with the closest BCoV sequence sharing 96.2% sequence identity, confirming the notion of a zoonotic origin of 2019‐nCoV. Despite the low heterogeneity of the 2019‐nCoV genomes, we could identify at least two hypervariable genomic hotspots, one of which is responsible for a Serine/Leucine variation in the viral ORF8‐encoded protein. Finally, we perform a full proteomic comparison with other coronaviridae, identifying key aminoacidic differences to be considered for antiviral strategies deriving from previous anti‐coronavirus approaches.	.
86
 Preprint DOI:	10.1101/2020.02.03.933226
3

The beginning of 2020 has seen the emergence of COVID-19 outbreak caused by a novel coronavirus, Severe Acute Respiratory Syndrome Coronavirus 2 (SARS-CoV-2). There is an imminent need to better understand this new virus and to develop ways to control its spread. In this study, we sought to gain insights for vaccine design against SARS-CoV-2 by considering the high genetic similarity between SARS-CoV-2 and SARS-CoV, which caused the outbreak in 2003, and leveraging existing immunological studies of SARS-CoV. By screening the experimentally-determined SARS-CoV-derived B cell and T cell epitopes in the immunogenic structural proteins of SARS-CoV, we identified a set of B cell and T cell epitopes derived from the spike (S) and nucleocapsid (N) proteins that map identically to SARS-CoV-2 proteins. As no mutation has been observed in these identified epitopes among the 120 available SARS-CoV-2 sequences (as of 21 February 2020),),),), immune targeting of these epitopes may potentially offer protection against this novel virus. For the T cell epitopes, we performed a population coverage analysis of the associated MHC alleles and proposed a set of epitopes that is estimated to provide broad coverage globally, as well as in China. Our findings provide a screened set of epitopes that can help guide experimental efforts towards the development of vaccines against SARS-CoV-2.

The beginning of 2020 has seen the emergence of COVID-19 outbreak caused by a novel coronavirus, Severe Acute Respiratory Syndrome Coronavirus 2 (SARS-CoV-2). There is an imminent need to better understand this new virus and to develop ways to control its spread. In this study, we sought to gain insights for vaccine design against SARS-CoV-2 by considering the high genetic similarity between SARS-CoV-2 and SARS-CoV, which caused the outbreak in 2003, and leveraging existing immunological studies of SARS-CoV. By screening the experimentally-determined SARS-CoV-derived B cell and T cell epitopes in the immunogenic structural proteins of SARS-CoV, we identified a set of B cell and T cell epitopes derived from the spike (S) and nucleocapsid (N) proteins that map identically to SARS-CoV-2 proteins. As no mutation has been observed in these identified epitopes among the 120 available SARS-CoV-2 sequences (as of 21 February 2020), immune targeting of these epitopes may potentially offer protection against this novel virus. For the T cell epitopes, we performed a population coverage analysis of the associated MHC alleles and proposed a set of epitopes that is estimated to provide broad coverage globally, as well as in China. Our findings provide a screened set of epitopes that can help guide experimental efforts towards the development of vaccines against SARS-CoV-2.
87
 Preprint DOI:	10.1101/2020.02.04.20020503
2

We developed a computational tool to assess the risks of novel coronavirus outbreaks outside of China. We estimate the dependence of the risk of a major outbreak in a country from imported cases on key parameters such as: (i) the evolution of the cumulative number of cases in mainland China outside the closed areas; (ii) the connectivity of the destination country with China, including baseline travel frequencies, the effect of travel restrictions, and the efficacy of entry screening at destination; and (iii) the efficacy of control measures in the destination country (expressed by the local reproduction number Rloc ). We found that in countries with low connectivity to China but with relatively high Rloc , the most beneficial control measure to reduce the risk of outbreaks is a further reduction in their importation number either by entry screening or travel restrictions. Countries with high connectivity but low Rloc benefit the most from policies that further reduce Rloc . Countries in the middle should consider a combination of such policies. Risk assessments were illustrated for selected groups of countries from America, Asia, and Europe. We investigated how their risks depend on those parameters, and how the risk is increasing in time as the number of cases in China is growing.
88
 Preprint DOI:	10.1101/2020.02.06.20020974
1

"BACKGROUNDSince December 2019, when coronavirus disease 2019 (Covid-19) emerged in Wuhan city and rapidly spread throughout China, data have been needed on the clinical characteristics of the affected patients.METHODSWe extracted data regarding 1099 patients with laboratory-confirmed Covid-19 from 552 hospitals in 30 provinces, autonomous regions,, and municipalities in mainland China through January 29, 2020. The primary composite end point was admission to an intensive care unit (ICU), the use of mechanical ventilation, or death.deathdeathdeath.RESULTSThe median age of the patients was 47 years; 41.9% of the patients were female. The primary composite end point occurred in 67 patients (6.1%), including 5.0% who were admitted to the ICU, 2.3% who underwent invasive mechanical ventilation, and 1.4% who died. . Only 1.9% of the patients had a history of direct contact with wildlife. Among nonresidents of Wuhan, 72.3% had contact with residents of Wuhan, including 31.3% who had visited the city. The most common symptoms were fever (43.8% on admission and 88.7% during hospitalization) and cough (67.8%).Diarrhea was uncommon (3.8%). The median incubation period was 4 days (interquartile range, 2 to 7). ). On admission, ground-glass opacity was the most common radiologic finding on chest computed tomography (CT) (56.4%). No radiographic or CT abnormality was found in 157 of 877 patients (17.9%) with nonsevere disease and in 5 of 173 patients (2.9%) with severe disease. Lymphocytopenia was present in 83.2% of the patients on admission..CONCLUSIONSDuring the first 2 months of the current outbreak, Covid-19 spread rapidly throughout China and caused varying degrees of illness. . Patients often presented without fever, and many did not have abnormal radiologic findings. (Funded by the National Health Commission of China and others.)"
89
 Preprint DOI:	10.1101/2020.02.10.20021758
1

"Objective: Widespread metabolic changes are seen in neurodegenerative disease and could be used as biomarkers for diagnosis and disease monitoring. They may also reveal disease mechanisms that could be a target for therapy. In this study we looked for blood-based biomarkers in syndromes associated with frontotemporal lobar degeneration. Methods: Plasma metabolomic profiles were measured from 134 patients with a syndrome associated with frontotemporal lobar degeneration (behavioural variant frontotemporal dementia n = 30, non fluent variant primary progressive aphasia n = 26, progressive supranuclear palsy n = 45, corticobasal syndrome n = 33) and 32 healthy controls. Results: Forty-nine of 842 metabolites were significantly altered in frontotemporal lobar degeneration syndromes (after false-discovery rate correction for multiple comparisons). These were distributed across a wide range of metabolic pathways including amino acids, energy and carbohydrate, cofactor and vitamin, lipid and nucleotide pathways. The metabolomic profile supported classification between frontotemporal lobar degeneration syndromes and controls with high accuracy (88.1-96.6%) while classification accuracy was lower between the frontotemporal lobar degeneration syndromes (72.1-83.3%). One metabolic profile, comprising a range of different pathways, was consistently identified as a feature of each disease versus controls: the degree to which a patient expressed this metabolomic profile was associated with their subsequent survival (hazard ratio 0.74 [0.59-0.93], p = 0.0018). Conclusions: The metabolic changes in FTLD are promising diagnostic and prognostic biomarkers. Further work is required to replicate these findings, examine longitudinal change, and test their utility in differentiating between FTLD syndromes that are pathologically distinct but phenotypically similar."
90
 Preprint DOI:	10.1101/2020.02.11.20020735
2

"BackgroundA novel coronavirus disease (COVID-19) outbreak due to the severe respiratory syndrome coronavirus (SARS-CoV-2) infection occurred in China in late December 2019. Facemask wearing with proper hand hygiene is considered an effective measure to prevent SARS-CoV-2 transmission, but facemask wearing has become a social concern due to the global facemask shortage. China is the major facemask producer in the world, contributing to 50% of global production. However, a universal facemask wearing policy would put an enormous burden on the facemask supply.MethodsWe performed a policy review concerning facemasks using government websites and mathematical modelling shortage analyses based on data obtained from the National Health Commission (NHC), the Ministry of Industry and Information Technology (MIIT), the Centre for Disease Control and Prevention (CDC), and General Administration of Customs (GAC) of the People's Republic of China. Three scenarios with respect to wearing facemasks were considered: (1) a universal facemask wearing policy implementation in all regions of mainland China; (2) a universal facemask wearing policy implementation only in the epicentre (Hubei province, China); and (3) no implementation of a universal facemask wearing policy..FindingsRegardless of different universal facemask wearing policy scenarios, facemask shortage would occur but eventually end during our prediction period (from 20 Jan 2020 to 30 Jun 2020). The duration of the facemask shortage described in the scenarios of a country-wide universal facemask wearing policy, a universal facemask wearing policy in the epicentre, and no universal facemask wearing policy were 132, seven, and four days, respectively. During the prediction period, the largest daily facemask shortages were predicted to be 589·5, 49·3, and 37·5 million in each of the three scenarios, respectively. . In any scenario, an N95 mask shortage was predicted to occur on 24 January 2020 with a daily facemask shortage of 2·2 million.InterpretationImplementing a universal facemask wearing policy in the whole of China could lead to severe facemask shortage. Without effective public communication, a universal facemask wearing policy could result in societal panic and subsequently, increase the nationwide and worldwide demand for facemasks. These increased demands could cause a facemask shortage for healthcare workers and reduce the effectiveness of outbreak control in the affected regions, eventually leading to a pandemic. To fight novel infectious disease outbreaks, such as COVID-19, governments should monitor domestic facemask supplies and give priority to healthcare workers. . The risk of asymptomatic transmission and facemask shortages should be carefully evaluated before introducing a universal facemask wearing policy in high-risk regions. . Public health measures aimed at improving hand hygiene and effective public communication should be considered along with the facemask policy."	."
91
 Preprint DOI:	10.1101/2020.02.11.20022186
5

Since the first suspected case of coronavirus disease-2019 (COVID-19) on December 1st, 2019, in Wuhan, Hubei Province, China, a total of 40,235 confirmed cases and 909 deaths have been reported in China up to February 10, 2020, evoking fear locally and internationally. Here, based on the publicly available epidemiological data for Hubei, China from January 11 to February 10, 2020, we provide estimates of the main epidemiological parameters. In particular, we provide an estimation of the case fatality and case recovery ratios, along with their 90% confidence intervals as the outbreak evolves. On the basis of a Susceptible-Infectious-Recovered-Dead (SIDR) model, we provide estimations of the basic reproduction number (R0), and the per day infection mortality and recovery rates. By calibrating the parameters of the SIRD model to the reported data, we also attempt to forecast the evolution of the outbreak at the epicenter three weeks ahead, i.e. until February 29. As the number of infected individuals, especially of those with asymptomatic or mild courses, is suspected to be much higher than the official numbers, which can be considered only as a subset of the actual numbers of infected and recovered cases in the total population, we have repeated the calculations under a second scenario that considers twenty times the number of confirmed infected cases and forty times the number of recovered, leaving the number of deaths unchanged.d. Based on the reported data, the expected value of R0 as computed considering the period from the 11th of January until the 18th of January, using the official counts of confirmed cases was found to be ∼4.6, , while the one computed under the second scenario was found to be ∼3.2. Thus, based on the SIRD simulations, the estimated average value of R0 was found to be ∼2.6 based on confirmed cases and ∼2 based on the second scenario. Our forecasting flashes a note of caution for the presently unfolding outbreak in China. Based on the official counts for confirmed cases, the simulations suggest that the cumulative number of infected could reach 180,000 (with a lower bound of 45,000) by February 29. . Regarding the number of deaths, simulations forecast that on the basis of the up to the 10th of February reported data, the death toll might exceed 2,700 (as a lower bound) by February 29. Our analysis further reveals a significant decline of the case fatality ratio from January 26 to which various factors may have contributed, such as the severe control measures taken in Hubei, China (e.g. quarantine and hospitalization of infected individuals), but mainly because of the fact that the actual cumulative numbers of infected and recovered cases in the population most likely are much higher than the reported ones. Thus, in a scenario where we have taken twenty times the confirmed number of infected and forty times the confirmed number of recovered cases, the case fatality ratio is around ∼0.15% in the total population. Importantly, based on this scenario, simulations suggest a slow down of the outbreak in Hubei at the end of February....
92
 Preprint DOI:	10.1101/2020.02.11.944462
1

The outbreak of a novel coronavirus (2019-nCoV) represents a pandemic threat that has been declared a public health emergency of international concern. The CoV spike (S) glycoprotein is a key target for vaccines, therapeutic antibodies, and diagnostics. To facilitate medical countermeasure development, we determined a 3.5-angstrom-resolution cryo–electron microscopy structure of the 2019-nCoV S trimer in the prefusion conformation. The predominant state of the trimer has one of the three receptor-binding domains (RBDs) rotated up in a receptor-accessible conformation. We also provide biophysical and structural evidence that the 2019-nCoV S protein binds angiotensin-converting enzyme 2 (ACE2) with higher affinity than does severe acute respiratory syndrome (SARS)-CoV S. Additionally, we tested several published SARS-CoV RBD-specific monoclonal antibodies and found that they do not have appreciable binding to 2019-nCoV S, suggesting that antibody cross-reactivity may be limited between the two RBDs. The structure of 2019-nCoV S should enable the rapid development and evaluation of medical countermeasures to address the ongoing public health crisis.
93
 Preprint DOI:	10.1101/2020.02.13.20022806
2

A novel coronavirus (SARS-CoV-2) first detected in Wuhan, China, has spread rapidly since December 2019, causing more than 100,000 confirmed infections and 4000 fatalities (as of 10 March 2020). The outbreak has been declared a pandemic by the WHO on Mar 11, 2020. Here, we explore how seasonal variation in transmissibility could modulate a SARS-CoV-2 pandemic. Data from routine diagnostics show a strong and consistent seasonal variation of the four endemic coronaviruses (229E, HKU1, NL63, OC43) and we parameterise our model for SARS-CoV-2 using using using these data. The model allows for many subpopulations of different size with variable parameters. Simulations of different scenarios show that plausible parameters result in a small peak in early 2020 in temperate regions of the Northern Hemisphere and a larger peak in winter 2020/2021. Variation in transmission and migration rates can result in substantial variation in prevalence between regions. While the uncertainty in parameters is large, the scenarios we explore show that transient reductions in the incidence rate might be due to a combination of seasonal variation and infection control efforts but do not necessarily mean the epidemic is contained. Seasonal forcing on SARS-CoV-2 should thus be taken into account in the further monitoring of the global transmission. The likely aggregated effect of seasonal variation, infection control measures, and transmission rate variation is a prolonged pandemic wave with lower prevalence at any given time, thereby providing a window of opportunity for better preparation of health care systems.
94
 Preprint DOI:	10.1101/2020.02.13.945485
1

The ongoing outbreak of viral pneumonia in China and across the world is associated with a new coronavirus, SARS-CoV-21. This outbreak has been tentatively associated with a seafood market in Wuhan, China, where the sale of wild animals may be the source of zoonotic infection2. Although bats are probable reservoir hosts for SARS-CoV-2, the identity of any intermediate host that may have facilitated transfer to humans is unknown. Here we report the identification of SARS-CoV-2-related coronaviruses in Malayan pangolins (Manis javanica) seized in anti-smuggling operations in southern China. Metagenomic sequencing identified pangolin-associated coronaviruses that belong to two sub-lineages of SARS-CoV-2-related coronaviruses, including one that exhibits strong similarity in the receptor-binding domain to SARS-CoV-2. The discovery of multiple lineages of pangolin coronavirus and their similarity to SARS-CoV-2 suggests that pangolins should be considered as possible hosts in the emergence of new coronaviruses and should be removed from wet markets to prevent zoonotic transmission.
95
 Preprint DOI:	10.1101/2020.02.14.20022897
1

The impact of the drastic reduction in travel volume within mainland China in January and February 2020 was quantified with respect to reports of novel coronavirus (COVID-19) infections outside China. Data on confirmed cases diagnosed outside China were analyzed using statistical models to estimate the impact of travel reduction on three epidemiological outcome measures: (i) the number of exported cases, (ii) the probability of a major epidemic, and (iii) the time delay to a major epidemic. From 28 January to 7 February 2020, we estimated that 226 exported cases (95% confidence interval: 86,449) were prevented, corresponding to a 70.4% reduction in incidence compared to the counterfactual scenario. The reduced probability of a major epidemic ranged from 7% to 20% in Japan, which resulted in a median time delay to a major epidemic of two days. Depending on the scenario, the estimated delay may be less than one day. As the delay is small, the decision to control travel volume through restrictions on freedom of movement should be balanced between the resulting estimated epidemiological impact and predicted economic fallout.
96
 Preprint DOI:	10.1101/2020.02.15.20023374
1

Mutations in the sphingomyelin phosphodiesterase 1 (SMPD1) gene were reported to be associated with Parkinson’s disease and dementia with Lewy bodies.. In the current study, we aimed to evaluate the role of SMPD1 variants in isolated rapid eye movement sleep behavior disorder (iRBD). SMPD1 and its untranslated regions were sequenced using targeted next-generation sequencing in 959 iRBD patients and 1287 controls from European descent. Our study reports no statistically significant association of SMPD1 variants and iRBD. It is hence unlikely that SMPD1 plays a major role in iRBD.
98
 Preprint DOI:	10.1101/2020.02.16.20023671
2

"BackgroundThe dynamic changes of lymphocyte subsets and cytokines profiles of patients with novel coronavirus disease (COVID-19) and their correlation with the disease severity remain unclear.MethodsPeripheral blood samples were longitudinally collected from 40 confirmed COVID-19 patients and examined for lymphocyte subsets by flow cytometry and cytokine profiles by specific immunoassays.FindingsOf the 40 COVID-19 patients enrolled, 13 severe cases showed significant and sustained decreases in lymphocyte counts [0·6 (0·6-0·8)] )] but increases in neutrophil counts [4·7 (3·6-5·8)] )] than 27 mild cases [1.1 (0·8-1·4); 2·0 (1·5-2·9)]. Further analysis demonstrated significant decreases in the counts of T cells, especially CD8+ T cells, as well as increases in IL-6, IL-10, IL-2 and IFN-γ levels in the peripheral blood in the severe cases compared to those in the mild cases. T cell counts and cytokine levels in severe COVID-19 patients who survived the disease gradually recovered at later time points to levels that were comparable to those of the mild cases. Moreover, the neutrophil-to-lymphocyte ratio (NLR) (AUC=0·93) ) and neutrophil-to-CD8+ T cell ratio (N8R) (AUC =0·94) ) were identified as powerful prognostic factors affecting the prognosis for severe COVID-19.InterpretationThe degree of lymphopenia and a proinflammatory cytokine storm is higher in severe COVID-19 patients than in mild cases, and is associated with the disease severity. N8R and NLR may serve as a useful prognostic factor for early identification of severe COVID-19 cases.FundingThe National Natural Science Foundation of China, the National Science and Technology Major Project, the Health Commission of Hubei Province, Huazhong University of Science and Technology, and the Medical Faculty of the University of Duisburg-Essen and Stiftung Universitaetsmedizin, Hospital Essen, Germany."
99
 Preprint DOI:	10.1101/2020.02.16.951913

1
 "The new coronavirus (SARS-CoV-2) outbreak from December 2019 in Wuhan, Hubei, China, has been declared a global public health emergency. Angiotensin I converting enzyme 2 (ACE2), is the host receptor by SARS-CoV-2 to infect human cells. Although ACE2 is reported to be expressed in lung, liver, stomach, ileum, kidney and colon, its expressing levels are rather low, especially in the lung. SARS-CoV-2 may use co-receptors/auxiliary proteins as ACE2 partner to facilitate the virus entry. To identify the potential candidates, we explored the single cell gene expression atlas including 119 cell types of 13 human tissues and analyzed the single cell co-expression spectrum of 51 reported RNA virus receptors and 400 other membrane proteins. Consistent with other recent reports, we confirmed that ACE2 was mainly expressed in lung AT2, liver cholangiocyte, colon colonocytes, esophagus keratinocytes, ileum ECs, rectum ECs, stomach epithelial cells, and kidney proximal tubules. Intriguingly, we found that the candidate co-receptors, manifesting the most similar expression patterns with ACE2 across 13 human tissues, are all peptidases, including ANPEP, DPP4 and ENPEP. Among them, ANPEP and DPP4 are the known receptors for human CoVs, suggesting ENPEP as another potential receptor for human CoVs. We also conducted “CellPhoneDB” analysis to understand the cell crosstalk between CoV-targets and their surrounding cells across different tissues. We found that macrophages frequently communicate with the CoVs targets through chemokine and phagocytosis signaling, highlighting the importance of tissue macrophages in immune defense and immune pathogenesis."

101
 Preprint DOI:	10.1101/2020.02.17.952903
1

The outbreak of a novel coronavirus, which was later formally named the severe acute respiratory coronavirus 2 (SARS-CoV-2), has caused a worldwide public health crisis. Previous studies showed that SARS-CoV-2 is highly homologous to SARS-CoV and infects humans through the binding of the spike protein to ACE2. Here, we have systematically studied the molecular mechanisms of human infection with SARS-CoV-2 and SARS-CoV by protein-protein docking and MD simulations. It was found that SARS-CoV-2 binds ACE2 with a higher affinity than SARS-CoV, which may partly explain that SARS-CoV-2 is much more infectious than SARS-CoV. In addition, the spike protein of SARS-CoV-2 has a significantly lower free energy than that of SARS-CoV, suggesting that SARS-CoV-2 is more stable and may survive a higher temperature than SARS-CoV. This provides insights into the evolution of SARS-CoV-2 because SARS-like coronaviruses have originated in bats. Our computation also suggested that the RBD-ACE2 binding for SARS-CoV-2 is much more temperature-sensitive than that for SARS-CoV. Thus, it is expected that SARS-CoV-2 would decrease its infection ability much faster than SARS-CoV when the temperature rises. These findings would be beneficial for the disease prevention and drug/vaccine development of SARS-CoV-2.
103
 Preprint DOI:	10.1101/2020.02.19.20025452
4

We estimate the distribution of serial intervals for 468 confirmed cases of coronavirus disease reported in China as of February 8, 2020. The mean interval was 3.96 days (95% CI 3.53–4.39 days), SD 4.75 days (95% CI 4.46–5.07 days); 12.6% of case reports indicated presymptomatic transmission.
104
 Preprint DOI:	10.1101/2020.02.19.956581
1

The emergence of SARS-CoV-2 has resulted in >90,000 infections and >3,000 deaths. Coronavirus spike (S) glycoproteins promote entry into cells and are the main target of antibodies. We show that SARS-CoV-2 S uses ACE2 to enter cells and that the receptor-binding domains of SARS-CoV-2 S and SARS-CoV S bind with similar affinities to human ACE2, correlating with the efficient spread of SARS-CoV-2 among humans. We found that the SARS-CoV-2 S glycoprotein harbors a furin cleavage site at the boundary between the S1/S2 subunits, which is processed during biogenesis and sets this virus apart from SARS-CoV and SARS-related CoVs. We determined cryo-EM structures of the SARS-CoV-2 S ectodomain trimer, providing a blueprint for the design of vaccines and inhibitors of viral entry. Finally, we demonstrate that SARS-CoV S murine polyclonal antibodies potently inhibited SARS-CoV-2 S mediated entry into cells, indicating that cross-neutralizing antibodies targeting conserved S epitopes can be elicited upon vaccination.
105
 Preprint DOI:	10.1101/2020.02.20.20025619
2

Previous studies have showed clinical characteristics of patients with the 2019 novel coronavirus disease (COVID-19) and the evidence of person-to-person transmission. Limited data are available for asymptomatic infections. This study aims to present the clinical characteristics of 24 cases with asymptomatic infection screened from close contacts and to show the transmission potential of asymptomatic COVID-19 virus carriers. Epidemiological investigations were conducted among all close contacts of COVID-19 patients (or suspected patients) in Nanjing, Jiangsu Province, China, from Jan 28 to Feb 9, 2020, both in clinic and in community. Asymptomatic carriers were laboratory-confirmed positive for the COVID-19 virus by testing the nucleic acid of the pharyngeal swab samples. Their clinical records, laboratory assessments, and chest CT scans were reviewed. As a result, none of the 24 asymptomatic cases presented any obvious symptoms while nucleic acid screening. Five cases (20.8%) developed symptoms (fever, cough, fatigue, etc.) during hospitalization. Twelve (50.0%) cases showed typical CT images of ground-glass chest and 5 (20.8%) presented stripe shadowing in the lungs. The remaining 7 (29.2%) cases showed normal CT image and had no symptoms during hospitalization. These 7 cases were younger (median age: 14.0 years; P=0.012) than the rest. None of the 24 cases developed severe COVID-19 pneumonia or died. The median communicable period, defined as the interval from the first day of positive nucleic acid tests to the first day of continuous negative tests, was 9.5 days (up to 21 days among the 24 asymptomatic cases). Through epidemiological investigation, we observed a typical asymptomatic transmission to the cohabiting family members, which even caused severe COVID-19 pneumonia. Overall, the asymptomatic carriers identified from close contacts were prone to be mildly ill during hospitalization. However, the communicable period could be up to three weeks and the communicated patients could develop severe illness. These results highlighted the importance of close contact tracing and longitudinally surveillance via virus nucleic acid tests. Further isolation recommendation and continuous nucleic acid tests may also be recommended to the patients discharged.
106
 Preprint DOI:	10.1101/2020.02.20.20025866
2

On 5 February 2020, in Yokohama, Japan, a cruise ship hosting 3,711 people underwent a 2-week quarantine after a former passenger was found with COVID-19 post-disembarking. As at 20 February, 634 persons on board tested positive for the causative virus. We conducted statistical modelling to derive the delay-adjusted asymptomatic proportion of infections, along with the infections’ timeline. The estimated asymptomatic proportion was 17.9% ((((95% credible interval (CrI): 15.5–20.2%). Most infections occurred before the quarantine start.
107
 Preprint DOI:	10.1101/2020.02.21.20026187
1

Objectives In December 2019, a novel coronavirus (SARS-CoV-2)-infected pneumonia (COVID-19) occurred in Wuhan, China. Laboratory-based diagnostic tests utilized real-time reverse transcriptase polymerase chain reaction (RT-PCR) on throat samples. This study evaluated the diagnostic value to analyzing throat and sputum samples in order to improve accuracy and detection efficiency. Methods Paired specimens of throat swabs and sputum were obtained from 54 cases, and RNA was extracted and tested for 2019-nCoV (equated with SARS-CoV-2) by the RT-PCR assay. Results The positive rates of 2019-nCoV from sputum specimens and throat swabs were 76.9% and 44.2%, respectively. Sputum specimens showed a significantly higher positive rate than throat swabs in detecting viral nucleic acid using the RT-PCR assay (p = 0.001). Conclusions The detection rates of 2019-nCoV from sputum specimens were significantly higher than those from throat swabs. We suggest that sputum would benefit for the detection of 2019-nCoV in patients who produce sputum. The results can facilitate the selection of specimens and increase the accuracy of diagnosis
108
 Preprint DOI:	10.1101/2020.02.22.20024927
1

Radiologic characteristics of 2019 novel coronavirus (2019-nCoV) infected pneumonia (NCIP) which had not been fully understood are especially important for diagnosing and predicting prognosis. We retrospective studied 27 consecutive patients who were confirmed NCIP, the clinical characteristics and CT image findings were collected, and the association of radiologic findings with mortality of patients was evaluated. 27 patients included 12 men and 15 women, with median age of 60 years (IQR 47–69). 17 patients discharged in recovered condition and 10 patients died in hospital. The median age of mortality group was higher compared to survival group (68 (IQR 63–73) vs 55 (IQR 35–60), P = 0.003). The comorbidity rate in mortality group was significantly higher than in survival group (80% vs 29%, P = 0.018). The predominant CT characteristics consisted of ground glass opacity (67%), bilateral sides involved (86%), both peripheral and central distribution (74%), and lower zone involvement (96%). The median CT score of mortality group was higher compared to survival group (30 (IQR 7–13) vs 12 (IQR 11–43), P = 0.021), with more frequency of consolidation (40% vs 6%, P = 0.047) and air bronchogram (60% vs 12%, P = 0.025). An optimal cutoff value of a CT score of 24.5 had a sensitivity of 85.6% and a specificity of 84.5% for the prediction of mortality. 2019-nCoV was more likely to infect elderly people with chronic comorbidities. CT findings of NCIP were featured by predominant ground glass opacities mixed with consolidations, mainly peripheral or combined peripheral and central distributions, bilateral and lower lung zones being mostly involved. A simple CT scoring method was capable to predict mortality.
109
 Preprint DOI:	10.1101/2020.02.23.20026856
2

"BackgroundRecent studies have focused on initial clinical and epidemiological characteristics of the coronavirus disease 2019 (COVID-19), which is the mainly revealing situation in Wuhan, Hubei.AimThis study aims to reveal more data on the epidemiological and clinical characteristics of COVID-19 patients outside of Wuhan, Zhejiang, China.DesignThis study was a retrospective case series.MethodsEighty-eight cases of laboratory-confirmed and three cases of clinically confirmed COVID-19 were admitted to five hospitals in Zhejiang province, China. Data were collected from 20 January 2020 to 11 February 2020.Results and discussionOf all 91 patients, 88 (96.70%) were laboratory-confirmed COVID-19 with throat swab samples that tested positive for SARS-Cov-2, three (3.30%) cases were clinically diagnosed. The median age of the patients was 50 (36.5–57) years, and female accounted for 59.34%. In this sample, 40 (43.96%) patients had contracted the disease from local cases, 31 (34.07%) patients had been to Wuhan/Hubei, eight (8.79%) patients had contacted with people from Wuhan, and 11 (12.09%) patients were diagnosed after having flown together in the same flight with no passenger that could later be identified as the source of infection. In particular within the city of Ningbo, 60.52% cases can be traced back to an event held in a temple. The most common symptoms were fever (71.43%), cough (60.44%) and fatigue (43.96%). The median of incubation period was 6 (interquartile range 3–8) days and the median time from the first visit to a doctor to the confirmed diagnosis was 1 (1–2) days. According to the chest computed tomography scans, 67.03% cases had bilateral pneumonia.ConclusionsSocial activity cluster, family cluster and flying alongside with persons already infected with COVID-11991199 were how people got infected with COVID-19 in Zhejiang."
110
 Preprint DOI:	10.1101/2020.02.24.20027649
2

An outbreak of COVID-19 developed aboard the Princess Cruises Ship during January–February 2020. Using mathematical modeling and time-series incidence data describing the trajectory of the outbreak among passengers and crew members, we characterize how the transmission potential varied over the course of the outbreak. Our estimate of the mean reproduction number in the confined setting reached values as high as ~11, which is higher than mean estimates reported from community-level transmission dynamics in China and Singapore (approximate range: 1.1–7). ).). Our findings suggest that Rt decreased substantially compared to values during the early phase after the Japanese government implemented an enhanced quarantine control. Most recent estimates of Rt reached values largely below the epidemic threshold, indicating that a secondary outbreak of the novel coronavirus was unlikely to occur aboard the Diamond Princess Ship.
111
 Preprint DOI:	10.1101/2020.02.26.20026971
2

"Background & AimsSome patients with SARS-CoV-2 infection have abnormal liver function. We aimed to clarify the features of COVID-19-related liver damage to provide references for clinical treatment.MethodsWe performed a retrospective, single-center study of 148 consecutive patients with confirmed COVID-19 (73 female, 75 male; mean age, 50 years) at the Shanghai Public Health Clinical Center from January 20 through January 31, 2020. Patient outcomes were followed until February 19, 2020. Patients were analyzed for clinical features, laboratory parameters (including liver function tests), medications, and length of hospital stay. Abnormal liver function was defined as increased levels of alanine and aspartate aminotransferase, gamma glutamyltransferase, alkaline phosphatase, and total bilirubin.ResultsFifty-five patients (37.2%) had abnormal liver function at hospital admission; 14.5% of these patients had high fever (14.5%), compared with 4.3% of patients with normal liver function (P = .027). ). Patients with abnormal liver function were more likely to be male, and had higher levels of procalcitonin and C-reactive protein. There was no statistical difference between groups in medications taken before hospitalization; a significantly higher proportion of patients with abnormal liver function (57.8%) had received lopinavir/ritonavir after admission compared to patients with normal liver function (31.3%). Patients with abnormal liver function had longer mean hospital stays (15.09 ± 4.79 days) than patients with normal liver function (12.76 ± 4.14 days) (P = .021).).ConclusionsMore than one third of patients admitted to the hospital with SARS-CoV-2 infection have abnormal liver function, and this is associated with longer hospital stay. A significantly higher proportion of patients with abnormal liver function had received lopinavir/ritonavir after admission; these drugs should be given with caution."
112
 Preprint DOI:	10.1101/2020.02.26.20028308
1

The Coronavirus Disease 2019 (COVID-19) outbreak is spreading globally. Although COVID-19 has now been declared a pandemic and risk for infection in the United States (US) is currently high, at the time of survey administration the risk of infection in the US was s low. It is important to understand the public perception of risk and trust in sources of information to better inform public health messaging. In this study, we surveyed the adult US population to understand their risk perceptions about the COVID-19 outbreak. We used an online platform to survey 718 adults in the US in early February 2020 using a questionnaire that we developed. Our sample was fairly similar to the general adult US population in terms of age, gender, race, ethnicity and education. We found that 69% of the respondents wanted the scientific/public health leadership (either the CDC Director or NIH Director) to lead the US response to COVID-19 outbreak as compared to 14% who wanted the political leadership (either the president or Congress) to lead the response. Risk perception was low (median score of 5 out of 10) with the respondents trusting health professionals and health officials for information on COVID-19. The majority of respondents were in favor of strict infection prevention policies to control the outbreak. Given our results, the public health/scientific leadership should be at the forefront of the COVID-19 response to promote trust.
113
 Preprint DOI:	10.1101/2020.02.26.964882
3

"A new coronavirus, known as severe acute respiratory syndrome coronavirus 2 (SARS-CoV-2), is the aetiological agent responsible for the 2019–2020 viral pneumonia outbreak of coronavirus disease 2019 (COVID-19)1,2,3,4. Currently, there are no targeted therapeutic agents for the treatment of this disease, and effective treatment options remain very limited. Here we describe the results of a programme that aimed to rapidly discover lead compounds for clinical use, by combining	structure-assisted drug design, virtual drug screening and high-throughput screening. This programme focused on identifying drug leads that target main protease (Mpro) of SARS-CoV-2: Mpro is a key enzyme of coronaviruses and has a pivotal role in mediating viral replication and transcription, making it an attractive drug target for SARS-CoV-25,6. We identified a mechanism-based inhibitor (N3) by computer-aided drug design, and then determined the crystal structure of Mpro of SARS-CoV-2 in complex with this compound. Through a combination of structure-based virtual and high-throughput screening, we assayed more than 10,000 compounds—including approved drugs, drug candidates in clinical trials and other pharmacologically active compounds—as inhibitors of Mpro. Six of these compounds inhibited Mpro, showing half-maximal inhibitory concentration values that ranged from 0.67 to 21.4 µM. One of these compounds (ebselen) also exhibited	promising antiviral activity in cell-based assays. Our results demonstrate the efficacy of our screening strategy, which can lead tto o the rapid discovery of drug leads with clinical potential in response to new infectious diseases for which no specific drugs or vaccines are available."
114
 Preprint DOI:	10.1101/2020.02.27.20028639
2

"BackgroundThe coronavirus disease 2019 (COVID-19) is rapidly spreading in China and more than 30 countries over last two months. COVID-19 has multiple characteristics distinct from other infectious diseases, including high infectivity during incubation, time delay between real dynamics and daily observed number of confirmed cases, and the intervention effects of implemented quarantine and control measures.MethodsWe develop a Susceptible, Un-quanrantined infected, Quarantined infected, Confirmed infected (SUQC) model to characterize the dynamics of COVID-19 and explicitly parameterize the intervention effects of control measures, which is more suitable for analysis than other existing epidemic models.ResultsThe SUQC model is applied to the daily released data of the confirmed infections to analyze the outbreak of COVID-19 in Wuhan, Hubei (excluding Wuhan), China (excluding Hubei) and four first-tier cities of China. We found that, before January 30, 2020, all these regions except Beijing had a reproductive number R > 1, and after January 30, all regions had a reproductive number R < 1, indicating that the quarantine and control measures are effective in preventing the spread of COVID-19. The confirmation rate of Wuhan estimated by our model is 0.0643, substantially lower than that of Hubei excluding Wuhan (0.1914), and that of China excluding Hubei (0.2189), but it jumps to 0.3229 after February 12 when clinical evidence was adopted in new diagnosis guidelines. The number of unquarantined infected cases in Wuhan on February 12, 2020 is estimated to be 3,509 and declines to 334 on February 21, 2020. After fitting the model with data as of February 21, 2020, we predict that the end time of COVID-19 in Wuhan and Hubei is around late March, around mid March for China excluding Hubei, and before early March 2020 for the four tier-one cities. A total of 80,511 individuals are estimated to be infected in China, among which 49,510 are from Wuhan, 17,679 from Hubei (excluding Wuhan), and the rest 13,322 from other regions of China (excluding Hubei). Note that the estimates are from a deterministic ODE model and should be interpreted with some uncertainty....ConclusionsWe suggest that rigorous quarantine and control measures should be kept before early March in Beijing, Shanghai, Guangzhou and Shenzhen, and before late March in Hubei. The model can also be useful to predict the trend of epidemic and provide quantitative guide for other countries at high risk of outbreak, such as South Korea, Japan, Italy and Iran."
115
 Preprint DOI:	10.1101/2020.02.27.20028829
2

"ObjectivesSince the first case of 2019 novel coronavirus (COVID-19) identified on Jan 20, 2020, in South Korea, the number of cases rapidly increased, resulting in 6284 cases including 42 deaths as of Mar 6, 2020. To examine the growth rate of the outbreak, we present the first study to report the reproduction number of COVID-19 in South Korea.MethodsThe daily confirmed cases of COVID-19 in South Korea were extracted from publicly available sources. By using the empirical reporting delay distribution and simulating the generalized growth model, we estimated the effective reproduction number based on the discretized probability distribution of the generation interval.ResultsWe identified four major clusters and estimated the reproduction number at 1.5 (95% CI: 1.4–1.6). In addition, the intrinsic growth rate was estimated at 0.6 (95% CI: 0.6, 0.7), and the scaling of growth parameter was estimated at 0.8 (95% CI: 0.7, 0.8),),),),),),),), indicating sub-exponential growth dynamics of COVID-19. . The crude case fatality rate is higher among males (1.1%) compared to females (0.4%)%)%)%) and increases with older age.ConclusionsOur results indicate an early sustained transmission of COVID-19 in South Korea and support the implementation of social distancing measures to rapidly control the outbreak."
116
 Preprint DOI:	10.1101/2020.02.27.967760
3

The new type of pneumonia caused by the SARS-CoV-2 (Severe acute respiratory syndrome coronavirus 2) has been declared as a global public health concern by WHO. As of April 3, 2020, more than 1,000,000 human infections have been diagnosed around the world, which exhibited apparent person-to-person transmission characteristics of this virus. The capacity of vertical transmission in SARS-CoV-2 remains controversial recently. Angiotensin-converting enzyme 2 (ACE2) is now confirmed as the receptor of SARS-CoV-2 and plays essential roles in human infection and transmission. In present study, we collected the online available single-cell RNA sequencing (scRNA-seq) data to evaluate the cell specific expression of ACE2 in maternal-fetal interface as well as in multiple fetal organs. Our results revealed that ACE2 was highly expressed in maternal-fetal interface cells including stromal cells and perivascular cells of decidua, and cytotrophoblast and syncytiotrophoblast in placenta. Meanwhile, ACE2 was also expressed in specific cell types of human fetal heart, liver and lung, but not in kidney. And in a study containing series fetal and post-natal mouse lung, we observed ACE2 was dynamically changed over the time, and ACE2 was extremely high in neonatal mice at post-natal day 1~3. In summary, this study revealed that the SARS-CoV-2 receptor was widely spread in specific cell types of maternal-fetal interface and fetal organs. And thus, both the vertical transmission and the placenta dysfunction/abortion caused by SARS-CoV-2 need to be further carefully investigated in clinical practice.
117
 Preprint DOI:	10.1101/2020.02.29.20029322
1

Since December 2019, more than 79,000 people have been diagnosed with infection of the Corona Virus Disease 2019 (COVID-19). A large number of medical staff was sent to Wuhan city and Hubei province to aid COVID-19 control. Psychological stress, especially vicarious traumatization caused by the COVID-19 pandemic, should not be ignored. To address this concern, the study employed a total of 214 general public and 526 nurses (i.e., 234 front-line nurses and 292 non-front-line nurses) to evaluate vicarious traumatization scores via a mobile app-based questionnaire. Front-line nurses are engaged in the process of providing care for patients with COVID-19. The results showed that the vicarious traumatization scores for front-line nurses including scores for physiological and psychological responses, were significantly lower than those of non-front-line nurses (P < 0.001). Interestingly, the vicarious traumatization scores of the general public were significantly higher than those of the front-line nurses (P < 0.001); however, no statistical difference was observed compared to the scores of non-front-line nurses (P > 0.05).). Therefore, increased attention should be paid to the psychological problems of the medical staff, especially non-front-line nurses, and general public under the situation of the spread and control of COVID-19. Early strategies that aim to prevent and treat vicarious traumatization in medical staff and general public are extremely necessary.
118
 Preprint DOI:	10.1101/2020.02.29.20029462
1

"BackgroundAssessment of possible infection with SARS‐CoV‐2, the novel coronavirus responsible for COVID‐19 illness, has been a major activity of infection services since the first reports of cases in December 2019.ObjectivesWe report a series of 68 patients assessed at a Regional Infection Unit in the UK.MethodsBetween 29 January 2020 and 24 February 2020, demographic, clinical, epidemiological and laboratory data were collected. We compared clinical features between patients not requiring admission for clinical reasons or antimicrobials with those assessed as needing either admission or antimicrobial treatment.ResultsPatients assessed were aged from 0 to 76 years; 36/68 were female. . Peaks of clinical assessments coincided with updates to the case definition for suspected COVID‐19. Microbiological diagnoses included SARS‐CoV‐2, mycoplasma pneumonia, influenza A, non‐SARS/MERS coronaviruses and rhinovirus/enterovirus. Nine of sixty‐eight received antimicrobials, 15/68 were admitted, 5 due to inability to self‐isolate. Patients requiring admission on clinical grounds or antimicrobials (14/68) were more likely to have fever or raised respiratory rate compared to those not requiring admission or antimicrobials.ConclusionsThe majority of patients had mild illness, which did not require clinical intervention. This finding supports a community testing approach, supported by clinicians able to review more unwell patients. Extensions of the epidemiological criteria for the case definition of suspected COVID‐19 lead to increased screening intensity; strategies must be in place to accommodate this in time for forthcoming changes as the epidemic develops."	."
119
 Preprint DOI:	10.1101/2020.02.29.965418
2

In this work, we use a within-host viral dynamic model to describe the SARS-CoV-2 kinetics in the host. Chest radiograph score data are used to estimate the parameters of that model. Our result shows that the basic reproductive number of SARS-CoV-2 in host growth is around 3.79. Using the same method we also estimate the basic reproductive number of MERS virus is 8.16 which is higher than SARS-CoV-2. The PRCC method is used to analyze the sensitivities of model parameters. Moreover, the drug effects on virus growth and immunity effect of patients are also implemented to analyze the model.
121
 Preprint DOI:	10.1101/2020.03.02.20026708
1

The ongoing coronavirus disease 2019 (COVID-19) outbreak expanded rapidly throughout China. Major behavioral, clinical, and state interventions were undertaken to mitigate the epidemic and prevent the persistence of the virus in human populations in China and worldwide. It remains unclear how these unprecedented interventions, including travel restrictions, affected COVID-19 spread in China. We used real-time mobility data from Wuhan and detailed case data including travel history to elucidate the role of case importation in transmission in cities across China and to ascertain the impact of control measures. Early on, the spatial distribution of COVID-19 cases in China was explained well by human mobility data. After the implementation of control measures, this correlation dropped and growth rates became negative in most locations, although shifts in the demographics of reported cases were still indicative of local chains of transmission outside of Wuhan. This study shows that the drastic control measures implemented in China substantially mitigated the spread of COVID-19.
122
 Preprint DOI:	10.1101/2020.03.02.20030189
1

"BackgroundThe novel coronavirus SARS-CoV-2 is a newly emerging virus. The antibody response in infected patient remains largely unknown, and the clinical values of antibody testing have not been fully demonstrated.MethodsA total of 173 patients with SARS-CoV-2 infection were enrolled. Their serial plasma samples (n=535) collected during the hospitalization were tested for total antibodies (Ab), IgM and IgG against SARS-CoV-2. The dynamics of antibodies with the disease progress was analyzed.ResultsAmong 173 patients, the seroconversion rate for Ab, IgM and IgG was 93.1%, 82.7% and 64.7%, respectively. The reason for the negative antibody findings in 12 patients might due to the lack of blood samples at the later stage of illness. The median seroconversion time for Ab, IgM and then IgG were day-11, day-12 and day-14, separately. The presence of antibodies was <40% among patients within 1-week since onset, and rapidly increased to 100.0% (Ab), 94.3% (IgM) and 79.8% (IgG) since day-15 after onset. In contrast, RNA detectability decreased from 66.7% (58/87) in samples collected before day-7 to 45.5% (25/55) during day 15-39. Combining RNA and antibody detections significantly improved the sensitivity of pathogenic diagnosis for COVID-19 (p<0.001), even in early phase of 1-week since onset (p=0.007). Moreover, a higher titer of Ab was independently associated with a worse clinical classification (p=0.006).ConclusionsThe antibody detection offers vital clinical information during the course of SARS-CoV-2 infection. The findings provide strong empirical support for the routine application of serological testing in the diagnosis and management of COVID-19 patients."
123
 Preprint DOI:	10.1101/2020.03.02.20030320
1

As of March 1, 2020, Iran had reported 987 novel coronavirus disease (COVID-19) cases, including 54 associated deaths. At least six neighboring countries (Bahrain, Iraq, Kuwait, Oman, Afghanistan, and Pakistan) had reported imported COVID-19 cases from Iran. In this study, air travel data and the numbers of cases from Iran imported into other Middle Eastern countries were used to estimate the number of COVID-19 cases in Iran. It was estimated that the total number of cases in Iran was 16 533 (95% confidence interval: 5925–35 538) by February 25, 2020, before the UAE and other Gulf Cooperation Council countries suspended inbound and outbound flights from Iran.
124
 Preprint DOI:	10.1101/2020.03.02.968388
1

Severe Acute Respiratory Syndrome coronavirus 2 (SARS‐CoV‐2) is rapidly spreading around the world. There is no existing vaccine or proven drug to prevent infections and stop virus proliferation. Although this virus is similar to human and animal SARS‐CoVs and Middle East Respiratory Syndrome coronavirus (MERS‐CoVs), the detailed information about SARS‐CoV‐2 proteins structures and functions is urgently needed to rapidly develop effective vaccines, antibodies, and antivirals. We applied high‐throughput protein production and structure determination pipeline at the Center for Structural Genomics of Infectious Diseases to produce SARS‐CoV‐2 proteins and structures. Here we report two high‐resolution crystal structures of endoribonuclease Nsp15/NendoU. We compare these structures with previously reported homologs from SARS and MERS coronaviruses.
125
 Preprint DOI:	10.1101/2020.03.05.20031773
2

Adjusting for delay from confirmation to death, we estimated case and infection fatality ratios (CFR, IFR) for coronavirus disease (COVID-19) on the Diamond Princess ship as 2.6% (95% confidence interval (CI): 0.89–6.7) and 1.3% (95% CI: 0.38–3.6), ), respectively. Comparing deaths on board with expected deaths based on naive CFR estimates from China, we estimated CFR and IFR in China to be 1.2% (95% CI: 0.3–2.7) and 0.6% (95% CI: 0.2–1.3), respectively.
126
 Preprint DOI:	10.1101/2020.03.06.20032193
1

"Temporal dynamics of certain human microbiotas have been described in longitudinal studies; variability often relates to modifiable factors or behaviors. Early studies of the urinary microbiota preferentially used samples obtained by transurethral catheterization to minimize vulvovaginal microbial contributions. Whereas voided specimens are preferred for longitudinal studies, the few studies that reported longitudinal data were limited to women with lower urinary tract (LUT) symptoms, due to ease of accessing a clinical population for sampling and the impracticality and risk of collecting repeated catheterized urine specimens in a nonclinical population. Here, we studied the microbiota of the LUT of nonsymptomatic, premenopausal women using midstream voided urine (MSU) specimens to investigate relationships between microbial dynamics and personal factors. Using 16S rRNA gene sequencing and a metaculturomics method called expanded quantitative urine culture (EQUC), we characterized the microbiotas of MSU and periurethral swab specimens collected daily for approximately 3 months from a small cohort of adult women. Participants were screened for eligibility, including the ability to self-collect paired urogenital specimens prior to enrollment. In this population, we found that measures of microbial dynamics related to specific participant-reported factors, particularly menstruation and vaginal intercourse. Further investigation of the trends revealed differences in the composition and diversity of LUT microbiotas within and across participants. These data, in combination with previous studies showing relationships between the LUT microbiota and LUT symptoms, suggest that personal factors relating to the genitourinary system may be an important consideration in the etiology, prevention, and/or treatment of LUT disorders.IMPORTANCE Following the discovery of the collective human urinary microbiota, important knowledge gaps remain, including the stability and variability of this microbial niche over time. Initial urinary studies preferentially utilized samples obtained by transurethral catheterization to minimize contributions from vulvovaginal microbes. However, catheterization has the potential to alter the urinary microbiota; therefore, voided specimens are preferred for longitudinal studies. In this report, we describe microbial findings obtained by daily assessment over 3 months in a small cohort of adult women. We found that, similarly to vaginal microbiotas, lower urinary tract (LUT) microbiotas are dynamic, with changes relating to several factors, particularly menstruation and vaginal intercourse. Our study results show that LUT microbiotas are both dynamic and resilient. They also offer novel opportunities to target LUT microbiotas by preventative or therapeutic means, through risk and/or protective factor modification."."."."
127
 Preprint DOI:	10.1101/2020.03.06.977876
2

"BackgroundThe outbreak of coronavirus disease (COVID-19) in China caused by the SARS-CoV-2 virus continually lead to worldwide human infections and deaths. It is currently no specific viral protein targeted therapeutics. Viral nucleocapsid protein is a potential antiviral drug target, serving multiple critical functions during the viral life cycle. However, the structural information of the SARS-CoV-2 nucleocapsid protein remains unclear.MethodsTo obtain the structural information of the SARS-CoV-2 nucleocapsid protein, we have determined the 2.7 Å crystal structure of the N-terminal RNA binding domain of SARS-CoV-2 nucleocapsid protein using X-ray crystallography technology. To explored the interaction mechanism, we complemented functional studies by in vitro surface plasmon resonance analysis and biolayer interferometry assays....ResultsAlthough the overall structure is similar to other reported coronavirus nucleocapsid protein N-terminal domain, the surface electrostatic potential characteristics between them are distinct. Further comparison with mild virus type HCoV-OC43 equivalent domain demonstrates a unique potential RNA binding pocket alongside the β-sheet core.ConclusionsOur data provide several atomic resolution features of the SARS-CoV-2 nucleocapsid protein N-terminal domain, guiding the design of novel antiviral agents specific targeting to SARS-CoV-2."
128
 Preprint DOI:	10.1101/2020.03.07.20031575
1

"BackgroundThe outbreak of coronavirus disease 2019 (COVID-19) caused by severe acute respiratory syndrome coronavirus 2 (SARS-CoV-2) in China has been declared a public health emergency of international concern. The cardiac injury is a common condition among the hospitalized patients with COVID-19. However, whether N terminal pro B type natriuretic peptide (NT-proBNP) predicted outcome of severe COVID-19 patients was unknown.MethodsThe study initially enrolled 102 patients with severe COVID-19 from a continuous sample. After screening out the ineligible cases, 54 patients were analyzed in this study. The primary outcome was in-hospital death defined as the case fatality rate. Research information and following-up data were obtained from their medical records.ResultsThe best cut-off value of NT-proBNP for predicting in-hospital death was 88.64 pg/mL with the sensitivity for 100% and the specificity for 66.67%. %.%.%.%. Patients with high NT-proBNP values (> 88.64 pg/mL) had a significantly increased risk of death during the days of following-up compared with those with low values (≤88.64 pg/mL). After adjustment for potential risk factors, NT-proBNP was independently correlated with in-hospital death.ConclusionNT-proBNP might be an independent risk factor for in-hospital death in patients with severe COVID-19."
129
 Preprint DOI:	10.1101/2020.03.08.20032946
2

"INTRODUCTIONCoronavirus disease 2019 (COVID-19), caused by severe acute respiratory syndrome–coronavirus 2 (SARS-CoV-2), has clear potential for a long-lasting global pandemic, high fatality rates, and incapacitated health systems. Until vaccines are widely available, the only available infection prevention approaches are case isolation, contact tracing and quarantine, physical distancing, decontamination, and hygiene measures. To implement the right measures at the right time, it is of crucial importance to understand the routes and timings of transmission.RATIONALEWe used key parameters of epidemic spread to estimate the contribution of different transmission routes with a renewal equation formulation, and analytically determined the speed and scale for effective identification and contact tracing required to stop the epidemic.RESULTSWe developed a mathematical model for infectiousness to estimate the basic reproductive number R0 and to quantify the contribution of different transmission routes. To parameterize the model, we analyzed 40 well-characterized source-recipient pairs and estimated the distribution of generation times (time from infection to onward transmission).).).). The distribution had a median of 5.0 days and standard deviation of 1.9 days. . We used published parameters for the incubation time distribution (median 5.2 days) and the epidemic doubling time (5.0 days) from the early epidemic data in China.The model estimated R0 = 2.0 in the early stages of the epidemic in China. The contributions to R0 included 46% from presymptomatic individuals (before showing symptoms), 38% from symptomatic individuals, 10% from asymptomatic individuals (who never show symptoms), and 6% from environmentally mediated transmission via contamination. Results on the last two routes are speculative. According to these estimates, presymptomatic transmissions alone are almost sufficient to sustain epidemic growth.To estimate the requirements for successful contact tracing, we determined the combination of two key parameters needed to reduce R0 to less than 1: the proportion of cases who need to be isolated, and the proportion of their contacts who need to be quarantined. For a 3-day delay in notification assumed for manual contact tracing, no parameter combination leads to epidemic control. . Immediate notification through a contact-tracing mobile phone app could, however, be sufficient to stop the epidemic if used by a sufficiently high proportion of the population.We propose an app, based on existing technology, that allows instant contact tracing. Proximity events between two phones running the app are recorded. Upon an individual’s COVID-19 diagnosis, contacts are instantly, automatically, and anonymously notified of their risk and asked to self-isolate. Practical and logistical factors (e.g., uptake, coverage, R0 in a given population) will determine whether an app is sufficient to control viral spread on its own, or whether additional measures to reduce R0 (e.g., physical distancing) are required. The performance of the app in scenarios with higher values of R0 can be explored at https://bdi-pathogens.shinyapps.io/covid-19-transmission-routes/.CONCLUSIONGiven the infectiousness of SARS-CoV-2 and the high proportion of transmissions from presymptomatic individuals, controlling the epidemic by manual contact tracing is infeasible. The use of a contact-tracing app that builds a memory of proximity contacts and immediately notifies contacts of positive cases would be sufficient to stop the epidemic if used by enough people, in particular when combined with other measures such as physical distancing. An intervention of this kind raises ethical questions regarding access, transparency, the protection and use of personal data, and the sharing of knowledge with other countries. Careful oversight by an inclusive advisory body is required..""..""
131
 Preprint DOI:	10.1101/2020.03.09.983064
2

The coronavirus disease 2019 (COVID-19) pandemic now has >2,000,000 confirmed cases worldwide. COVID-19 is currently diagnosed using quantitative RT-PCR methods, but the capacity of quantitative RT-PCR methods is limited by their requirement of high-level facilities and instruments. We developed and evaluated reverse transcription loop-mediated isothermal amplification (RT-LAMP) assays to detect genomic RNA of severe acute respiratory syndrome coronavirus 2 (SARS-CoV-2), the causative virus of COVID-19. RT-LAMP assays reported in this study can detect as low as 100 copies of SARS-CoV-2 RNA. Cross-reactivity of RT-LAMP assays to other human coronaviruses was not observed. A colorimetric detection method was adapted for this RT-LAMP assay to enable higher throughput.
132
 Preprint DOI:	10.1101/2020.03.09.983247
1

The recent outbreak of coronavirus disease (COVID-19) caused by SARS-CoV-2 infection in Wuhan, China has posed a serious threat to global public health. To develop specific anti-coronavirus therapeutics and prophylactics, the molecular mechanism that underlies viral infection must first be defined. Therefore, we herein established a SARS-CoV-2 spike (S) protein-mediated cell–cell fusion assay and found that SARS-CoV-2 showed a superior plasma membrane fusion capacity compared to that of SARS-CoV. We solved the X-ray crystal structure of six-helical bundle (6-HB) core of the HR1 and HR2 domains in the SARS-CoV-2 S protein S2 subunit, revealing that several mutated amino acid residues in the HR1 domain may be associated with enhanced interactions with the HR2 domain. We previously developed a pan-coronavirus fusion inhibitor, EK1, which targeted the HR1 domain and could inhibit infection by divergent human coronaviruses tested, including SARS-CoV and MERS-CoV. Here we generated a series of lipopeptides derived from EK1 and found that EK1C4 was the most potent fusion inhibitor against SARS-CoV-2 S protein-mediated membrane fusion and pseudovirus infection with IC50s of 1.3 and 15.8 nM, about 241- and 149-fold more potent than the original EK1 peptide, respectively. EK1C4 was also highly effective against membrane fusion and infection of other human coronavirus pseudoviruses tested, including SARS-CoV and MERS-CoV, as well as SARSr-CoVs, and potently inhibited the replication of 5 live human coronaviruses examined, including SARS-CoV-2. Intranasal application of EK1C4 before or after challenge with HCoV-OC43 protected mice from infection, suggesting that EK1C4 could be used for prevention and treatment of infection by the currently circulating SARS-CoV-2 and other emerging SARSr-CoVs.
133
 Preprint DOI:	10.1101/2020.03.09.984856
1

Tilorone is a 50-year-old synthetic small-molecule compound with antiviral activity that is proposed to induce interferon after oral administration. This drug is used as a broad-spectrum antiviral in several countries of the Russian Federation. We have recently described activity in vitro and in vivo against the Ebola virus. After a broad screening of additional viruses, we now describe in vitro activity against Chikungunya
134
 Preprint DOI:	10.1101/2020.03.10.20033605
1

"BackgroundThe ongoing epidemics of coronavirus disease 2019 (COVID-19) have caused serious concerns about its potential adverse effects on pregnancy. There are limited data on maternal and neonatal outcomes of pregnant women with COVID-19 pneumonia.MethodsWe conducted a case-control study to compare clinical characteristics, maternal and neonatal outcomes of pregnant women with and without COVID-19 pneumonia.ResultsDuring January 24 to February 29, 2020, there were sixteen pregnant women with confirmed COVID-19 pneumonia and eighteen suspected cases who were admitted to labor in the third trimester. Two had vaginal delivery and the rest took cesarean section. Few patients presented respiratory symptoms (fever and cough) on admission, but most had typical chest CT images of COVID-19 pneumonia. Compared to the controls, COVID-19 pneumonia patients had lower counts of white blood cells (WBC), neutrophils, C-reactive protein (CRP), and alanine aminotransferase (ALT) on admission. Increased levels of WBC, neutrophils, eosinophils, and CRP were found in postpartum blood tests of pneumonia patients. There were three (18.8%) and three (16.7%) %) of the mothers with confirmed or suspected COVID-19 pneumonia had preterm delivery due to maternal complications, which were significantly higher than the control group. None experienced respiratory failure during hospital stay. COVID-19 infection was not found in the newborns and none developed severe neonatal complications.ConclusionSevere maternal and neonatal complications were not observed in pregnant women with COVID-19 pneumonia who had vaginal delivery or caesarean section. Mild respiratory symptoms of pregnant women with COVID-19 pneumonia highlight the need of effective screening on admission."
135
 Preprint DOI:	10.1101/2020.03.11.20034314
1

We model the COVID-19 coronavirus epidemic in China. We use early reported case data to predict the cumulative number of reported cases to a final size. The key features of our model are the timing of implementation of major public policies restricting social movement, the identification and isolation of unreported cases, and the impact of asymptomatic infectious cases.
136
 Preprint DOI:	10.1101/2020.03.13.20034496
2

We assess the health and wellbeing of normal adults living and working after one month of confinement to contain the COVID-19 outbreak in China. On Feb 20–21, 2020, we surveyed 369 adults in 64 cities in China that varied in their rates of confirmed coronavirus cases on their health conditions, distress and life satisfaction. 27% of the participants worked at the office, 38% resorted to working from home, and 25% stopped working due to the outbreak. Those who stopped working reported worse mental and physical health conditions as well as distress. The severity of COVID-19 in an individual's home city predicts their life satisfaction, and this relationship is contingent upon individuals’ existing chronic health issues and their hours of exercise. Our evidence supports the need to pay attention to the health of people who were not infected by the virus, especially for people who stopped working during the outbreak. Our results highlight that physically active people might be more susceptible to wellbeing issues during the lockdown. Policymakers who are considering introducing restrictive measures to contain COVID-19 may benefit from understanding such health and wellbeing implications.
137
 Preprint DOI:	10.1101/2020.03.13.20035410
1

"•COVID-19 mortality rates outside Hubei and Wuhan are nearly constant. •COVID-19 recovery rates in Wuhan and Hubei grow exponentially. •Over 40,000 aided health workers help Hubei to effectively treat patients. •Newly supplied beds allow over 38,000 patients in Wuhan to be treated in hospitals."
138
 Preprint DOI:	10.1101/2020.03.13.20035428
1

An outbreak of new coronavirus SARS-CoV-2 was occurred in Wuhan, China and rapidly spread to other cities and nations. The standard diagnostic approach that widely adopted in the clinic is nucleic acid detection by real-time RT-PCR. However, the false-negative rate of the technique is unneglectable and serological methods are urgently warranted. Here, we presented the colloidal gold-based immunochromatographic (ICG) strip targeting viral IgM or IgG antibody and compared it with real-time RT-PCR. The sensitivity of ICG assay with IgM and IgG combinatorial detection in nucleic acid confirmed cases were 11.1%, 92.9% and 96.8% at the early stage (1–7 days after onset), intermediate stage (8–14 days after onset), and late stage (more than 15 days), respectively. The ICG detection capacity in nucleic acid-negative suspected cases was 43.6%. In addition, the concordance of whole blood samples and plasma showed Cohen's kappa value of 0.93, which represented the almost perfect agreement between two types of samples. In conclusion, serological ICG strip assay in detecting SARS-CoV-2 infection is both sensitive and consistent, which is considered as an excellent supplementary approach in clinical application.
139
 Preprint DOI:	10.1101/2020.03.13.20035485
2

We constructed a simple Susceptible–Exposed–Infectious–Removed model of the spread of COVID-19. The model is parametrised only by the average incubation period, t, and two rate parameters: contact rate, ß, and exclusion rate, <U+03B3>. The rates depend on nontherapeutic interventions and determine the basic reproduction number, R0 = ß/<U+03B3>, and, together with t, the daily multiplication coefficient in the early exponential phase, <U+03B8>. Initial R0 determines the reduction of ß required to contain the spread of the epidemic. We demonstrate that introduction of a a cascade of multiple exposed states enables the model to reproduce the distributions of the incubation period and the serialinterval reported by epidemiologists. Using the model, we consider a hypothetical scenario in which ß is modulated solely by anticipated changes of social behaviours: first, ß decreases in response to a surge of daily new cases, pressuring people to self-isolate, and then, over longer time scale, ß increases as people gradually accept the risk. In this scenario, initial abrupt epidemic spread is followed by a plateau and slow regression, which, although economically and socially devastating, grants time to develop and deploy vaccine or atleast limit daily cases to a manageable number..
140
 Preprint DOI:	10.1101/2020.03.13.20035568
2

"Background: Given the extensive time needed to conduct a nationally representative household survey and the commonly low response rate of phone surveys, rapid online surveys may be a promising method to assess and track knowledge and perceptions among the general public during fast-moving infectious disease outbreaks.Objective: This study aimed to apply rapid online surveying to determine knowledge and perceptions of coronavirus disease 2019 (COVID-19) among the general public in the United States and the United Kingdom.Methods: An online questionnaire was administered to 3000 adults residing in the United States and 3000 adults residing in the United Kingdom who had registered with Prolific Academic to participate in online research. Prolific Academic established strata by age (18-27, 28-37, 38-47, 48-57, or ≥58 years), sex (male or female), and ethnicity (white, black or African American, Asian or Asian Indian, mixed, or “other”), as well as all permutations of these strata. The number of participants who could enroll in each of these strata was calculated to reflect the distribution in the US and UK general population. Enrollment into the survey within each stratum was on a first-come, first-served basis. Participants completed the questionnaire between February 23 and March 2, 2020.Results: A total of 2986 and 2988 adults residing in the United States and the United Kingdom, respectively, completed the questionnaire. Of those, 64.4% (1924/2986) of US participants and 51.5% (1540/2988) of UK participants had a tertiary education degree, 67.5% (2015/2986) of US participants had a total household income between US $20,000 and US $99,999, and 74.4% (2223/2988) of UK participants had a total household income between £15,000 and £74,999. US and UK participants’ median estimate for the probability of a fatal disease course among those infected with severe acute respiratory syndrome coronavirus 2 (SARS-CoV-2) was 5.0% (IQR 2.0%-15.0%) and 3.0% (IQR 2.0%-10.0%), respectively. Participants generally had good knowledge of the main mode of disease transmission and common symptoms of COVID-19. However, a substantial proportion of participants had misconceptions about how to prevent an infection and the recommended care-seeking behavior. For instance, 37.8% (95% CI 36.1%-39.6%) of US participants and 29.7% (95% CI 28.1%-31.4%) of UKUKUKUKUKUK participants thought that wearing a common surgical mask was “highly effective” in protecting them from acquiring COVID-19, and 25.6% (95% CI 24.1%-27.2%) of US participants and 29.6% (95% CI 28.0%-31.3%) of UK participants thought it was prudent to refrain from eating at Chinese restaurants. Around half (53.8%, 95% CI 52.1%-55.6%) of US participants and 39.1% (95% CI 37.4%-40.9%) of UK participants thought that children were at an especially high risk of death when infected with SARS-CoV-2.Conclusions: The distribution of participants by total household income and education followed approximately that of the US and UK general population. The findings from this online survey could guide information campaigns by public health authorities, clinicians, and the media. More broadly, rapid online surveys could be an important tool in tracking the public’s knowledge and misperceptions during rapidly moving infectious disease outbreaks."
141
 Preprint DOI:	10.1101/2020.03.13.991570
1

The outbreak of coronavirus disease 2019 (COVID-19) caused by severe acute respiratory syndrome–coronavirus 2 (SARS-CoV-2) has now become a pandemic, but there is currently very little understanding of the antigenicity of the virus. We therefore determined the crystal structure of CR3022, a neutralizing antibody previously isolated from a convalescent SARS patient, in complex with the receptor binding domain (RBD) of the SARS-CoV-2 spike (S) protein at 3.1-angstrom resolution.... CR3022 targets a highly conserved epitope, distal from the receptor binding site, that enables cross-reactive binding between SARS-CoV-2 and SARS-CoV. Structural modeling further demonstrates that the binding epitope can only be accessed by CR3022 when at least two RBDs on the trimeric S protein are in the “up” conformation and slightly rotated. These results provide molecular insights into antibody recognition of SARS-CoV-2.
142
 Preprint DOI:	10.1101/2020.03.14.20036202
1

Based on the official data modeling, this paper studies the transmission process of the Corona Virus Disease 2019 (COVID-19). The error between the model and the official data curve is quite small. . At the same time, it realized forward prediction and backward inference of the epidemic situation, and the relevant analysis help relevant countries to make decisions.
143
 Preprint DOI:	10.1101/2020.03.15.20032870
1

This report describes the isolation, molecular characterization, and phylogenetic analysis of the first three complete genomes of severe acute respiratory syndrome coronavirus 2 (SARS‐CoV‐2) isolated from three patients involved in the first outbreak of COVID‐19 in Lombardy, Italy. Early molecular epidemiological tracing suggests that SARS‐CoV‐2 was present in Italy weeks before the first reported cases of infection.
145
 Preprint DOI:	10.1101/2020.03.15.20036293
2

Governments around the world must rapidly mobilize and make difficult policy decisions to mitigate the coronavirus disease 2019 (COVID-19) pandemic. Because deaths have been concentrated at older ages, we highlight the important role of demography, particularly, how the age structure of a population may help explain differences in fatality rates across countries and how transmission unfolds. We examine the role of age structure in deaths thus far in Italy and South Korea and illustrate how the pandemic could unfold in populations with similar population sizes but different age structures, showing a dramatically higher burden of mortality in countries with older versus younger populations. This powerful interaction of demography and current age-specific mortality for COVID-19 suggests that social distancing and other policies to slow transmission should consider the age composition of local and national contexts as well as intergenerational interactions. We also call for countries to provide case and fatality data disaggregated by age and sex to improve real-time targeted forecasting of hospitalization and critical care needs....
146
 Preprint DOI:	10.1101/2020.03.15.20036350
1

Between January 24th and March 10th, a total of 2,370 individuals had contact with the first 30 cases of COVID-19. There were 13 individuals who contracted COVID-19 resulting in a secondary attack rate of 0.55% (95% CI 0.31–0.96). There were 119 household contacts, of which 9 individuals developed COVID-19 resulting in a secondary attack rate of 7.56% (95% CI 3.7–14.26).
147
 Preprint DOI:	10.1101/2020.03.15.20036368
1

We report the first 7,755 patients with confirmed COVID-19 in Korea as of March 12th, 2020. A total of 66 deaths have been recorded, giving a case fatality proportion of 0.9%. Older people, and those with comorbidities were at a higher risk of a fatal outcome. The highest number of cases of COVID-19 were in Daegu, followed by Gyeongbuk.. This summary may help to understand the disease dynamics in the early phase of the COVID-19 outbreaks, and may therefore, guide future public health measures.
149
 Preprint DOI:	10.1101/2020.03.15.20036707
2

We report temporal patterns of viral shedding in 94 patients with laboratory-confirmed COVID-19 and modeled COVID-19 infectiousness profiles from a separate sample of 77 infector–infectee transmission pairs. We observed the highest viral load in throat swabs at the time of symptom onset, and inferred that infectiousness peaked on or before symptom onset. We estimated that 44% (95% confidence interval, 25–69%) of secondary cases were infected during the index cases’ presymptomatic stage, in settings with substantial household clustering, active case finding and quarantine outside the home. Disease control measures should be adjusted to account for probable substantial presymptomatic transmission.
150
 Preprint DOI:	10.1101/2020.03.15.992818
1

The SARS-CoV-2 epidemic has rapidly spread outside China with major outbreaks occurring in Italy, South Korea, and Iran. Phylogenetic analyses of whole-genome sequencing data identified a distinct SARS-CoV-2 clade linked to travellers returning from Iran to Australia and New Zealand. This study highlights potential viral diversity driving the epidemic in Iran, and underscores the power of rapid genome sequencing and public data sharing to improve the detection and management of emerging infectious diseases.
151
 Preprint DOI:	10.1101/2020.03.15.992925
1

The recent epidemic outbreak of a novel human coronavirus called SARS-CoV-2 causing the respiratory tract disease COVID-19 has reached worldwide resonance and a global effort is being undertaken to characterize the molecular features and evolutionary origins of this virus. In this paper, we set out to shed light on the SARS-CoV-2/host receptor recognition, a crucial factor for successful virus infection. Based on the current knowledge of the interactome between SARS-CoV-2 and host cell proteins, we performed Master Regulator Analysis to detect which parts of the human interactome are most affected by the infection. We detected, amongst others, affected apoptotic and mitochondrial mechanisms, and a downregulation of the ACE2 protein receptor, notions that can be used to develop specific therapies against this new virus.
152
 Preprint DOI:	10.1101/2020.03.16.20035014
1

At present, PCR-based nucleic acid detection cannot meet the demands for coronavirus infectious disease (COVID-19) diagnosis. Two hundred fourteen confirmed COVID-19 patients who were hospitalized in the General Hospital of Central Theater Command of the People’s Liberation Army between 18 January and 26 February 2020 were recruited. Two enzyme-linked immunosorbent assay (ELISA) kits based on recombinant severe acute respiratory syndrome coronavirus 2 (SARS-CoV-2) nucleocapsid protein (rN) and spike protein (rS) were used for detecting IgM and IgG antibodies, and their diagnostic feasibility was evaluated. Among the 214 patients, 146 (68.2%) and 150 (70.1%) were successfully diagnosed with the rN-based IgM and IgG ELISAs, respectively; 165 (77.1%) and 159 (74.3%) were successfully diagnosed with the rS-based IgM and IgG ELISAs, respectively. The positive rates of the rN-based and rS-based ELISAs for antibody (IgM and/or IgG) detection were 80.4% and 82.2%, respectively. The sensitivity of the rS-based ELISA for IgM detection was significantly higher than that of the rN-based ELISA. We observed an increase in the positive rate for IgM and IgG with an increasing number of days post-disease onset (d.p.o.), but the positive rate of IgM dropped after 35 d.p.o. The positive rate of rN-based and rS-based IgM and IgG ELISAs was less than 60% during the early stage of the illness, 0 to 10 d.p.o., and that of IgM and IgG was obviously increased after 10 d.p.o. ELISA has a high sensitivity, especially for the detection of serum samples from patients after 10 d.p.o., so it could be an important supplementary method for COVID-19 diagnosis.
153
 Preprint DOI:	10.1101/2020.03.16.20037135
1

"BackgroundChloroquine and hydroxychloroquine have been found to be efficient on SARS-CoV-2, and reported to be efficient in Chinese COV-19 patients. We evaluate the role of hydroxychloroquine on respiratory viral loads.Patients and methodsFrench Confirmed COVID-19 patients were included in a single arm protocol from early March to March 16th, to receive 600mg of hydroxychloroquine daily and their viral load in nasopharyngeal swabs was tested daily in a hospital setting. Depending on their clinical presentation, azithromycin was added to the treatment. Untreated patients from another center and cases refusing the protocol were included as negative controls. Presence and absence of virus at Day6-post inclusion was considered the end point.ResultsSix patients were asymptomatic, 22 had upper respiratory tract infection symptoms and eight had lower respiratory tract infection symptoms....Twenty cases were treated in this study and showed a significant reduction of the viral carriage at D6-post inclusion compared to controls, and much lower average carrying duration than reported of untreated patients in the literature. Azithromycin added to hydroxychloroquine was significantly more efficient for virus elimination.ConclusionDespite its small sample size our survey shows that hydroxychloroquine treatment is significantly associated with viral load reduction/disappearance in COVID-19 patients and its effect is reinforced by azithromycin."
154
 Preprint DOI:	10.1101/2020.03.17.20037408
1

"ObjectivesThe December 2019 outbreak of coronavirus has once again thrown the vexed issue of quarantine into the spotlight, with many countries asking their citizens to ‘self-isolate’ if they have potentially come into contact with the infection. However, adhering to quarantine is difficult. Decisions on how to apply quarantine should be based on the best available evidence to increase the likelihood of people adhering to protocols. We conducted a rapid review to identify factors associated with adherence to quarantine during infectious disease outbreaks.Study designThe study design is a rapid evidence review.MethodsWe searched Medline, PsycINFO and Web of Science for published literature on the reasons for and factors associated with adherence to quarantine during an infectious disease outbreak.ResultsWe found 3163 articles and included 14 in the review. Adherence to quarantine ranged from as little as 0 up to 92.8%. The main factors which influenced or were associated with adherence decisions were the knowledge people had about the disease and quarantine procedure, social norms, perceived benefits of quarantine and perceived risk of the disease, as well as practical issues such as running out of supplies or the financial consequences of being out of work.ConclusionsPeople vary in their adherence to quarantine during infectious disease outbreaks. To improve this, public health officials should provide a timely, clear rationale for quarantine and information about protocols; emphasise social norms to encourage this altruistic behaviour; increase the perceived benefit that engaging in quarantine will have on public health; and ensure that sufficient supplies of food, medication and other essentials are provided."
155
 Preprint DOI:	10.1101/2020.03.18.20037101
1

Public health preparedness for coronavirus (CoV) disease 2019 (COVID-19) is challenging in the absence of setting-specific epidemiological data. Here we describe the epidemiology of seasonal CoVs (sCoVs) and other cocirculating viruses in the West of Scotland, United Kingdom. We analyzed routine diagnostic data for >70 000 episodes of respiratory illness tested molecularly for multiple respiratory viruses between 2005 and 2017. Statistical associations with patient age and sex differed between CoV-229E, CoV-OC43, and CoV-NL63. Furthermore, the timing and magnitude of sCoV outbreaks did not occur concurrently, and coinfections were not reported. With respect to other cocirculating respiratory viruses, we found evidence of positive, rather than negative, interactions with sCoVs. These findings highlight the importance of considering cocirculating viruses in the differential diagnosis of COVID-19. Further work is needed to establish the occurrence/degree of cross-protective immunity conferred across sCoVs and with COVID-19, as well as the role of viral coinfection in COVID-19 disease severity.
156
 Preprint DOI:	10.1101/2020.03.18.20038059
1

"A new coronavirus, severe acute respiratory syndrome coronavirus 2 (SARS-CoV-2), has recently emerged to cause a human pandemic. Although molecular diagnostic tests were rapidly developed, serologic assays are still lacking, yet urgently needed. Validated serologic assays are needed for contact tracing, identifying the viral reservoir, and epidemiologic studies. We developed serologic assays for detection of SARS-CoV-2 neutralizing, spike protein-specific, and nucleocapsid-specific antibodies. Using serum samples from patients with PCR-confirmed SARS-CoV-2 infections, other coronaviruses, or other respiratory pathogenic infections, we validated and tested various antigens in different in-house and commercial ELISAs. We demonstrated that most PCR-confirmed SARS-CoV-2-infected persons seroconverted by 2 weeks after disease onset.. We found that commercial S1 IgG or IgA ELISAs were of lower specificity, and sensitivity varied between the 2 assays; the IgA ELISA showed higher sensitivity. Overall, the validated assays	described can be instrumental for detection of SARS-CoV-2-specific	antibodies for diagnostic, seroepidemiologic, and vaccine evaluation	studies."
159
 Preprint DOI:	10.1101/2020.03.19.20038950
1

"Background: The rapid spread of COVID-19 virus from China to other countries and outbreaks of disease require an epidemiological analysis of the disease in the shortest time and an increased awareness of effective interventions. The purpose of this study was to estimate the COVID-19 epidemic in Iran based on the SIR model. The results of the analysis of the epidemiological data of Iran from January 22 to March 24, 2020 were investigated and prediction was made until April 15, 2020. Methods: By estimating the three parameters of time-dependent transmission rate, time-dependent recovery rate, and timedependent death rate from Covid-19 outbreak in China, and using the number of Covid-19 infections in Iran, we predicted the number of patients for the next month in Iran. Each of these parameters was estimated using GAM models. All analyses were conducted in R software using the mgcv package. Results: Based on our predictions of Iran about 29000 people will be infected from March 25 to April 15, 2020.. On average, 1292 people with COVID-19 are expected to be infected daily in Iran. The epidemic peaks within 3 days (March 25 to March 27, 2020) and reaches its highest point on March 25, 2020 with 1715 infected cases. Conclusion: The most important point is to emphasize the timing of the epidemic peak, hospital readiness, government measures and public readiness to reduce social contact."
161
 Preprint DOI:	10.1101/2020.03.21.001933
1

The emergence of SARS-CoV-2 has resulted in nearly 1,280,000 infections and 73,000 deaths globally so far. This novel virus acquired the ability to infect human cells using the SARS-CoV cell receptor hACE2. Because of this, it is essential to improve our understanding of the evolutionary dynamics surrounding the SARS-CoV-2 hACE2 interaction. One way theory predicts selection pressures should shape viral evolution is to enhance binding with host cells. We first assessed evolutionary dynamics in select betacoronavirus spike protein genes to predict whether these genomic regions are under directional or purifying selection between divergent viral lineages, at various scales of relatedness. With this analysis, we determine a region inside the receptor-binding domain with putative sites under positive selection interspersed among highly conserved sites, which are implicated in structural stability of the viral spike protein and its union with human receptor ACE2. Next, to gain further insights into factors associated with recognition of the human host receptor, we performed modeling studies of five different betacoronaviruses and their potential binding to hACE2. Modeling results indicate that interfering with the salt bridges at hot spot 353 could be an effective strategy for inhibiting binding, and hence for the prevention of SARS-CoV-2 infections. We also propose that a glycine residue at the receptor-binding domain of the spike glycoprotein can have a critical role in permitting bat SARS-related coronaviruses to infect human cells.
162
 Preprint DOI:	10.1101/2020.03.22.20040600
2

"Genetic variability across the three major histocompatibility complex (MHC) class I genes (human leukocyte antigen A [HLA-A], -B, and -C genes) may affect susceptibility to and severity of the disease caused by severe acute respiratory syndrome coronavirus 2 (SARS-CoV-2), the virus responsible for coronavirus disease 2019 (COVID-19). We performed a comprehensive in silico analysis of viral peptide-MHC class I binding affinity across 145 HLA-A, -B, and -C genotypes for all SARS-CoV-2 peptides. We further explored the potential for cross-protective immunity conferred by prior exposure to four common human coronaviruses. The SARS-CoV-2 proteome was successfully sampled and was represented by a diversity of HLA alleles. However, we found that HLA-B*46:01 had the fewest predicted binding peptides for SARS-CoV-2, suggesting that individuals with this allele may be particularly vulnerable to COVID-19, as they were previously shown to be for SARS (M. Lin, H.-T. Tseng, J. A. Trejaut, H.-L. Lee, et al., BMC Med Genet 4:9, 2003, https://bmcmedgenet.biomedcentral.com/articles/10.1186/1471-2350-4-9). Conversely, we found that HLA-B*15:03 showed the greatest capacity to present highly conserved SARS-CoV-2 peptides that are shared among common human coronaviruses, suggesting that it could enable cross-protective T-cell-based immunity. Finally, we reported global distributions of HLA types with potential epidemiological ramifications in the setting of the current pandemic."
163
 Preprint DOI:	10.1101/2020.03.24.20041020
2

"Objective To review and critically appraise published and preprint reports of prediction models for diagnosing coronavirus disease 2019 (covid-19) in patients with suspected infection, for prognosis of patients with covid-19, and for detecting people in the general population at increased risk of becoming infected with covid-19 or being admitted to hospital with the disease.Design Living systematic review and critical appraisal.Data sources PubMed and Embase through Ovid, Arxiv, medRxiv, and bioRxiv up to 7 April 2020.Study selection Studies that developed or validated a multivariable covid-19 related prediction model.Data extraction At least two authors independently extracted data using the CHARMS (critical appraisal and data extraction for systematic reviews of prediction modelling studies) checklist; risk of bias was assessed using PROBAST (prediction model risk of bias assessment tool).Results 4909 titles were screened, and 51 studies describing 66 prediction models were included. The review identified three models for predicting hospital admission from pneumonia and other events (as proxy outcomes for covid-19 pneumonia) in the general population; 47 diagnostic models for detecting covid-19 (34 were based on medical imaging); and 16 prognostic models for predicting mortality risk, progression to severe disease,,,, or length of hospital stay. The most frequently reported predictors of presence of covid-19 included age, body temperature, , signs and symptoms, sex, blood pressure, and creatinine. . The most frequently reported predictors of severe prognosis in patients with covid-19 included age and features derived from computed tomography scans. C index estimates ranged from 0.73 to 0.81 in prediction models for the general population, from 0.65 to more than 0.99 in diagnostic models, and from 0.85 to 0.99 in prognostic models.... All models were rated at high or unclear risk of bias, mostly because of non-representative selection of control patients, exclusion of patients who had not experienced the event of interest by the end of the study, high risk of model overfitting, and vague reporting. Most reports did not include any description of the study population or intended use of the models, and calibration of the model predictions was rarely assessed.Conclusion Prediction models for covid-19 are quickly entering the academic literature to support medical decision making at a time when they are urgently needed. This review indicates that proposed models are poorly reported, at high risk of bias, and their reported performance is probably optimistic.... Hence, we do not recommend any of these reported prediction models to be used in current practice. . Immediate sharing of well documented individual participant data from covid-19 studies and collaboration are urgently needed to develop more rigorous prediction models, and validate promising ones. The predictors identified in included models should be considered as candidate predictors for new models. Methodological guidance should be followed because unreliable predictions could cause more harm than benefit in guiding clinical decisions. Finally, studies should adhere to the TRIPOD (transparent reporting of a multivariable prediction model for individual prognosis or diagnosis) reporting guideline."
164
 Preprint DOI:	10.1101/2020.03.24.20042432
1

"ObjectiveThe purpose of this study was to distinguish the imaging features of COVID-19 from those of other infectious pulmonary diseases and evaluate the diagnostic value of chest CT for suspected COVID-19 patients.MethodsAdult patients suspected of COVID-19 aged >18 years who underwent chest CT scans and reverse-transcription polymerase chain reaction (RT-PCR) tests within 14 days of symptom onset were enrolled. The enrolled patients were confirmed and grouped according to the results of the RT-PCR tests. The basic demographics, single chest CT features, and combined chest CT features were analyzed for the confirmed and nonconfirmed groups.ResultsA total of 130 patients were enrolled, with 54 testing positive and 76 testing negative. The typical CT imaging features of the positive group were ground glass opacities (GGOs), the crazy-paving pattern and air bronchogram. The lesions were mostly distributed bilaterally and close to the lower lungs or the pleura. When features were combined, GGOs with bilateral pulmonary distribution and GGOs with pleural distribution were more common among the positive patients, found in 31 (57.4%) and 30 patients (55.6%), respectively. The combinations were almost all statistically significant (P < .05), except for the combination of GGOs with consolidation. Most combinations presented relatively low sensitivity but extremely high specificity. The average specificity of these combinations was approximately 90%.ConclusionsThe combinations with GGOs could be useful in the identification and differential diagnosis of COVID-19, alerting clinicians to isolate patients for prompt treatment and repeat RT-PCR tests until the end of incubation."
165
 Preprint DOI:	10.1101/2020.03.24.20042598
1

We analyzed age-/sex-specific morbidity and mortality data from the SARS-CoV-2 pandemic in China and Republic of Korea (ROK). Data from China exhibit a Gaussian distribution with peak morbidity in the 50–59-year cohort, while the ROK data have a bimodal distribution with the highest morbidity in the 20–29-year cohort.
167
 Preprint DOI:	10.1101/2020.03.27.20043661
1

"Given the rapidly progressing coronavirus disease 2019 (COVID-19) pandemic, this report on a US cohort of 54 COVID-19 patients from Stanford Hospital and data regarding risk factors for severe disease obtained at initial clinical presentation is highly important and immediately clinically relevant. We identified low presenting oxygen saturation as predictive of severe disease outcomes, such as diagnosis of pneumonia, acute respiratory distress syndrome, and admission to the intensive care unit, and also replicated data from China suggesting an association between hypertension and disease severity. Clinicians will benefit by tools to rapidly risk stratify patients at presentation by likelihood of progression to severe disease."
168
 Preprint DOI:	10.1101/2020.03.29.20043547
1

"Background: In December 2019, a few coronavirus disease (COVID-19) cases were first reported in Wuhan, Hubei, China. Soon after, increasing numbers of cases were detected in other parts of China, eventually leading to a disease outbreak in China. As this dreadful disease spreads rapidly, the mass media has been active in community education on COVID-19 by delivering health information about this novel coronavirus, such as its pathogenesis, spread, prevention, and containment.Objective: The aim of this study was to collect media reports on COVID-19 and investigate the patterns of media-directed health communications as well as the role of the media in this ongoing COVID-19 crisis in China.Methods: We adopted the WiseSearch database to extract related news articles about the coronavirus from major press media between January 1, 2020, and February 20, 2020. We then sorted and analyzed the data using Python software and Python package Jieba. We sought a suitable topic number with evidence of the coherence number. We operated latent Dirichlet allocation topic modeling with a suitable topic number and generated corresponding keywords and topic names. We then divided these topics into different themes by plotting them into a 2D plane via multidimensional scaling.Results: After removing duplications and irrelevant reports, our search identified 7791 relevant news reports. We listed the number of articles published per day. According to the coherence value, we chose 20 as the number of topics and generated the topics’ themes and keywords. These topics were categorized into nine main primary themes based on the topic visualization figure. The top three most popular themes were prevention and control procedures, medical treatment and research, and global or local social and economic influences, accounting for 32.57% (n=2538), 16.08% (n=1258), and 11.79% (n=919) 9) of the collected reports, respectively.Conclusions: Topic modeling of news articles can produce useful information about the significance of mass media for early health communication. Comparing the number of articles for each day and the outbreak development, we noted that mass media news reports in China lagged behind the development of COVID-19. The major themes accounted for around half the content and tended to focus on the larger society rather than on individuals. The COVID-19 crisis has become a worldwide issue, and society has become concerned about donations and support as well as mental health among others. We recommend that future work addresses the mass media’s actual impact on readers during the COVID-19 crisis through sentiment analysis of news data."
169
 Preprint DOI:	10.1101/2020.03.30.20044099
1

COVID-19 has become a pandemic. The influence of meteorological factors on the transmission and spread of COVID-19 is of interest. This study sought to examine the associations of daily average temperature (AT) and relative humidity (ARH) with the daily counts of COVID-19 cases in 30 Chinese provinces (in Hubei from December 1, 2019 to February 11, 2020 and in other provinces from January 20, 2020 to Februarys 11, 2020). A Generalized Additive Model (GAM) was fitted to quantify the province-specific associations between meteorological variables and the daily cases of COVID-19 during the study periods. In the model, the 14-day exponential moving averages (EMAs) of AT and ARH, and their interaction were included with time trend and health-seeking behavior adjusted. Their spatial distributions were visualized. AT and ARH showed significantly negative associations with COVID-19 with a significant interaction between them (0.04, 95% confidence interval: 0.004–0.07) in Hubei. Every 1 °C increase in the AT led to a decrease in the daily confirmed cases by 36% to 57% when ARH was in the range from 67% to 85.5%. Every 1% increase in ARH led to a decrease in the daily confirmed cases by 11% to 22% when AT was in the range from 5.04 °C to 8.2 °C. However, these associations were not consistent throughout Mainland China.
170
 Preprint DOI:	10.1101/2020.03.30.20047852
1

Infection caused by SARS-CoV-2 can result in severe respiratory complications and death. Patients with a compromised immune system are expected to be more susceptible to a severe disease course. In this report we suggest that patients with systemic lupus erythematous might be especially prone to severe COVID-19 independent of their immunosuppressed state from lupus treatment. Specifically, we provide evidence in lupus to suggest hypomethylation and overexpression of ACE2, which is located on the X chromosome and encodes a functional receptor for the SARS-CoV-2 spike glycoprotein. Oxidative stress induced by viral infections exacerbates the DNA methylation defect in lupus, possibly resulting in further ACE2 hypomethylation and enhanced viremia. In addition, demethylation of interferon-regulated genes, NFκB, and key cytokine genes in lupus patients might exacerbate the immune response to SARS-CoV-2 and increase the likelihood of cytokine storm. These arguments suggest that inherent epigenetic dysregulation in lupus might facilitate viral entry, viremia, and an excessive immune response to SARS-CoV-2. Further, maintaining disease remission in lupus patients is critical to prevent a vicious cycle of demethylation and increased oxidative stress, which will exacerbate susceptibility to SARS-CoV-2 infection during the current pandemic. Epigenetic control of the ACE2 gene might be a target for prevention and therapy in COVID-19.
171
 Preprint DOI:	10.1101/2020.03.31.016170
1

Yeast tolerates a low pH and high solvent concentrations. The permeability of the plasma membrane (PM) for small molecules is low and lateral diffusion of proteins is slow. These findings suggest a high degree of lipid order, which raises the question of how membrane proteins function in such an environment. The yeast PM is segregated into the Micro-Compartment-of-Can1 (MCC) and Pma1 (MCP), which have different lipid compositions. We extracted proteins from these microdomains via stoichiometric capture of lipids and proteins in styrene-maleic-acid-lipid-particles (SMALPs). We purified SMALP-lipid-protein complexes by chromatography and quantitatively analyzed periprotein lipids located within the diameter defined by one SMALP. Phospholipid and sterol concentrations are similar for MCC and MCP, but sphingolipids are enriched in MCP. Ergosterol is depleted from this periprotein lipidome, whereas phosphatidylserine is enriched relative to the bulk of the plasma membrane. Direct detection of PM lipids in the 'periprotein space' supports the conclusion that proteins function in the presence of a locally disordered lipid state.
172
 Preprint DOI:	10.1101/2020.04.02.20051029
1

"Background Severe acute respiratory syndrome coronavirus-2 (SARS-CoV-2) is the causative	agent of coronavirus disease 2019 (COVID-19), which was declared a	global pandemic by the World Health Organization on 11th March 2020. The treatment guidelines for COVID-19 vary between countries, yet there is no approved treatment to date.AimTo report any evidence of therapeutics used for the management of patients with COVID-19 in clinical practice since emergence of the virus. Methods A systematic review protocol was developed based on the PRISMA statement. Articles for review were selected from Embase, Medline and Google	Scholar. Readily accessible peer-reviewed, full articles in English	published from 1st December 2019 to 26th March	2020 were included. The search terms included combinations of: COVID, SARS-COV-2, glucocorticoids, convalescent plasma, antiviral and	antibacterial. There were no restrictions on the types of study eligible for inclusion. Results Four hundred and forty-nine articles were identified in the literature	search; of these, 41 studies were included in this review. These were	clinical trials (N=3), case reports (N=7), case series (N=10), and retrospective (N=11) and prospective (N=10) observational studies. Thirty-six studies were conducted in China	(88%). Corticosteroid treatment was reported most frequently (N=25), followed by lopinavir (N=21) and oseltamivir (N=16). Conclusions This is the first systematic review to date related to medication used to	treat patients with COVID-19. Only 41 studies were eligible for	inclusion, most of which were conducted in China. Corticosteroid	treatment was reported most frequently in the literature."
173
 Preprint DOI:	10.1101/2020.04.02.20051128
1

Covid-19 originated in Wuhan and rippled across China. We investigate how the geographical distance of working adults to the epicenter of Wuhan predicts their burnout - emotional, physical and mental exhaustion due to excessive and prolonged stress. .Preliminary results of a survey of 308 working adults in 53 cities showed working adults’ distance to the epicenter of Wuhan had an inverted U-shaped relationship with their burnout. Such results help to identify regions where people may need more psychiatric assistance, with direct implications for healthcare practitioners and policymakers.

175
 Preprint DOI:	10.1101/2020.04.03.20050195
2

"ObjectivesTo establish the optimal parameters for group testing of pooled specimens for the detection of SARS-CoV-2.MethodsThe most efficient pool size was determined to be five specimens using a web-based application. From this analysis, 25 experimental pools were created using 50 µL from one SARS-CoV-2 positive nasopharyngeal specimen mixed with 4 negative patient specimens (50 µL each) for a total volume of 250 µL. Viral RNA was subsequently extracted from each pool and tested using the CDC SARS-CoV-2 RT-PCR assay. Positive pools were consequently split into individual specimens and tested by extraction and PCR. This method was also tested on an unselected group of 60 nasopharyngeal specimens grouped into 12 pools.ResultsAll 25 pools were positive with cycle threshold (Ct) values within 0 and 5.03 Ct of the original individual specimens. The analysis of 60 specimens determined that 2 pools were positive followed by identification of 2 individual specimens among the 60 tested. This testing was accomplished while using 22 extractions/PCR tests, a savings of 38 reactions.ConclusionsWhen the incidence rate of SARS-CoV-2 infection is 10% or less, group testing willwillwill result in the saving of reagents and personnel time with an overall increase in testing capability of at least 69%."
176
 Preprint DOI:	10.1101/755652
1

Inhibitory codon pairs and poly(A) tracts within the translated mRNA cause ribosome stalling and reduce protein output. The molecular mechanisms that drive these stalling events, however, are still unknown. Here, we use a combination of in vitro biochemistry, ribosome profiling, and cryo‐EM to define molecular mechanisms that lead to these ribosome stalls. First, we use an in vitro reconstituted yeast translation system to demonstrate that inhibitory codon pairs slow elongation rates which are partially rescued by increased tRNA concentration or by an artificial tRNA not dependent on wobble base‐pairing. Ribosome profiling data extend these observations by revealing that paused ribosomes with empty A sites are enriched on these sequences. Cryo‐EM structures of stalled ribosomes provide a structural explanation for the observed effects by showing decoding‐incompatible conformations of mRNA in the A sites of all studied stall‐ and collision‐inducing sequences. Interestingly, in the case of poly(A) tracts, the inhibitory conformation of the mRNA in the A site involves a nucleotide stacking array. Together, these data demonstrate a novel mRNA ‐induced mechanisms of translational stalling in eukaryotic ribosomes.
177
 Preprint DOI:	10.1101/756064
1

"Background: Module detection algorithms relying on modularity maximization suffer from an inherent resolution limit that hinders detection of small topological modules, especially in molecular networks where most biological processes are believed to form small and compact communities. We propose a novel modular refinement approach that helps finding functionally significant modules of molecular networks.Results: The module refinement algorithm improves the quality of topological modules in protein-protein interaction networks by finding biologically functionally significant modules. The algorithm is based on the fact that functional modules in biology do not necessarily represent those corresponding to maximum modularity. Larger modules corresponding to maximal modularity are incrementally re-modularized again under specific constraints so that smaller yet topologically and biologically valid modules are recovered. We show improvement in quality and functional coverage of modules using experiments on synthetic and real protein-protein interaction networks. We also compare our results with six existing methods available for clustering biological networks.Conclusion: The proposed algorithm finds smaller but functionally relevant modules that are undetected by classical quality maximization approaches for modular detection. The refinement procedure helps to detect more functionally enriched modules in protein-protein interaction networks, which are also more coherent with functionally characterised gene sets."
178
 Preprint DOI:	10.1101/768648
1

"With emerging resistance to frontline treatments, it is vital that new drugs are identified to target Plasmodium falciparum. One of the most critical processes during parasites asexual lifecycle	is the invasion and subsequent egress of red blood cells (RBCs). Many	unique parasite ligands, receptors and enzymes are employed during	egress and invasion that are essential for parasite proliferation and	survival, therefore making these processes druggable targets. To	identify potential inhibitors of egress and invasion, we screened the	Medicines for Malaria Venture Pathogen Box, a 400 compound library	against neglected tropical diseases, including 125 with antimalarial	activity. For this screen, we utilised transgenic parasites expressing a bioluminescent reporter, nanoluciferase (Nluc), to measure inhibition	of parasite egress and invasion in the presence of the Pathogen Box	compounds. At a concentration of 2 µM, we found 15 compounds that	inhibited parasite egress by >40% and 24 invasion-specific compounds	that inhibited invasion by >90%. We further characterised 11 of these inhibitors through cell-based assays and live cell microscopy, and	found two compounds that inhibited merozoite maturation in schizonts, one compound that inhibited merozoite egress, one compound that directly inhibited parasite invasion and one compound that slowed down invasion	and arrested ring formation. The remaining compounds were general growth inhibitors that acted during the egress and invasion phase of the cell	cycle. We found the sulfonylpiperazine, MMV020291, to be the most	invasion-specific inhibitor, blocking successful merozoite	internalisation within human RBCs and having no substantial effect on	other stages of the cell cycle. This has significant implications for	the possible development of an invasion-specific inhibitor as an	antimalarial in a combination based therapy, in addition to being a	useful tool for studying the biology of the invading parasite."
179
 Preprint DOI:	10.1101/773812
1

The actin nucleators Spire and Cappuccino synergize to promote actin assembly, but the mechanism of their synergy is controversial. Together these proteins promote the formation of actin meshes, which are conserved structures that regulate the establishment of oocyte polarity. Direct interaction between Spire and Cappuccino is required for oogenesis and for in vitro synergistic actin assembly. This synergy is proposed to be driven by elongation and the formation of a ternary complex at filament barbed ends, or by nucleation and interaction at filament pointed ends. To mimic the geometry of Spire and Cappuccino in vivo, we immobilized Spire on beads and added Cappuccino and actin. Barbed ends, protected by Cappuccino, grow away from the beads while pointed ends are retained, as expected for nucleation-driven synergy. We found that Spire is sufficient to bind barbed ends and retain pointed ends of actin filaments near beads and we identified Spire’s barbed-end binding domain. Loss of barbed-end binding increases nucleation by Spire and synergy with Cappuccino in bulk pyrene assays and on beads. Importantly, genetic rescue by the loss-of-function mutant indicates that barbed-end binding is not necessary for oogenesiss.oogenesiss. Thus, increased nucleation is a critical element of synergy both in vitro and in vivo.	.
180
 Preprint DOI:	10.1101/774372
1

"Taxonomic and functional information from microbial communities can be	efficiently obtained by metagenome profiling, which requires databases	of genes and genomes to which sequence reads are mapped. However, the	databases that accompany metagenome profilers are not updated at a pace	that matches the increase in available microbial genomes, and unifying	database content across metagenome profiling tools can be cumbersome. To address this, we developed Struo, a modular pipeline that automatizes	the acquisition of genomes from public repositories and the construction of custom databases for multiple metagenome profilers. The use of	custom databases that broadly represent the known microbial diversity by incorporating novel genomes results in a substantial increase in	mappability of reads in synthetic and real metagenome datasets."
181
 Preprint DOI:	10.1101/782359
1

This investigation examined anthropometric, hormonal, and physiological differences between advanced (ADV; n = 8, 27.8 ± 4.2 years, 170 ± 11 cm, 79.8 ± 13.3 kg) and recreational (REC; n = 8, 33.5 ± 8.1 years, 172 ± 14 cm, 76.3 ± 19.5 kg) CrossFit (CF) trained participants in comparison to physically-active controls (CON; n = 7, 27.5 ± 6.7 years, 171 ± 14 cm, 74.5 ± 14.3 kg). ADV and REC were distinguished by their past competitive success. REC and CON were resistance-trained (>2 years) and exercised on 3–5 days·wk-1 for the past year, but CON utilized traditional resistance and cardiovascular exercise. All participants provided a fasted, resting blood sample and completed assessments of resting metabolic rate, body composition, muscle morphology, isometric mid-thigh pull strength, peak aerobic capacity, and a 3-minute maximal cycle ergometer sprint across two separate occasions (separated by 3–7 days). Blood samples were analyzed for testosterone, cortisol, and insulin-like growth factor-1. Compared to both REC and CON, one-way analysis of variance revealed ADV to possess lower body fat percentage (6.7–8.3%, p = 0.007), greater bone and non-bone lean mass (12.5–26.8%, p ≤ 0.028), muscle morphology characteristics (14.2–59.9%, p < 0.05), isometric strength characteristics (15.4–41.8%, p < 0.05), peak aerobic capacity (18.8–19.1%, p = 0.002), and 3-minute cycling performance (15.4–51.1%, p ≤ 0.023). No differences were seen between REC and CON, or between all groups for resting metabolic rate or hormone concentrations. These data suggest ADV possess several physiological advantages over REC and CON, whereas similar physiological characteristics were present in individuals who have been regularly participating in either CF or resistance and cardiovascular training for the past year.
182
 Preprint DOI:	10.1101/782839
1

"BackgroundPhoronids, rhynchonelliform and linguliform brachiopods show striking similarities in their embryonic fate maps, in particular in their axis specification and regionalization. However, although brachiopod development has been studied in detail and demonstrated embryonic patterning as a causal factor of the gastrulation mode (protostomy vs deuterostomy), molecular descriptions are still missing in phoronids. To understand whether phoronids display underlying embryonic molecular mechanisms similar to those of brachiopodsbrachiopodsbrachiopodsbrachiopods, here we report the expression patterns of anterior (otx, gsc, six3/6, nk2.1), posterior (cdx, bra) and endomesodermal (foxA, gata4/5/6, twist) markers during the development of the protostomic phoronid Phoronopsis harmeri.ResultsThe transcription factors foxA, gata4/5/6 and cdx show conserved expression in patterning the development and regionalization of the phoronid embryonic gut, with foxA expressed in the presumptive foregut, gata4/5/6 demarcating the midgut and cdx confined to the hindgut. Furthermore, six3/6, usually a well-conserved anterior marker, shows a remarkably dynamic expression, demarcating not only the apical organ and the oral ectoderm, but also clusters of cells of the developing midgut and the anterior mesoderm, similar to what has been reported for brachiopods, bryozoans and some deuterostome Bilateria. Surprisingly, brachyury, a transcription factor often associated with gastrulation movements and mouth and hindgut development, seems not to be involved with these patterning events in phoronids.ConclusionsOur description and comparison of gene expression patterns with other studied Bilateria reveals that the timing of axis determination and cell fate distribution of the phoronid shows highest similarity to that of rhynchonelliform brachiopods, which is likely related to their shared protostomic mode of development. . Despite these similarities, the phoronid Ph. harmeri also shows particularities in its development, which hint to divergences in the arrangement of gene regulatory networks responsible for germ layer formation and axis specification."
183
 Preprint DOI:	10.1101/785691
2

The evolution of regulatory networks in Bacteria has largely been explained at macroevolutionary scales through lateral gene transfer and gene duplication. Transcription factors (TF) have been found to be less conserved across species than their target genes (TG). This would be expected if TFs accumulate mutations faster than TGs. This hypothesis is supported by several lab evolution studies which found TFs, especially global regulators, to be frequently mutated. Despite these studies, the contribution of point mutations in TFs to the evolution of regulatory network is poorly understood. We tested if TFs show greater genetic variation than their TGs using whole-genome sequencing data from a large collection of Escherichia coli isolates. TFs were less diverse than their TGs across natural isolates, with TFs of large regulons being more conserved. In contrast, TFs showed higher mutation frequency in adaptive laboratory evolution experiments. However, over long-term laboratory evolution spanning 60 000 generations, mutation frequency in TFs gradually declined after a rapid initial burst. Extrapolating the dynamics of genetic variation from long-term laboratory evolution to natural populations, we propose that point mutations, conferring large-scale gene expression changes, may drive the early stages of adaptation but gene regulation is subjected to stronger purifying selection post adaptation.
184
 Preprint DOI:	10.1101/791269
1

"Aim	Biotic interactions can determine rarity and	commonness of species, however, evidence that rare and common species	respond differently to biotic stress is scarce. This is because biotic	interactions are notoriously context dependent and traits leading to	success in one habitat might be costly or unimportant in another. We aim to identify plant characteristics that are related to biotic	interactions and may drive patterns of rarity and commonness, taking	environmental context into account. Location Switzerland. Methods In a multispecies experiment, we compared the response to biotic interactions of 19 rare and 21 widespread congeneric plant	species in Switzerland, while also accounting for variation in	environmental conditions of the species' origin. Results Our results restrict the long-standing hypothesis that widespread species are superior competitors to rare species to only	those species originating from resource-rich habitats, in which	competition is usually strong. Tolerance to herbivory and ambient	herbivore damage, on the other hand, did not differ between widespread	and rare species. In accordance with the resource-availability	hypothesis, widespread species from resource-rich habitats where more	damaged by herbivores (less defended) than widespread species from	resource-poor habitats—such a growth-defence trade-off was lacking in	rare species. This indicates that the evolutionary important trade-off	between traits increasing competitive ability and defence is present in	widespread species but may have been lost or never evolved in rare	species. Main conclusions Our results indicate that biotic interactions, above	all competition, might indeed set range limits and underlines the	importance of including context dependency in studies comparing traits	of common and rare or invasive and non-invasive species."
185

Preprint DOI 10.1101/793000

Redone because PubMed had a truncated abstract

Objective Rates of overweight and obesity epidemic have risen significantly in the past few decades, and 34% of adults and 15-20% of children and adolescents in the United States are now obese. Melanocortin receptor 4 (MC4R), contributes to appetite control in hypothalamic neurons and is a target for future anti-obesity treatments (such as setmelanotide) or novel drug development effort. Proper MC4R trafficking regulation in hypothalamic neurons is crucial for normal neural control of homeostasis and is altered in obesity and in presence of lipids. The mechanisms underlying altered MC4R trafficking in the context of obesity is still unclear. Here, we discovered that C2CD5 expressed in the hypothalamus is involved in the regulation of MC4R endocytosis. This study unmasked a novel trafficking protein nutritionally regulated in the hypothalamus providing a novel target for MC4R dependent pathways involved in bodyweight homeostasis and Obesity.

Methods To evaluate the expression of C2cd5, we first used in situ hybridization and RNAscope technology in combination with electronic microscopy. For in vivo, we characterized the energy balance of wild type (WT) and C2CD5 whole-body knockout (C2CD5KO) mice fed normal chow (NC) and/or western-diet (high-fat/high-sucrose/cholesterol) (WD). To this end, we performed comprehensive longitudinal assessment of bodyweight, energy balance (food intake, energy expenditure, locomotor activity using TSE metabolic cages), and glucose homeostasis. In addition, we evaluated the consequence of loss of C2CD5 on feeding behavior changes normally induced by MC4R agonist (Melanotan, MTII) injection in the paraventricular hypothalamus (PVH). For in vitro approach, we tease out the role of C2CD5 and its calcium sensing domain C2 in MC4R trafficking. We focused on endocytosis of MC4R using an antibody feeding experiment (in a neuronal cell line - Neuro2A (N2A) stably expressing HA-MC4R-GFP; against HA-tag and analyzed by flux cytometry).

Results We found that 1) the expression of hypothalamic C2CD5 is decreased in diet-induced obesity models compared to controls, 2) mice lacking C2CD5 exhibit an increase in food intake compared to WT mice, 3) C2CD5 interacts with endocytosis machinery in hypothalamus, 4) loss of functional C2CD5 (lacking C2 domain) blunts MC4R endocytosis in vitro and increases MC4R at the surface that fails to respond to MC4R ligand, and, 5) C2CD5KO mice exhibit decreased acute responses to MTII injection into the PVH.

Conclusions Based on these, we conclude that hypothalamic C2CD5 is involved in MC4R endocytosis and regulate bodyweight homeostasis. These studies suggest that C2CD5 represents a new protein regulated by metabolic cues and involved in metabolic receptor endocytosis. C2CD5 represent a new target and pathway that could be targeted in Obesity.

186
 Preprint DOI:	10.1101/796318
2

Cancer proteogenomics promises new insights into cancer biology and treatment efficacy by integrating genomics, transcriptomics and protein profiling including modifications by mass spectrometry (MS). A critical limitation is sample input requirements that exceed many sources of clinically important material. Here we report a proteogenomics approach for core biopsies using tissue-sparing specimen processing and microscaled proteomics. As a demonstration, we analyze core needle biopsies from ERBB2 positive breast cancers before and 48–72 h after initiating neoadjuvant trastuzumab-based chemotherapy. We show greater suppression of ERBB2 protein and both ERBB2 and mTOR target phosphosite levels in cases associated with pathological complete response, and identify potential causes of treatment resistance including the absence of ERBB2 amplification, insufficient ERBB2 activity for therapeutic sensitivity despite ERBB2 amplification, and candidate resistance mechanisms including androgen receptor signaling, mucin overexpression and an inactive immune microenvironment. The clinical utility and discovery potential of proteogenomics at biopsy-scale warrants further investigation.	.	.
187
 Preprint DOI:	10.1101/797183
5

As development proceeds, inductive cues are interpreted by competent tissues in a spatially and temporally restricted manner. While key inductive signaling pathways within competent cells are well-described at a molecular level, the mechanisms by which tissues lose responsiveness to inductive signals are not well understood. Localized activation of Wnt signaling before zygotic gene activation in Xenopus laevis leads to dorsal development, but competence to induce dorsal genes in response to Wnts is lost by the late blastula stage. We hypothesize that loss of competence is mediated by changes in histone modifications leading to a loss of chromatin accessibility at the promoters of Wnt target genes. We use ATAC-seq to evaluate genome-wide changes in chromatin accessibility across several developmental stages. Based on overlap with p300 binding, we identify thousands of putative cis-regulatory elements at the gastrula stage, including sites that lose accessibility by the end of gastrulation and are enriched for pluripotency factor binding motifs. Dorsal Wnt target gene promoters are not accessible after the loss of competence in the early gastrula while genes involved in mesoderm and neural crest development maintain accessibility at their promoters. Inhibition of histone deacetylases increases acetylation at the promoters of dorsal Wnt target genes and extends competence for dorsal gene induction by Wnt signaling. Histone deacetylase inhibition, however, is not sufficient to extend competence for mesoderm or neural crest induction. These data suggest that chromatin state regulates the loss of competence to inductive signals in a context-dependent manner.
188
 Preprint DOI:	10.1101/813378
2

miR-375 is a highly abundant miRNA in Merkel cell carcinoma (MCC). In other cancers, it acts as either a tumor suppressor or oncogene. While free-circulating miR-375 serves as a surrogate marker for tumor burden in patients with advanced MCC, its function within MCC cells has not been established. Nearly complete miR-375 knockdown in MCC cell lines was achieved using antagomiRs via nucleofection. The cell viability, growth characteristics, and morphology were not altered by this knockdown. miR-375 target genes and related signaling pathways were determined using Encyclopedia of RNA Interactomes (ENCORI) revealing Hippo signaling and epithelial to mesenchymal transition (EMT)--)-)--)-related genes likely to be regulated. Therefore, their expression was analyzed by multiplexed qRT-PCR after miR-375 knockdown, demonstrating only a limited change in expression. In summary, highly effective miR-375 knockdown in classical MCC cell lines did not significantly change the cell viability, morphology, or oncogenic signaling pathways. These observations render miR-375 an unlikely intracellular oncogene in MCC cells, thus suggesting that likely functions of miR-375 for the intercellular communication of MCC should be addressed.
189
 Preprint DOI:	10.1101/815662
1

"Stromal collagen is upregulated surrounding a solid tumor and presents	as dense, thick, linearized, and aligned bundles. The collagen bundles	are continually remodeled during tumor progression, and their	orientation with respect to the tumor boundary has been correlated with	invasive state. Currently, reconstituted-collagen gels are the standard in vitro tumor cell-extracellular matrix interaction model. The reticular, dense, and isotropic nanofiber (~900 nm-diameter, on average) gels do	not, howeverhoweverhoweverhowever, recapitulate the in vivo structural features of	collagen bundling and alignment. Here, we present a rapid and simple	method to fabricate bundles of collagen type I, whose average thickness	may be varied between about 4 µm and 9 µm dependent upon diluent	temperature and ionic strength. The durability and versatility of the	collagen bundles was demonstrated with their incorporation into two in vitro models where the thickness and alignment of the collagen bundles resembled various in vivo arrangements. First, collagen bundles aligned by a microfluidic device	elicited cancer cell contact guidance and enhanced their directional	migration. Second, the presence of the collagen bundles in a bio-inert	agarose hydrogel was shown to provide a route for cancer cell outgrowth. The unique structural features of the collagen bundles advance the	physiological relevance of in vitro collagen-based tumor models for accurately capturing tumor cell-extracellular matrix interactions."
190
 Preprint DOI:	10.1101/819037
1

Molecular interactions at the cellular interface mediate organized assembly of single cells into tissues and, thus, govern the development and physiology of multicellular organisms. Here, we developed a cell-type-specific, spatiotemporally resolved approach to profile cell-surface proteomes in intact tissues. Quantitative profiling of cell-surface proteomes of Drosophila olfactory projection neurons (PNs) in pupae and adults revealed global downregulation of wiring molecules and upregulation of synaptic molecules in the transition from developing to mature PNs. A proteome-instructed in vivo screen identified 20 cell-surface molecules regulating neural circuit assembly, many of which belong to evolutionarily conserved protein families not previously linked to neural development. Genetic analysis further revealed that the lipoprotein receptor LRP1 cell-autonomously controls PN dendrite targeting, contributing to the formation of a precise olfactory map. These findings highlight the power of temporally resolved in situ cell-surface proteomic profiling in discovering regulators of brain wiring.
192
 Preprint DOI:	10.1101/832675
1

"The tricarboxylic acid (TCA) cycle is a central metabolic hub in most cells. Virulence functions of bacterial pathogens such as facultative intracellular Salmonella enterica serovar Typhimurium (S. Typhimurium) are closely connected to cellular metabolism. During systematic analyses of mutant strains with defects in the TCA cycle, a strain deficient in all fumarase isoforms (ΔfumABC) elicited a unique metabolic profile. Alongside fumarate, S. Typhimurium ΔfumABC accumulates intermediates of the glycolysis and pentose phosphate pathway. Analyses by metabolomics and proteomics revealed that fumarate accumulation redirects carbon fluxes toward glycogen synthesis due to high (p)ppGpp levels. In addition, we observed reduced abundance of CheY, leading to altered motility and increased phagocytosis of S. Typhimurium by macrophages. Deletion of glycogen synthase restored normal carbon fluxes and phagocytosis and partially restored levels of CheY. We propose that utilization of accumulated fumarate as carbon source induces a status similar to exponential- to stationary-growth-phase transition by switching from preferred carbon sources to fumarate, which increases (p)ppGpp levels and thereby glycogen synthesis. Thus, we observed a new form of interplay between metabolism of S. Typhimurium and cellular functions and virulence.IMPORTANCE We performed perturbation analyses of the tricarboxylic acid cycle of the gastrointestinal pathogen Salmonella enterica serovar Typhimurium. The defect of fumarase activity led to accumulation of fumarate but also resulted in a global alteration of carbon fluxes, leading to increased storage of glycogen. Gross alterations were observed in proteome and metabolome compositions of fumarase-deficient Salmonella. In turn, these changes were linked to aberrant motility patterns of the mutant strain and resulted in highly increased phagocytic uptake by macrophages. Our findings indicate that basic cellular functions and specific virulence functions in Salmonella critically depend on the proper function of the primary metabolism."
193
 Preprint DOI:	10.1101/838821
1

Ribosome biogenesis is tightly regulated through stress-sensing pathways that impact genome stability, aging and senescence. In Saccharomyces cerevisiae, ribosomal RNAs are transcribed from rDNA located on the right arm of chromosome XII. Numerous studies reveal that rDNA decondenses into a puff-like structure during interphase, and condenses into a tight loop-like structure during mitosis. Intriguingly, a novel and additional mechanism of increased mitotic rDNA compaction (termed hypercondensation) was recently discovered that occurs in response to temperature stress (hyperthermic-induced) and is rapidly reversible. Here, we report that neither changes in condensin binding or release of DNA during mitosis, nor mutation of factors that regulate cohesin binding and release, appear to play a critical role in hyperthermic-induced rDNA hypercondensation. A candidate genetic approach revealed that deletion of either HSP82 or HSC82 (Hsp90 encoding heat shock paralogs) result in significantly reduced hyperthermic-induced rDNA hypercondensation. Intriguingly, Hsp inhibitors do not impact rDNA hypercondensation. In combination, these findings suggest that Hsp90 either stabilizes client proteins, which are sensitive to very transient thermic challenges, or directly promotes rDNA hypercondensation during preanaphase. Our findings further reveal that the high mobility group protein Hmo1 is a negative regulator of mitotic rDNA condensation, distinct from its role in promoting premature condensation of rDNA during interphase upon nutrient starvation.
194
 Preprint DOI:	10.1101/838870
1

"Ten microsatellite loci were developed and validated for the endangered	cactus species Coleocephalocereus purpureus. The markers were obtained	from sequences generated by whole genome shotgun sequencing approaches. A testing group of 36 specimens of the main population were genotyped and all described markers presented suitable outcomes to population genetic studies, showing polymorphic status for C. purpureus testing group with clean and reproducible amplification. No evidence for scoring errors, null alleles or linkage disequilibrium was detected. Number of alleles	per locus ranged from 3 to 6 and expected heterozygosity ranged from	0.78 to 0.99. These new microsatellite loci are suitable to be used in	future diversity and structure population studies of C. purpureus."
195
 Preprint DOI:	10.1101/842203
1

Intra-tumoral heterogeneity (ITH) could represent clonal evolution where subclones with greater fitness confer more malignant phenotypes and invasion constitutes an evolutionary bottleneck. Alternatively, ITH could represent branching evolution with invasion of multiple subclones. The two models respectively predict a hierarchy of subclones arranged by phenotype, or multiple subclones with shared phenotypes. We delineate these modes of invasion by merging ancestral, topographic, and phenotypic information from 12 human colorectal tumors (11 carcinomas, 1 adenoma) obtained through saturation microdissection of 325 small tumor regions. The majority of subclones (29/46, 60%) share superficial and invasive phenotypes. Of 11 carcinomas, 9 show evidence of multiclonal invasion, and invasive and metastatic subclones arise early along the ancestral trees. Early multiclonal invasion in the majority of these tumors indicates the expansion of co-evolving subclones with similar malignant potential in absence of late bottlenecks and suggests that barriers to invasion are minimal during colorectal cancer growth.
196
 Preprint DOI:	10.1101/852541
1

Trophic interactions can result in changes to the abundance and distribution of habitat-forming species that dramatically reduce ecosystem functioning. In the coastal zone of the Aleutian Archipelago, overgrazing by herbivorous sea urchins that began in the 1990s resulted in widespread deforestation of the region’s kelp forests, which led to lower macroalgal abundances and higher benthic irradiances. We examined how this deforestation impacted ecosystem function by comparing patterns of net ecosystem production (NEP), gross primary production (GPP), ecosystem respiration (Re), and the range between GPP and Re in remnant kelp forests, urchin barrens, and habitats that were in transition between the two habitat types at nine islands that spanned more than 1000 kilometers of the archipelago. Our results show that deforestation, on average, resulted in a 24% reduction in GPP, a 26% reduction in Re, and a 24% reduction in the range between GPP and Re. . Further, the transition habitats were intermediate to the kelp forests and urchin barrens for these metrics.... These opposing metabolic processes remained in balance; however, which resulted in little-to-no changes to NEP. These effects of deforestation on ecosystem productivity, however, were highly variable between years and among the study islands. .In light of the worldwide declines in kelp forests observed in recent decades, our findings suggest that marine deforestation profoundly affects how coastal ecosystems function.
197
 Preprint DOI:	10.1101/859892
1

The telomerase reverse transcriptase (TERT) gene is responsible for telomere maintenance in germline and stem cells, and is re-expressed in 90% of human cancers. CpG methylation in the TERT promoter (TERTp) was correlated with TERT mRNA expression. Furthermore, two hotspot mutations in TERTp, dubbed C228T and C250T, have been revealed to facilitate binding of transcription factor ETS/TCF and subsequent TERT expression. This study aimed to elucidate the combined contribution of epigenetic (promoter methylation and chromatin accessibility) and genetic (promoter mutations) mechanisms in regulating TERT gene expression in healthy skin samples and in melanoma cell lines (n = 61). We unexpectedly observed that the methylation of TERTp was as high in a subset of healthy skin cells, mainly keratinocytes, as in cutaneous melanoma cell lines. In spite of the high promoter methylation fraction in wild-type (WT) samples, TERT mRNA was only expressed in the melanoma cell lines with either high methylation or intermediate methylation in combination with TERT mutations. TERTp methylation was positively correlated with chromatin accessibility and TERT mRNA expression in 8 melanoma cell lines. Cooperation between epigenetic and genetic mechanisms were best observed in heterozygous mutant cell lines as chromosome accessibility preferentially concerned the mutant allele. Combined, these results suggest a complex model in which TERT expression requires either a widely open chromatin state in TERTp-WT samples due to high methylation throughout the promoter or a combination of moderate methylation fraction/chromatin accessibility in the presence of the C228T or C250T mutations.
198
 Preprint DOI:	10.1101/863167
1

Isolation of high molecular weight DNA from gastropod molluscs and its subsequent PCR amplification is considered difficult due to excessive mucopolysaccharides secretion which co-precipitate with DNA and obstruct successful amplification. In an attempt to address this issue, we describe a modified CTAB DNA extraction method that proved to work significantly better with a number of freshwater and terrestrial gastropod taxa. We compared the performance of this method with Qiagen® DNeasy Blood and Tissue Kit. Reproducibility of amplification was verified using a set of taxon-specific primers, wherein modified CTAB extracted DNA could be replicated at least four out of five times but kit extracted DNA could not be replicated. In addition, sequence quality was significantly better with CTAB extracted DNA. This could be attributed to the removal of polyphenolic compounds by polyvinyl pyrrolidone which is the only difference between conventional and modified CTAB DNA extraction methods for animals. The genomic DNA isolated using modified CTAB protocol was of high quality (A260/280 ≥ 1.80) and could be used for downstream reactions even after long-term storage (more than 2 years).
199
 Preprint DOI:	10.1101/865089
1

The pathogenesis of spinal cord injury (SCI) remains poorly understood and treatment remains limited. Emerging evidence indicates that post-SCI inflammation is severe but the role of reactive astrogliosis not well understood given its implication in ongoing inflammation as damaging or neuroprotective. We have completed an extensive systematic study with MRI, histopathology, proteomics and ELISA analyses designed to further define the severe protracted and damaging inflammation after SCI in a rat model. We have identified 3 distinct phases of SCI: acute (first 2 days), inflammatory (starting day 3) and resolution (>3 months) in 16 weeks follow up. Actively phagocytizing, CD68+/CD163- macrophages infiltrate myelin-rich necrotic areas converting them into cavities of injury (COI) when deep in the spinal cord. Alternatively, superficial SCI areas are infiltrated by granulomatous tissue, or arachnoiditis where glial cells are obliterated. In the COI, CD68+/CD163- macrophage numbers reach a maximum in the first 4 weeks and then decline. Myelin phagocytosis is present at 16 weeks indicating ongoing inflammatory damage. The COI and arachnoiditis are defined by a wall of progressively hypertrophied astrocytes. MR imaging indicates persistent spinal cord edema that is linked to the severity of inflammation. Microhemorrhages in the spinal cord around the lesion are eliminated, presumably by reactive astrocytes within the first week post-injury. Acutely increased levels of TNF-alpha, IL-1beta, IFN-gamma and other pro-inflammatory cytokines, chemokines and proteases decrease and anti-inflammatory cytokines increase in later phases. In this study we elucidated a number of fundamental mechanisms in pathogenesis of SCI and have demonstrated a close association between progressive astrogliosis and reduction in the severity of inflammation.
200
 Preprint DOI:	10.1101/869313
1

"Colibactin is a genotoxic gut microbiome metabolite long suspected of playing an etiological role in colorectal cancer. Evidence suggests that colibactin forms DNA interstrand cross-links (ICLs) in eukaryotic cells and activates ICL repair pathways, leading to the production of ICL-dependent DNA double-strand breaks (DSBs). Here we show that colibactin ICLs can evolve directly to DNA DSBs. Using the topology of supercoiled plasmid DNA as a proxy for alkylation adduct stability, we find that colibactin-derived ICLs are unstable toward depurination and elimination of the 3' phosphate. This ICL degradation pathway leads progressively to single strand breaks (SSBs) and subsequently DSBs. The spontaneous conversion of ICLs to DSBs is consistent with the finding that nonhomologous end joining repair-deficient cells are sensitized to colibactin-producing bacteria. The results herein refine our understanding of colibactin-derived DNA damage and underscore the complexities underlying the DSB phenotype."
