## Supplemental Material 2. Annotations before reconciliation for "Preprints in motion: tracking changes between preprint posting and journal publication during a pandemic"

"With respect to observations by Monti et al,1 a survey featuring 31 questions related to rheumatic diseases (RDs) during the covid-19 pandemic was administered to members of the Indian Rheumatology Association.Of 861 invitees, 221 (25.7%; 92.7% adult rheumatologists, 52.2% academicians) ) responded. Most perceived the need for a change in the management of RDs (online supplementary files). Almost half (47.5%) reduced the usage of biological disease modifyinig anti rheumatic drugs (bDMARDs), whereas only 12.2% did so for csDMARDs (figure 1). Of the respondents, 66.5% were more inclined to initiate hydroxychloroquine (HCQ) in patients with borderline indications, whereas 14% disagreed with this approach. Nearly two-thirds (64.2%) were less inclined to change the major immunosuppressant (IS) for impending flare, with 58.3% deferring rituximab (RTX), followed closely by cyclophosphamide, antitumour necrosis factors (anti-TNFs), Janus kinase inhibitors (JAKinibs) and other bDMARDs. An earlier taper of glucocorticoids was preferred by 57.9% in inactive disease. There was lack of consensus on continuing IS infusions."
175
 Preprint DOI:	10.1101/2020.04.03.20050195
2
