## Supplementary material for "Preprints in motion: tracking changes between preprint posting and journal publication during a pandemic": Supp Methods 1

### Preprint-->paper evaluation form

* Required

1. Who are you? *

*Mark only one oval.*

- ( ) JC
- ( ) MP
- ( ) GD
- ( ) JP

2. Which dataset? *

*Mark only one oval.*

- ( ) COVID
- ( ) Control

3. Manuscript number (according to sheet) *

_______________________

4. Re-enter manuscript number (according to sheet) *

_______________________

5. Has the author list changed?

*Check all that apply.*

- [ ] No
- [ ] Yes, authors added
- [ ] Yes, authors removed
- [ ] Yes, change in corresponding/co-corresponding authors

6. Has the abstract changed?

*Mark only one oval.*

- No change or minor: change in wording, if any, does not change main conclusion(s)
- Yes, significant: altered wording or numbers leading to a softening/strengthening of main conclusions(s)
- Yes, major: a discrete change in the main conclusion(s)
- Yes, massive: main conclusion(s) of the paper reversed

7. How many figures/panels in the preprint (not supplement)?

*Check all that apply.*

| | 1 panel | 2 panels | 3 panels | 4 panels | 5 panels | 6 panels | 7 panels | 8 panels | 9 pane |
|---|---|---|---|---|---|---|---|---|---|
| Figure 1 | ☐ | ☐ | ☐ | ☐ | ☐ | ☐ | ☐ | ☐ | ☐ |
| Figure 2 | ☐ | ☐ | ☐ | ☐ | ☐ | ☐ | ☐ | ☐ | ☐ |
| Figure 3 | ☐ | ☐ | ☐ | ☐ | ☐ | ☐ | ☐ | ☐ | ☐ |
| Figure 4 | ☐ | ☐ | ☐ | ☐ | ☐ | ☐ | ☐ | ☐ | ☐ |
| Figure 5 | ☐ | ☐ | ☐ | ☐ | ☐ | ☐ | ☐ | ☐ | ☐ |
| Figure 6 | ☐ | ☐ | ☐ | ☐ | ☐ | ☐ | ☐ | ☐ | ☐ |
| Figure 7 | ☐ | ☐ | ☐ | ☐ | ☐ | ☐ | ☐ | ☐ | ☐ |
| Figure 8 | ☐ | ☐ | ☐ | ☐ | ☐ | ☐ | ☐ | ☐ | ☐ |
| Figure 9 | ☐ | ☐ | ☐ | ☐ | ☐ | ☐ | ☐ | ☐ | ☐ |
| Figure 10 or more (combine additional) | ☐ | ☐ | ☐ | ☐ | ☐ | ☐ | ☐ | ☐ | ☐ |

8. How many tables in the preprint (not supplement)?

*Mark only one oval.*

| | 0 | 1 | 2 | 3 | 4 | 5 | |
|---|---|---|---|---|---|---|---|
| | ◯ | ◯ | ◯ | ◯ | ◯ | ◯ | Choose 5 for >=5 |

9. How many figures/panels in the main paper (not supplement)?

*Check all that apply.*

| | 1 panel | 2 panels | 3 panels | 4 panels | 5 panels | 6 panels | 7 panels | 8 panels | 9 pane |
|---|---|---|---|---|---|---|---|---|---|
| Figure 1 | ☐ | ☐ | ☐ | ☐ | ☐ | ☐ | ☐ | ☐ | ☐ |
| Figure 2 | ☐ | ☐ | ☐ | ☐ | ☐ | ☐ | ☐ | ☐ | ☐ |
| Figure 3 | ☐ | ☐ | ☐ | ☐ | ☐ | ☐ | ☐ | ☐ | ☐ |
| Figure 4 | ☐ | ☐ | ☐ | ☐ | ☐ | ☐ | ☐ | ☐ | ☐ |
| Figure 5 | ☐ | ☐ | ☐ | ☐ | ☐ | ☐ | ☐ | ☐ | ☐ |
| Figure 6 | ☐ | ☐ | ☐ | ☐ | ☐ | ☐ | ☐ | ☐ | ☐ |
| Figure 7 | ☐ | ☐ | ☐ | ☐ | ☐ | ☐ | ☐ | ☐ | ☐ |
| Figure 8 | ☐ | ☐ | ☐ | ☐ | ☐ | ☐ | ☐ | ☐ | ☐ |
| Figure 9 | ☐ | ☐ | ☐ | ☐ | ☐ | ☐ | ☐ | ☐ | ☐ |
| Figure 10 or more (combine additional) | ☐ | ☐ | ☐ | ☐ | ☐ | ☐ | ☐ | ☐ | ☐ |

10. How many tables in the paper (not supplement)?

*Mark only one oval.*

| | 0 | 1 | 2 | 3 | 4 | 5 | |
|---|---|---|---|---|---|---|---|
| | ◯ | ◯ | ◯ | ◯ | ◯ | ◯ | Choose 5 for >=5 |

11. Does the change between preprint and paper in the main figures (including tables) reflect a change in content or outcomes? *

*Check all that apply.*

- [ ] No, no real changes at all (including reorganisation)
- [ ] No, panels or tables have just been moved around (including to supplement if available)
- [ ] Yes, significant additional content/outcomes have been added
- [ ] Yes, significant content/outcomes have been removed

12. Does the preprint have a supplement? How many files or items (figures, spreadsheets, extended methods)?

*Mark only one oval.*

| | 0 | 1 | 2 | 3 | 4 | 5 | |
|---|---|---|---|---|---|---|---|
| | ◯ | ◯ | ◯ | ◯ | ◯ | ◯ | Select option 5 for >=5 |

13. If the preprint does have a supplement, does it contain figures?

*Mark only one oval.*

◯ Yes

◯ No

14. Does the paper have a supplement? How many files or items (figures, spreadsheets, extended methods)?

*Mark only one oval.*

| | 0 | 1 | 2 | 3 | 4 | 5 | |
|---|---|---|---|---|---|---|---|
| | ◯ | ◯ | ◯ | ◯ | ◯ | ◯ | Select option 5 for >=5 |

15. If the paper does have a supplement, does it contain figures?

*Mark only one oval.*

- ( ) Yes
- ( ) No

16. Is the source data (including code) more accessible after publication?

*Check all that apply.*

- [ ] No, same as preprint - available only upon request
- [ ] No, same as the preprint - available through repositories or supplementary files
- [ ] Yes, provided as additional supplementary files
- [ ] Yes, provided through repositories

Other: [ ] ______________________

17. Are open peer reviews and decision letters available?

*Mark only one oval.*

- ( ) Yes, via Review Commons or any other post-publication review scheme
- ( ) Yes, provided by the journal
- ( ) No
- ( ) Don't know/ cannot locate this information

18. Any other comments?

______________________

______________________

______________________

______________________

______________________

This content is neither created nor endorsed by Google.

Google Forms
