## Supplemental Method 2. Instructions for annotation for "Preprints in motion: tracking changes between preprint posting and journal publication during a pandemic"

Make one highlight/annotation for each change (including replacements or changes, where text has been both removed and added). Highlight the word(s) most important to the change. Add an annotation in the comment below.

### Annotation rubric

You can enclose comments in parentheses.

Select one attribute from each section below and link together with underscores, for example:

results_nounchange_1-

results_removed_3

conclusion_effect-_2

#### Section

Context

Background or methods

results

A statement linked directly to data

conclusion

Interpretations and/or implications

#### Type of change

added

New assertion

removed

Assertion removed

nounchange

One noun is substituted for another (fever -> high fever)

effectreverse

The opposite assertion is now being made (word “negatively” added)

effect+

The effect is now stronger (changes in verbs/adjectives/adverbs)

effect-

The effect is now weaker (changes in verbs/adjectives/adverbs)

stat+

Statistical significance increased (expressed as number or in words)

stat-

Statistical significance decreased (expressed as number or in words)

statinfo

Addition/removal of statistical information (like a new test or confidence intervals)

#### Degree

1 (Significant): minorly alters a main conclusion of the paper

1- (Significant): **softens** a main conclusion of the paper

1+ (Significant): **strengthens** a main conclusion of the paper

2 (Major): a discrete change in a main conclusion of the paper

3 (Massive): a main conclusion of the paper reversed
