## Supplemental Method 3. Method notes for comparing Word files for "Preprints in motion: tracking changes between preprint posting and journal publication during a pandemic"

### Generating Word files

These new columns were added to our sheet listing preprint pairs.

| **line_break_and_label** | **line_break_and_fiducial** | **exclude** | **page_breaks** |
| --- | --- | --- | --- |

“Exclude” column was generated with this formula: =if(AND(R2="NA",Q2="Copied"),"keep","exclude")

It was intended to keep only rows NOT marked “1st paragraph” (where we copied the first paragraph if there is no abstract) and where we’ve noted that we want to exclude a paper because it has not yet been reviewed on F1000R or it is a duplicate.

Filtered for “keep” in this column.

Word version: 16.0.13001.20254

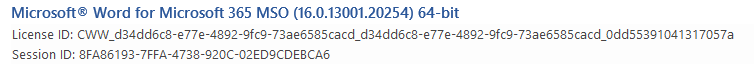

For preprint abstracts: Paste values of paper #, 2 line break columns, preprint DOI, preprint abstract, and page break column into Word document. Replace “??????” with “^m” to insert page breaks and “((((((“ with “^l” to insert line breaks

For published abstracts: Paste values of paper #, line break columns, paper DOI, **paper abstract**, and page break column into Word document. Replace “??????” with “^m” to insert page breaks and “((((((“ with “^l” to insert line breaks

Many published abstracts had lots of paragraph breaks. Removed (with replace) all paragraph breaks (^p) from both documents.

Settings for “compare documents”

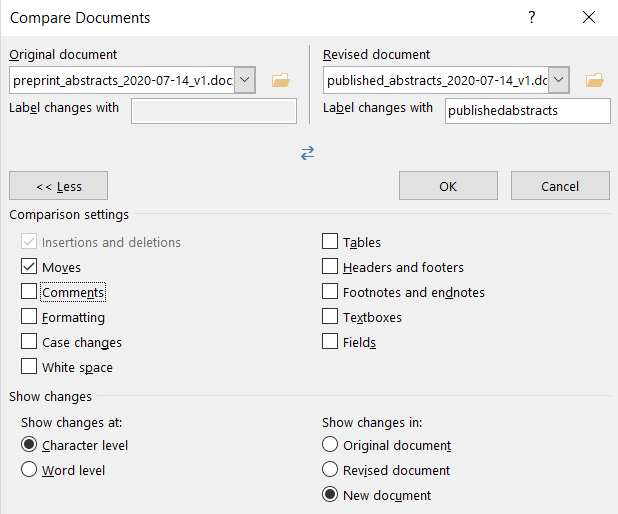

Also changed “advanced track changes options” to make insertions blue.

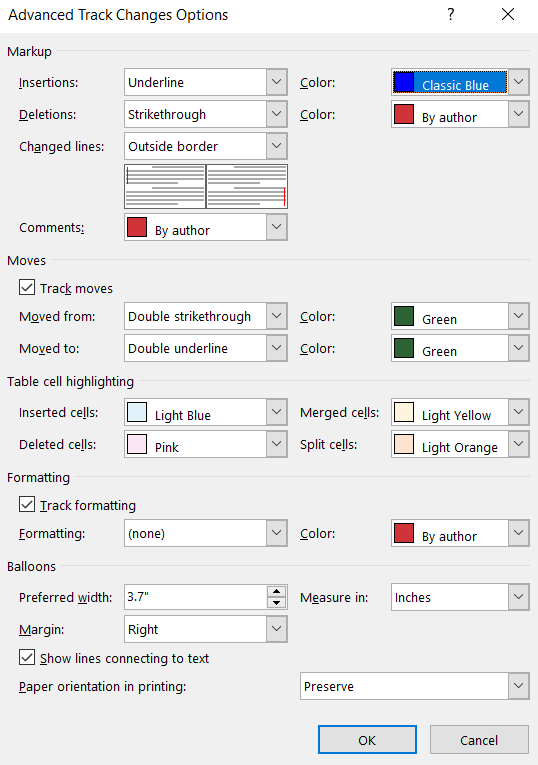

Since the alignment failed in several places originally (see 131 and 46 in 2020-07-13 file), added fiducial text to one of the line break columns in the spreadsheet. The two sentences (“We have manually inserted this sentence to assist with proper alignment of the text” and “These words will be deleted after the abstracts are properly combined.”) sometimes get broken up by the compare algorithm, but removing them separately (with replace, with track changes off) after comparing the documents seems to leave a clean file.

### Extracting markup

File/Print - then select “List of Markup” under settings.

Print to pdf.

Split pdf into 70 page documents, then opened with Google Docs and saved as .txt.

Final .txt file compiled from these pieces.

### Replacing abstracts with v1 of preprint version

*After we started our analysis, we noticed that we have used v2 preprint abstracts. The text below explains how we corrected this.*

For preprint abstracts: Paste values of paper #, 2 line break columns, preprint DOI, **version**, preprint abstract, and page break column into Word document. Replace “??????” with “^m” to insert page breaks and “((((((“ with “^l” to insert line breaks

Removed (with replace) all paragraph breaks (^p)

For published abstracts: Paste values of paper #, line break columns, preprint DOI, **version**, **paper abstract**, and page break column into Word document. Replace “??????” with “^m” to insert page breaks and “((((((“ with “^l” to insert line breaks

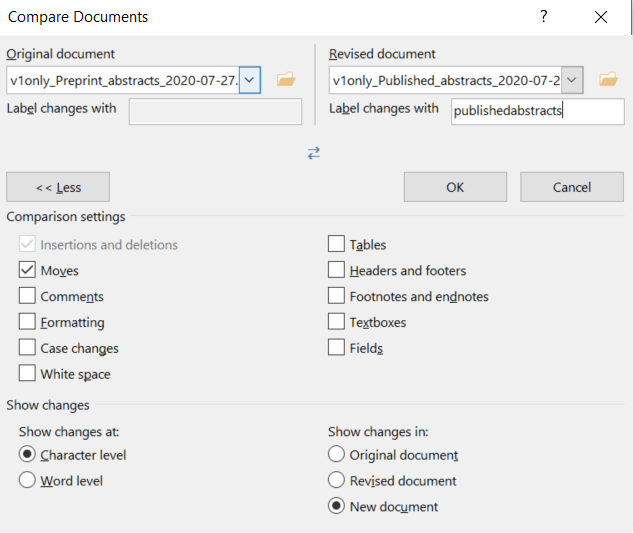

Remove fiducial sentences with replace AFTER the comparison. Print/save as .txt as above.
