## Supplemental Text 1. Key for journal abbreviations for "Preprints in motion: tracking changes between preprint posting and journal publication during a pandemic"

| **Journal title** | **Abbreviated label** |
| --- | --- |
| 3 Biotech | 3Btc |
| Antimicrobial Agents and Chemotherapy | AAaC |
| ACS Applied Bio Materials | AABM |
| Acta Neuropathologica Communications | AcNC |
| Acta Biomaterialia | ActB |
| American Journal of Clinical Pathology | AJoCP |
| American Heart Journal | AmHJ |
| Archives of Iranian Medicine | AoIM |
| Archives of Public Health | AoPH |
| Annals of the Rheumatic Diseases | AotRD |
| Acta Pharmaceutica Sinica B | APSB |
| Biochemical and Biophysical Research Communications | BaBRC |
| Brain, Behavior, and Immunity | BBaI |
| Biochemistry | Bchm |
| Biogerontology | Bgrn |
| Blood | Blod |
| BMC Genomics | BMCG |
| BMC Pediatrics | BMCP |
| BMC Endocrine Disorders | BMED |
| BMJ | BMJ |
| BMJ Open | BMJO |
| Bioinformatics | Bnfr |
| Brain | Bran |
| ClinicoEconomics and Outcomes Research | CaOR |
| Clinical Chemistry and Laboratory Medicine (CCLM) | CCaLM( |
| Cell | Cell |
| Clinical Gastroenterology and Hepatology | CGaH |
| Canadian Journal of Anesthesia/Journal canadien d'anesthésie | CJoAcd |
| Clinical Infectious Diseases | ClID |
| Cell Reports | CllRp |
| Cell Research | CllRs |
| Clinical Epigenetics | ClnE |
| Clinical Immunology | ClnI |
| Clinical Pharmacokinetics | ClnP |
| Clinical Microbiology and Infection | CMaI |
| Cancers | Cncr |
| Diversity and Distributions | DvaD |
| Developmental Biology | DvlB |
| EBioMedicine | EBMd |
| EClinicalMedicine | EClM |
| eLife | eLif |
| Emerging Microbes & Infections | EM&I |
| Emerging Infectious Diseases | EmID |
| Epidemics | Epdm |
| European Psychiatry | ErpP |
| European Respiratory Journal | ErRJ |
| Eurosurveillance | Ersr |
| ESMO Open | ESMO |
| EvoDevo | EvDv |
| Evolutionary Applications | EvlA |
| F1000Research | F100 |
| Frontiers in Integrative Neuroscience | FiIN |
| Frontiers in Plant Science | FiPS |
| Genetics in Medicine | GniM |
| Genetics | Gntc |
| Gastroenterology | Gstr |
| Heart Rhythm | HrtR |
| Influenza and Other Respiratory Viruses | IaORV |
| Infection, Genetics and Evolution | IGaE |
| International Journal for Parasitology | IJfP |
| International Journal of Antimicrobial Agents | IJoAA |
| IEEE Journal of Biomedical and Health Informatics | IJoBaHI |
| International Journal of Behavioral Nutrition and Physical Activity | IJoBNaPA |
| International Journal of Cancer | IJoC |
| International Journal of Environmental Research and Public Health | IJoERaPH |
| International Journal of Infectious Diseases | IJoID |
| Infectious Disease Modelling | InDM |
| Infection Prevention in Practice | IPiP |
| JMIR Medical Informatics | JMMI |
| Journal of Allergy and Clinical Immunology | JoAaCI |
| Journal of Clinical and Translational Science | JoCaTS |
| Journal of Clinical Pathology | JoCP |
| Journal of Hospital Infection | JoHI |
| Journal of Medical Internet Research | JoMIR |
| Journal of Medical Virology | JoMV |
| Journal of Neurology, Neurosurgery & Psychiatry | JoNN&P |
| Journal of Public Health | JoPH |
| Journal of the Intensive Care Society | JotICS |
| Journal of The Royal Society Interface | JoTRSI |
| Journal of Clinical Microbiology | JrnlofClnclMc |
| Journal of Clinical Medicine | JrnlofClnclMd |
| Journal of Infection | JroI |
| Journal of Neurology | JroN |
| Journal of Psychopharmacology | JroP |
| Journal of Virology | JroV |
| Molecular & Cellular Proteomics | M&CP |
| Mathematical Biosciences and Engineering | MBaE |
| mBio | mBio |
| Molecular Biology of the Cell | MBotC |
| Molecular Biology Reports | MlBR |
| Molecular Autism | MlcA |
| Multiple Sclerosis Journal | MlSJ |
| Mathematical Modelling of Natural Phenomena | MMoNP |
| mSphere | mSph |
| Metabolism | Mtbl |
| Neurology - Neuroimmunology Neuroinflammation | N-NN |
| Nature | Natr |
| Nucleic Acids Research | NcAR |
| New England Journal of Medicine | NEJoM |
| NeuroImage: Clinical | NI:C |
| Neurology | Nrlg |
| Neurobiology of Aging | NroA |
| Nature Machine Intelligence | NtMI |
| Nature Communications | NtrC |
| Nature Microbiology | NtrMc |
| Nature Medicine | NtrMd |
| Open Forum Infectious Diseases | OFID |
| Osong Public Health and Research Perspectives | OPHaRP |
| Public Health | PblH |
| Progress in Neuro-Psychopharmacology and Biological Psychiatry | PiNaBP |
| PLOS ONE | PLOO |
| Polar Biology | PlrB |
| PLOS Neglected Tropical Diseases | PNTD |
| Proceedings of the National Academy of Sciences | PotNAoS |
| Protein Science | PrtS |
| Psychological Medicine | PsyM |
| Psychiatry Research | PsyR |
| QJM: An International Journal of Medicine | QAIJoM |
| Quantitative Biology | QntB |
| Respiratory Research | RspR |
| Supportive Care in Cancer | SCiC |
| Science China Life Sciences | SCLS |
| Science | Scnc |
| Science of The Total Environment | SoTTE |
| Signal Transduction and Targeted Therapy | STaTT |
| Swiss Medical Weekly | SwMW |
| Therapeutic Advances in Neurological Disorders | TAiND |
| The American Journal of Emergency Medicine | TAJoEM |
| The British Journal of Psychiatry | TBJoP |
| The EMBO Journal | TEMJ |
| The Lancet Microbe | ThLM |
| The Journal of Infectious Diseases | TJoID |
| The Journal of Molecular Diagnostics | TJoMD |
| The Journal of Pathology: Clinical Research | TJoPCR |
| The Lancet Infectious Diseases | TLID |
| Translational Psychiatry | TrnP |
| Vaccine | Vccn |
| Virus Evolution | VrsE |
| Viruses | Vrss |
