## Supplementary tables 2 & 3 significant and representative changes for "Preprints in motion: tracking changes between preprint posting and journal publication during a pandemic"

### Supplementary table 2. Examples of changes in abstracts between the preprint and published version of an article

| **Example** | **Preprint DOI** | **Change (strikethrough: removed, bold: added)**  *Note that artifacts introduced by Word, such as duplicate numbers and symbols, have been removed.* | **Full annotation** |
| --- | --- | --- | --- |
| Added | 10.1101/2020.02.29.20029322 | To address this concern, the study employed a total of 214 general public ~~(GP)~~ and 526 nurses **(i.e., 234 front-line nurses and 292 non-front-line nurses)** to evaluate ~~VT~~ **vicarious traumatization** scores via a mobile app-based questionnaire. | Context_added_1 |
| Removed | 10.1101/19006478 | Multifocal **(MF)/multicentric (MC))** breast cancer is generally considered to be where two or more breast tumours are present within the same breast, ~~but are clearly separated with no intervening in situ or invasive disease.~~ | Context_removed_1 |
| Noun change | 10.1101/19009589 | We found that transportation potential was higher for source countries with seasonal dengue activity, high passenger traffic, high incidence rates, ~~lower economic status~~ **high epidemic vulnerability**, and in geographical proximity to a destination country in Europe. | Results_nounchange_1 |
| Effect + | 7  10.1101/2020.03.09.983247 | EK1C4 was also highly effective against membrane fusion and infection of other human coronavirus pseudoviruses tested, including SARS-CoV and MERS-CoV, as well as SARSr-CoVs, and potently ~~inhibiting~~ **inhibited the** replication of ~~4~~ **5** live human coronaviruses examined, including SARS-CoV-2. | Results_effect+_1 |
| Effect - | 10.1101/2020.01.23.916395 | The mean estimate of R0 for the 2019-nCoV ranges from ~~3.30 (95%CI: 2.73-3.96) to 5.47 (95%CI: 4.16-7.10)~~,**2.24 to 3.58** , and **is** significantly larger than 1 | Results_effect-_1 |
| Stat+ | 10.1101/19010975 | Logistic regression AUCs were 0.~~82~~**83**, 0.~~82~~**83**, 0.~~79~~**81** for three different regularization schemes; tree boosting AUC was 0.81; MLP AUC was 0.~~81~~**83** | Results_stat+_1+ |
| Stat+ (wording) | 10.1101/19006031 | ~~These findings reveal a modest negative association .:~~ **Findings describe small but robust associations** between lifetime MDD and **lower** cognitive performance. within a population-based sample. | Conclusions_stat+_1+ |
| Stat- | 10.1101/19011064 | Greater paternal engagement (OR 1.~~59~~**5** (1.~~13~~09, 2.~~21~~**07**) was associated with [...] | Results_stat-_1- |
| Stat- (wording) | 10.1101/19005710 | Elevated STE from 5- to 19 year-olds indicates that school-aged children were **likely** the most important transmitters of infection during the autumn wave of the 2009 pandemic in the US**A.** | Conclusion_stat-_1- |
| Statinfo | 10.1101/19007013 | The ~~MR~~ **Mendelian randomisation** analysis and **single** ~~SNP~~-**nucleotide polymorphism** analysis, however, did not support this**. (odds ratio for lifetime smoking on suicidal ideation, 0.050; 95% CI -0.027 to 0.127; odds ratio on suicide attempts, 0.053; 95% CI, -0.003 to 0.110)**. | Results_statinfo_1+ |

### Supplementary Table 3. All changes in abstracts that resulted in a major conclusion change

| **Preprint DOI** | **Change (strikethrough: removed, bold: added)**  *Note that artifacts introduced by Word, such as duplicate numbers and symbols, have been removed.* | **Annotation with significant changes**  *Note that other annotations appearing in the same passage are not shown.* |
| --- | --- | --- |
| 10.1101/19006833 | ~~Conclusion~~ Reproductive risk factor distributions are different from European populations but exhibited etiologic heterogeneity by age at diagnosis and ER status similar to other populations. **Differences in reproductive patterns and subtype heterogeneity are consistent with racial disparities in subtype distributions.** | conclusions_added_2 (reconciled, premise of study but racial disparities not noted elsewhere) |
| 10.1101/19008367 | We conclude that ML algorithms combined with EMR capture early life ASD risk~~.~~ **as well as reveal previously unknown features to be associated with ASD-risk**. Such approaches may be able to enhance the ability for accurate and efficient early detection of ASD in large populations of children. | Conclusions_added_2 |
| 10.1101/19012823 | Conclusion: The study suggests ~~the~~ **that factors such as age, educational achievement, and** access to ~~mobile technology is the predominant determinant~~ **internet are significant predictors** of ~~utilization~~ **acceptability of a mHealth application among cancer patients**. | Conclusions_added_2 *(duplicate annotation also present)* |
| 10.1101/2020.01.03.20016436 | Children ~~from richer households, urban regions, primary maternal education and male~~ **in Addis Ababa and Dire Dawa were seven times more likely to have full vaccination compared to children living in the Afar region.** | results_added_2 |
| 10.1101/2020.01.26.20018754 | The me~~di~~an time from illness onset to ~~hospitalization~~ **hospital admission (for treatment and/or isolation)** was estimated at 3**–4 days without truncation and at 5–9 days when right truncated**. | Results_added_2 |
| Ibid. | Based on the ~~estimate of the~~ 95th percentile estimate of the incubation period, we recommend that the length of ~~isolation and~~ quarantine should be at least ~~nine~~ **14** days. | Conclusion_effect+_2 (recommendation length changed substantially) (reconciled – change could be very impactful) |
| 10.1101/2020.01.31.20019901 | **We** estimated that the median **daily** reproduction number~~, R, fluctuated between 1.6-2.9 from mid-December to mid-January 2020. We found that the US, Australia and France had more confirmed cases with travel history to Wuhan than the model predicted, and estimated that there were 29,500 (14,300-85,700) prevalent symptomatic cases in Wuhan on 23rd January 2020, when~~ **(Rt) in Wuhan declined from 2·35 (95% CI 1·15–4·77) 1 week before** travel restrictions were introduced~~.~~ | Results_nounchange_2 (fluctuated -> declined) |
| Ibid. | Based on our estimates of ~~R~~**Rt, assuming SARS-like variation**, we calculated [...] | **context_added_2** |
| Ibid. | **Our** results show that ~~2019-nCoV has substantial potential for ongoing human-to-human~~ **COVID-19** transmission~~, and exported cases from~~ **probably declined in** Wuhan ~~may have increased prior to travel restrictions being introduced on 23rd~~ **during late** January, 2020~~.~~**, coinciding with the introduction of travel control measures**.  Results_removed_1  Results_added_1  Results_removed_1 (increase before travel restriction)  Conclusions_nounchange_2 (from concluding cases increased until travel restrictions to concluding that they decreased after travel restrictions) | Conclusions_nounchange_2 (from concluding cases increased until travel restrictions to concluding that they decreased after travel restrictions) |
| 10.1101/2020.02.06.20020974 | **During the first 2 months of the current outbreak, Covid-19 spread rapidly throughout China and caused varying degrees of illness**. | Conclusions_added_2 |
|  | **Patients often presented** without **fever, and many did not** have abnormal [1] radiological findings | Conclusions_added_2 |
|  | ~~Severe pneumonia was independently associated with either the admission to intensive care unit, mechanical ventilation, or death in multivariate competing-risk model (sub-distribution hazards ratio, 9.80; 95% confidence interval, 4.06 to 23.67).~~ | Results_removed_2 |
|  | ~~The disease severity (including oxygen saturation, respiratory rate, blood leukocyte/lymphocyte count and chest X-ray/CT manifestations) predict poor clinical outcomes .~~ | Conclusions_removed_2 (revised abstract does not talk about predicting outcomes) |
| 10.1101/2020.02.11.20020735 | **Regardless of different universal facemask wearing policy scenarios, facemask shortage would occur but eventually end during our prediction period (from 20 Jan 2020 to 30 Jun 2020).** | Results_effectreverse_2 (preprint abstract makes no mention of end of mask shortage) |
|  | **The duration of the facemask shortage described in the scenarios of a country-wide universal facemask wearing policy, a universal facemask wearing policy in the epicentre, and no universal facemask wearing policy were 132, seven, and four days, respectively .** | results_added_2 |
|  | **During the prediction period, the largest daily facemask shortages were predicted to be 589·5, 49·3, and 37·5 million in each of the three scenarios, respectively .** | Results_added_2 |
|  | **In any scenario, an N95 mask shortage was predicted to occur on 24 January 2020 with a daily facemask shortage of 2·2 million** | Context_added_2 |
|  | **Implementing a universal facemask wearing policy in the whole of China could lead to severe facemask shortage.** | Conclusions_added_2 |
|  | **Without effective public communication, a universal facemask wearing policy could result in societal panic and subsequently, increase the nationwide and worldwide demand for facemasks .** | Conclusions_added_2 |
|  | **To fight novel infectious disease outbreaks, such as COVID-19, governments should monitor domestic facemask supplies and give priority to healthcare workers.** | Conclusions_added_2 |
|  | **The risk of asymptomatic transmission and facemask shortages should be carefully evaluated before introducing a universal facemask wearing policy in high-risk regions.** | Conclusions_added_2 |
|  | **Public health measures aimed at improving hand hygiene and effective public communication should be considered along with the facemask policy.** | Conclusions_added_2 |
| 10.1101/2020.02.11.20022186 | **Thus, in a scenario where we have taken twenty times the confirmed number of infected and forty times the confirmed number of recovered cases, the case fatality ratio is around ∼0.15% in the total population.** | Conclusions_added_2 |
|  | **Importantly, based on this scenario, simulations suggest a slow down of the outbreak in Hubei at the end of February.** | Conclusions_added_2 |
| 10.1101/2020.02.16.20023671 | N8R **and NLR** may serve as a useful prognostic factor for early identification of severe COVID-19 cases. | Conclusions_added_2 |
| 10.1101/2020.02.19.956581 | We determined ~~a~~ cryo-~~electron microscopy structure~~ **EM structures** of the SARS-CoV-2 S ectodomain trimer, ~~demonstrating spontaneous opening of the receptor-binding domain, and~~ providing a blueprint for the design of vaccines and inhibitors of viral entry. | Result_removed_2 (reconciled) |
| 10.1101/2020.02.27.20028829 | **In addition, the intrinsic growth rate was estimated at 0.6 (95% CI: 0.6, 0.7), and the scaling of growth parameter was estimated at 0.8 (95% CI: 0.7, 0.8),, indicating sub-exponential growth dynamics of COVID-19.** | Results_added_2 (reconciled, sex difference added) |
|  | **The crude case fatality rate is higher among males (**1~~.6), which indicates~~**1%) compared to females (0.4%) and increases with older age.** | Results_added_2+ |
| 10.1101/2020.02.27.967760 | **And thus, both** the vertical transmission **and the placenta dysfunction/abortion caused by SARS-CoV-2 need to be further carefully investigated in clinical practice.** | Conclusions_added_2 (reconciled, potential for dysfunction/abortion added) |
| 10.1101/2020.02.29.20029322 | **The** results showed that the ~~VT scores slightly increased across periods of aiding COVID-19 control, although no statistical difference was noted (P = 0.083). ). However, the study found lower scores for VT in nurses [median = 69; interquartile range (IQR) = 56-85] than those of the GP (median = 75.5; IQR = 62-88.3) (P = 0.017). In addition, the VT~~ **vicarious traumatization** scores for front-line nurses ~~(FLNs; median = 64; IQR = 52-75),~~ including scores for physiological and psychological responses, were significantly lower than those of non-front-line nurses ~~(nFLNs; median = 75.5; IQR = 63-92)~~ (P < 0.001)/ | Results_removed_2 (reconciled, nurses split into two groups, extensive changes in statistics) |
| 10.1101/2020.03.08.20032946 | **According to these estimates, presymptomatic transmissions alone are almost sufficient to sustain epidemic growth.** | Conclusions_added_2 |
|  | **The use of a contact-tracing app that builds a memory of proximity contacts and immediately notifies contacts of positive cases would be sufficient to stop the epidemic if used by enough people, in particular when combined with other measures such as physical distancing.** | Conclusion_added_2 |
|  | **An intervention of this kind raises ethical questions regarding access, transparency, the protection and use of personal data, and the sharing of knowledge with other countries. Careful oversight by an inclusive advisory body is required.** | Conclusion_added_2 |
| 10.1101/2020.03.09.984856 | After a broad screening of additional viruses, we now describe in vitro activity against Chikungunya ~~virus (CHIK) and Middle Eastern Respiratory Syndrome Coronavirus (MERS-CoV).~~ | Conclusion_removed_2 |
| 10.1101/2020.03.13.20035485 | ~~Even higher reduction should take place in France, Germany~~ **the incubation period** and ~~Spain for which β > 1.3 (as of March 10, 2020).~~ | Conclusions_removed_2 (no longer prescribing actions to other countries) |
|  | ~~Considering longer-term dynamics in the case when the initial exponential growth is not contained by quarantine, we analyzed a~~ **the serial interval reported by epidemiologists. Using the model, we consider a hypothetical** scenario in which ~~rE~~**ß is modulated solely by anticipated changes of social behaviours: first, ß** decreases in response to ~~the~~ **a** surge of daily new cases ~~which convinces~~**, pressuring** people to **self-**isolate ~~themselves.~~ | Results_removed_2 |
|  | [...] although economically and socially devastating, grants time to develop~~, produce,~~ and ~~distribute~~ **deploy** vaccine **or at least limit daily cases to a manageable number.** | Conclusion_added_2 |
| 10.1101/2020.03.19.20038950 | **Based on our predictions of Iran about 29000 people will be infected from March 25 to April 15, 2020.** | Results_added_2 |
|  | On average, **12**92~~5~~ people with COVID-19 are expected to be infected daily in Iran. | Results_effect+_2 |
|  | The epidemic peaks within one week (15.03.20203 days (March 25 to 03.21.March 27, 2020) and reaches its highest point on 03.18.March 25, 2020 with 171265 infected cases | Results_effect+_2 |
| 10.1101/2020.03.24.20041020 | **Hence, we do not recommend any of these reported prediction models to be used in current practice.** | conclusion_added_2 |
|  | **Finally, studies should adhere to the TRIPOD (transparent reporting of a multivariable prediction model for individual prognosis or diagnosis) reporting guideline.** | Conclusion_added_2 |
| 10.1101/773812 | ~~We propose that Spire stimulates nucleation by Cappuccino in a manner similar to the collaboration between APC and mDia1.~~ | Conclusions_removed_2 (above it does say that “pointed ends are retained” but it’s not as clear that Spire is directly binding those) *(duplicate annotation also present)* |
| 10.1101/785691 | **However, over** long-term **laboratory** evolution ~~experiment. However, as~~ **spanning 60 000 generations,** | Results_effect+_2 |
|  | **mutation frequency in TFs gradually declined after a rapid initial burst.** | Results_added_2 (much longer evolution 60000 vs 10000 totally changed the outcome??) |
| 10.1101/852541 | These ~~patterns were consistent across nine islands spanning more than 1000 kilometers of the archipelago~~ | Results_removed_3 |
|  | **effects of deforestation on ecosystem productivity, however, were highly variable between years and among the study islands**. | Results_added_3 |
